# Competing constraints shape the non-equilibrium limits of cellular decision making

**DOI:** 10.1101/2022.07.01.498451

**Authors:** Nicholas C. Lammers, Avi I. Flamholz, Hernan G. Garcia

## Abstract

Gene regulation is central to cellular function. Yet, despite decades of work, we lack quantitative models that can predict how transcriptional control emerges from molecular interactions at the gene locus. Thermodynamic models of transcription, which assume that gene circuits operate at equilibrium, have previously been employed with considerable success in the context of bacterial systems. However, the presence of ATP-dependent processes within the eukaryotic transcriptional cycle suggests that equilibrium models may be insufficient to capture how eukaryotic gene circuits sense and respond to input transcription factor concentrations. Here, we employ simple kinetic models of transcription to investigate how energy dissipation within the transcriptional cycle impacts the rate at which genes transmit information and drive cellular decisions. We find that biologically plausible levels of energy input can lead to significant gains in how rapidly gene loci transmit information, but discover that the regulatory mechanisms underlying these gains change depending on the level of interference from non-cognate activator binding. When interference is low, information is maximized by harnessing energy to push the sensitivity of the transcriptional response to input transcription factors beyond its equilibrium limits. Conversely, when interference is high, conditions favor genes that harness energy to increase transcriptional specificity by proofreading activator identity. Our analysis further reveals that equilibrium gene regulatory mechanisms break down as transcriptional interference increases, suggesting that energy dissipation may be indispensable in systems where non-cognate factor interference is sufficiently large.

## Introduction

Throughout biology, systems must make accurate decisions under time constraints using noisy molecular machinery. Eukaryotic gene regulation exemplifies this challenge: genes must read out input concentrations of transcription factor proteins and respond by producing appropriate levels of gene product (mRNA and eventually protein) in order to drive downstream cellular decisions. Interestingly, the gene activity underlying cellular decision-making is often subject to large amounts of noise. Indeed, experiments across a wide range of organisms have revealed that eukaryotic transcription is highly stochastic, occurring in episodic bursts (Bothma et al., 2014; Tantale et al., 2016; Nicolas et al., 2017; Lionnet and Wu, 2021)—periods of activity interspersed with periods of transcriptional silence—that unfold over timescales ranging from minutes to hours (Lammers et al., 2020). Because of this stochasticity, the transcription rate is a noisy reflection of transcription factor concentration. Over time, the accumulation of gene products tends to average out this noise, but biological processes must operate under time constraints: cells in developing fruit fly embryos have only minutes to determine their developmental fates (et al. Alberts B, Johnson A, Lewis J, 2002; Desponds et al., 2020), antigen recognition in T-cells unfolds over a single day (Obst, 2015), and cells in adult tissues are constrained by mRNA half-lives that range from minutes to days (Pérez-Ortín et al., 2013).

A key question, therefore, is how the molecular architecture of gene loci—the number and identity of biochemical steps in the transcriptional cycle and the reaction rates connecting these steps—dictates the amount of time needed for bursty gene expression to drive accurate cellular decisions. In particular, while it is widely accepted that processes within the eukaryotic transcriptional cycle consume biochemical energy (Coulon et al., 2013; Wong and Gunawardena, 2020), we do not yet know what non-equilibrium should “look like” in the context of transcriptional systems. Indeed, it remains challenging not only to predict unambiguous signatures of energy expenditure that can be detected experimentally (Hammar et al., 2014; Park et al., 2019; Eck et al., 2020), but also to establish how energy consumption can be harnessed to improve gene regulatory performance in the first place (Zoller et al., 2021).

Here, we use concepts from information theory and statistical physics as a lens to investigate how energy dissipation impacts the timescale on which gene circuits can drive cellular decisions. We consider a simple binary choice scenario wherein a cell must decide, as rapidly as possible, whether it is subjected to a high (*c*_1_) or low (*c*_0_) concentration of a transcriptional activator based on the transcriptional output of a gene locus. The basis for this decision is the gene’s input-output function (Figure 1A and B), which emerges from microscopic interactions between input activator molecules and their target gene loci (Figure 1C) that induce differences in the output dynamics of transcriptional bursting (Figure 1D) for high and low activator concentrations. In turn, these differences in burst dynamics drive different rates of mRNA accumulation (Figure 1E). Because each ON/OFF fluctuation is stochastic, the resulting gene expression levels are noisy, and the cell must wait some time *T* before it is possible to accurately distinguish between *c*_1_ and *c*_0_. Our central question in this work is whether energy dissipation within the molecular processes driving transcription allows gene loci to decrease the decision time, *T*, and, if so, how this performance gain manifests in terms of measurable features of the transcriptional input-output function.

**Figure 1.**
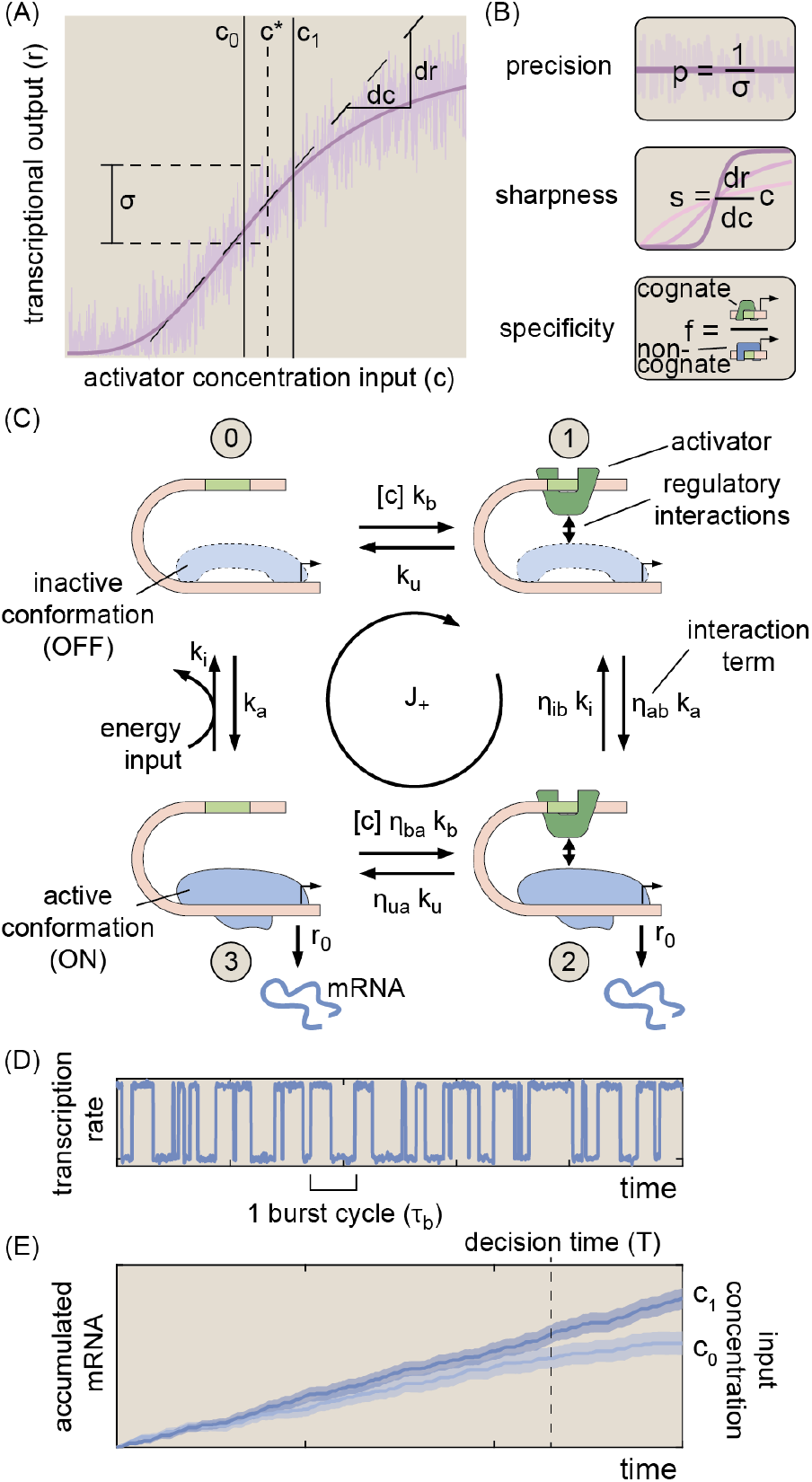
Three regulatory features shaping transcriptional information transmission. **(A)** Gene regulatory input-output function illustrating the basic biological problem considered in this work. Here, a cell must distinguish between two activator concentrations, *c*_0_ and *c*_1_, based on the transcriptional output of a gene locus (purple curve). **(B)** We examine how three regulatory features of the transcriptional input-output function—sharpness, precision, and specificity—combine to dictate the rate at which the transcriptional output drives biological decisions. **(C)** Four-state MWC-like model (Phillips and Orme, 2020) of transcription used as the foundation of our investigations. Here, a single activator (green square) may bind to a specific site at the gene locus, and mRNA production occurs when the gene locus switches to its active (ON) conformation. A hypothetical energy input is depicted along the rate from state 3 to state 0. In practice, our framework permits non-equilibrium driving to occur along any of the eight transition rates in the model. **(D)** Simulated burst dynamics for one realization of the model shown in (C). Activator binding drives different burst dynamics at loci exposed to high and low activator concentrations. The burst cycle time is defined as the average time required to complete one ON → OFF → ON cycle and sets the timescale over which biological decisions unfold. **(E)** Illustrative simulation results for accumulated mRNA levels driven by *c*_1_ and *c*_0_. Solid lines show trajectories for a single locus, and shaded regions indicate the standard deviation of levels taken across 100 simulated trajectories. The vertical dashed line indicates the “decision time,” when the expected mRNA levels driven by *c*_1_ and *c*_0_ are sufficiently different to permit an accurate decision about the input activator concentration.

There are multiple ways in which energy dissipation could alter the input-output behavior of a gene locus to improve cellular decision-making. As illustrated in Figure 1A and B, non-equilibrium processes could increase sensitivity to differences in input transcription factor concentration (“sharpness”) or suppress transcriptional noise (“precision”). Since our model assumes that, in addition to the concentration of the cognate activator, *C*, the gene locus is subject to some level of non-cognate factors, *W*, energy dissipation could also buffer against interference from off-target activation (“specificity”).

Recent works have begun to uncover a complex space of tradeoffs among these three aspects of transcriptional performance both at and away from thermodynamic equilibrium. A recent study found that systems operating at thermodynamic equilibrium suffer from strict tradeoffs between transcriptional specificity and transcriptional precision, but this tradeoff can be overcome by gene circuits that spend energy to enhance specificity through a scheme reminiscent of classical kinetic proofreading (Shelansky and Boeger, 2020; Ninio, 1975; Hopfield, 1974). Likewise, a separate study demonstrated that energy dissipation can enhance transcriptional sharpness (Estrada et al., 2016). Interestingly, while energy can increase sharpness and specificity *separately*, another study found that nonequilibrium levels of specificity come at the cost of sub-optimal sharpness (Grah et al., 2020). The authors also found that energy dissipation tends to *decrease* transcriptional precision, although this conclusion likely hinges on the study’s modeling assumptions (Grah et al., 2020). Despite this progress, it remains unclear how these non-equilibrium gains and tradeoffs ultimately impact how effectively gene circuits can harness differences in transcription factor concentrations to drive cellular decisions.

In this work, we identify a key quantity, the rate of information transmission from input transcription factor concentrations to output transcription rates as the quantitative link between energy-dependent changes in the transcriptional input-output function (Figure 1B) and the speed at which gene loci drive accurate biological decisions (Figure 1E) (Siggia and Vergassola, 2013; Desponds et al., 2020). We use this rate as a lens to examine the interplay between energy dissipation and cellular decision-making. We consider model gene circuits with varying numbers of activator binding sites. We also examine models with different numbers of molecular steps in the activation pathway, since transcriptional activation is also thought to require multiple molecular steps beyond activator binding itself, such as the localization of key transcription factors to the gene locus (Nogales et al., 2017).

We demonstrate that energy dissipation increases the rate at which genes can drive cellular decisions for all models considered. Moreover, the presence of multiple activation steps enables gene loci to more effectively harness energy to increase information transmission. At the level of the transcriptional input-output function (Figure 1A), while energy input can drive increases in all three regulatory features considered (sharpness, precision, and specificity; Figure 1B), genes cannot realize these non-equilibrium gains simultaneously. In particular, we show that the upper limit of information transmission is defined by a shifting tradeoff between sharpness and specificity. When the relative concentration of wrong-to-right activator species is small (e.g., in the fruit fly embryo), non-equilibrium gene circuits that maximize sharpness drive the fastest decisions. However, when the ratio of non-cognate to cognate activator concentrations is larger than the intrinsic difference in their binding affinities (e.g., in mammalian cells), gene circuits must instead prioritize transcriptional specificity.

In closing, we identify hallmarks of non-equilibrium gene regulation that may be amenable to experimental detection. We use our model to illustrate how simple point mutations in activator binding sites can lead to robust signatures of non-equilibrium regulatory processes. Additionally, our findings emphasize the importance of using theoretical models that account for noncognate factor binding when interpreting experimental measurements of gene expression. Altogether, this work provides a rigorous foundation for interrogating the role of energy dissipation in eukaryotic gene circuit regulation.

## Results

### A. A simple model for probing the interplay between energy and information in transcription

We sought to establish gene circuit models that capture two essential characteristics of eukaryotic transcription. First, gene regulation hinges upon interactions between specific and general transcription factors. Although salient regulatory information tends to reside exclusively in a few specific transcription factors targeted to binding sites within enhancers (Vincent et al., 2016), these proteins are not sufficient to give rise to transcription. Instead, transcription and transcriptional control depend on interactions between specific regulatory factors and other key molecular players at the gene locus, such as mediators (Grah et al., 2020; Malik and Roeder, 2016; Rybakova et al., 2015; Kagey et al., 2010), RNA polymerase (Tantale et al., 2016), nucleosomes (Shelansky and Boeger, 2020; Mirny, 2010), and various sub-units of the pre-initiation complex (Nogales et al., 2017). While these factors do not themselves carry biological information, they constitute key molecular steps within the transcriptional cycle. This multiplicity of molecular players implies that gene loci may exist in multiple distinct molecular states corresponding to different binding configurations of specific and general molecules (e.g., (Biddle et al., 2019)). Moreover, some of these processes—e.g., nucleosome displacement (Zhou et al., 2016), pre-initiation complex assembly (Taatjes, 2017), and RNA polymerase initiation (Yan and Gralla, 1997)—entail the dissipation of biochemical energy, opening the door to non-equilibrium behaviors.

Second, it has recently become apparent that eukaryotic transcription is characterized by stochastic, episodic bursts of activity interspersed with periods of transcriptional silence (Bothma et al., 2014; Fukaya et al., 2016; Little et al., 2013; Zoller et al., 2018; Tantale et al., 2016; Lammers et al., 2020). Since the concentration of specific transcription factors can regulate burst dynamics (Lammers et al., 2020; Zoller et al., 2018; Xu et al., 2015), a simple model would suggest that transcriptional bursts originate from the binding and unbinding of specific transcription factors. Although this may be the case in some yeast genes (Donovan et al., 2019), recent *in vivo* measurements in higher eukaryotes have revealed that activators and repressors typically bind DNA for seconds, rather than minutes or hours (Lammers et al., 2020; Lionnet and Wu, 2021). This temporal disconnect between bursting and transcription factor binding suggests a model in which transcriptional burst cycles—corresponding to OFF → ON → OFF fluctuations in the locus conformation (Figure 1D)—are not determined by transcription factor binding alone, but entail additional molecular reactions that are decoupled from the timescale of activator binding.

Together, these observations support a Monod– Wyman–Changeux (MWC)-like framework (Phillips and Orme, 2020; Grah et al., 2020; Shelansky and Boeger, 2020; Mirny, 2010) for modeling transcription wherein specific transcription factors act as effector molecules, conditioning the frequency with which the gene locus fluctuates between active and inactive transcriptional conformations. The simplest model that meets this description is one where a transcriptional activator binds to a single binding site at the gene locus, and where a second molecular reaction dictates fluctuations between two conformations: an inactive (OFF) state where no mRNA is produced and a transcriptionally active (ON) state where mRNA is produced at rate *r*_0_.

If we neglect the binding of non-cognate transcription factors, this leads to the model shown in Figure 1C. This model contains four basal reaction rates: the transcription factor binding and unbinding rates (*k*_b_ and *k*_u_) and the locus activation and deactivation rates (*k*_a_ and *k*_i_). We leave the molecular identity of this locus activation step unspecified, but in principle, it may be any of the elements of the general transcriptional machinery mentioned above. In addition to these basal rates, the *η* terms in Figure 1C capture interactions between the transcription factor and activation step. Here, the first subscript indicates which molecular reaction the *η* term modifies (binding or unbinding; activation or inactivation), and the second subscript indicates the molecule performing the modification (bound activator “b” or activated molecular step “a”). For instance, *η*_ab_ encodes the degree to which the rate of locus activation is modified by having a transcription factor bound at the locus (*η*_ab_ *>* 1 corresponds to an activating transcription factor). Lastly, the average rate of mRNA production in this model is simply equal to 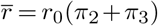, where *π*_*i*_ is the steady-state probability of finding the system in state *i*.

### B. Calculating energy dissipation rates and decision times

At equilibrium, all state transitions in our model must obey the law of microscopic reversibility. Energy dissipation along one or more of the microscopic transitions shown in Figure 1C lifts this strict equilibrium constraint and opens the door to novel forms of non-equilibrium gene regulatory logic. For the model shown in Figure 1C, the energy dissipated per unit time (Φ) can be expressed as

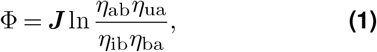

where the *η* terms are defined in Figure 1C and the net cycle flux, ***J***, encodes the degree to which microscopic transitions in the system are biased in the clockwise (***J*** *>* 0) or counterclockwise (***J*** *<* 0) direction (Hill, 1989). See Appendix A.5 for further details. Φ is a strictly positive quantity with units of k_B_T per unit time that indicates how “near” or “far” a system is from thermodynamic equilibrium (Hill, 1989; Lang et al., 2014). For ease of comparison across different realizations of our model gene circuit, we express Φ in units of k_B_T per burst cycle (“energy per burst”).

Our central aim is to understand how energy dissipation impacts the rate at which gene loci transmit information and drive cellular decisions. For simplicity, we assume that *c*_0_ and *c*_1_ are constant over time. We also stipulate that the difference between these concentrations (*δc*) is relatively small, such that *δc* = *c*_1_ − *c*_0_ = 0.1*c*^∗^, where *c*^∗^ is the midpoint concentration *c*^∗^ = (*c*_1_ + *c*_0_)*/*2. This value of *δc* is equivalent, for example, to concentration differences for the activator Bicoid between adjacent nuclei in early fruit fly development (Gregor et al., 2007). Figure 1E shows trends indicating the predicted integrated transcriptional output of a gene locus when it is exposed to high or low activator concentrations. Intuitively, it should be easier to distinguish between these two scenarios when (i) the difference between average transcript production rates (slope of the lines in Figure 1E) is large or (ii) the noise (shaded regions) in the accumulated output is small.

IR codifies this intuition, providing a quantitative measure of a gene’s ability to read out and respond to different input activator concentrations. Formally, IR is defined as the rate of change in the Kullback–Leibler divergence (Cover and Thomas, 2006) between our two hypotheses (*C* = *c*_0_ and *C* = *c*_1_) given the expected transcriptional output of our model gene circuit. If we take the noise in the transcriptional output to be approximately Gaussian (see Appendix B), IR can be expressed as

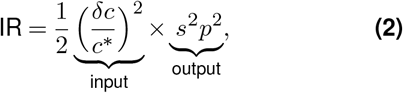

where IR is strictly positive and has units of information per unit time and *s* and *p* are the sharpness and precision of the transcriptional response, respectively, as defined in Figure 1B. See Appendix C for a full derivation of this expression. We note that the native units of Equation 2 are natural log units (“nats”). For simplicity, we give all informational quantities in the more familiar “bits,” such that IR has units of bits per burst cycle (“bits per burst”). Additionally, the precision term, *p*, pertains solely to noise from intrinsic fluctuations between microscopic states at the gene locus and does not account for Poisson noise resulting from mRNA synthesis. In general, this noise is expected to be small relative to the noise from locus fluctuations for the parameter regimes considered (see Appendix D for details).

Equation 2 contains two terms: an input component that encodes the size of the activator concentration gradient and an output component that depends on the sharpness and precision of the transcriptional input-output function (Figure 1A and B). This expression provides quantitative support for the intuitions outlined above. IR can be increased both by increasing the difference between the transcription rates driven by *c*_1_ and *c*_0_ (i.e., increasing the sharpness) and by decreasing the noise level (i.e., increasing precision). Moreover, since both *s* and *p* can be calculated analytically from the microscopic reaction rates in our gene circuit (see Appendix A), Equation 2 allows us to calculate and compare information rates for gene circuits with different microscopic reaction rates.

The IR, in turn, dictates how rapidly cells can distinguish between the two activator concentrations, *c*_0_ and *c*_1_, based on the accumulated transcriptional output of a gene circuit. Previous works (Siggia and Vergassola, 2013; Desponds et al., 2020) have established that the theoretical lower limit for the time required to distinguish between *c*_0_ and *c*_1_ is given by

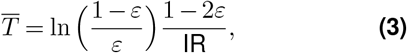

where *ε* is the probability of being wrong, i.e., choosing *c*_1_ when the true value is *c*_0_ (or vice versa) (see Appendix E and (Desponds et al., 2020) for details). We note the error-tolerance *ε* in Equation 3 is extrinsic to the gene circuit model and depends on the nature of the downstream cellular processes. Unless otherwise noted, we follow (Desponds et al., 2020) and set *ε* = 0.32, equivalent to an error level of “1 sigma.”

### C. Energy dissipation increases the rate of information transmission

Utilizing our framework, we investigated whether increasing the energy dissipated by our model gene circuit, Φ, increases the rate at which this circuit drives cellular decisions between *c*_0_ and *c*_1_. We expanded methods employed in (Estrada et al., 2016; Eck et al., 2020) to develop an algorithm capable of systematically exploring how different transition rates dictate gene circuit features. This algorithm can determine the maximum IR achievable by different realizations of our gene circuit as a function of energy dissipation. See Appendices F and G for details regarding its implementation and validation.

Figure 2A shows the relation between IR and Φ resulting from our numerical analysis. Here, each circle represents IR and Φ values for a single realization of our gene circuit (Figure 1C), as defined by its complement of transition rate values. Near equilibrium, our analysis reveals that gene circuits can transmit information no faster than 0.035 bits per burst (far left-hand side of Figure 2A). According to Equation 3, this means that the best equilibrium gene circuits require at least 110 burst cycles to drive a decision between concentrations *c*_1_ and *c*_0_ with an error probability of 32% when these concentrations differ by 10% (Figure 2B). In the developing fruit fly embryo (*D. melanogaster*), where the burst timescale (*τ*_*b*_) is approximately 2 minutes (Lammers et al., 2020), this translates to a decision time of 3.7 hours, far too long to meet the time constraints imposed by early nuclear cleavage cycles (8–60 minutes (et al. Alberts B, Johnson A, Lewis J, 2002)). Our equilibrium gene circuit would require even longer times in adult nematode (*C. elegans*) and mouse (*M. musculus*) cells, where *τ*_*b*_ is much higher, with measurements ranging from 61 to 105 minutes (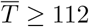 hours, (Lee et al., 2019)) and 30 minutes to multiple hours (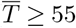 hours, (Lammers et al., 2020)), respectively. In each case, these timescales likely exceed decision time limits imposed by mRNA decay or cellular division times, which set upper limits on the time over which gene output can be averaged (horizontal lines in Figure 2B; see Appendix H for further details).

**Figure 2.**
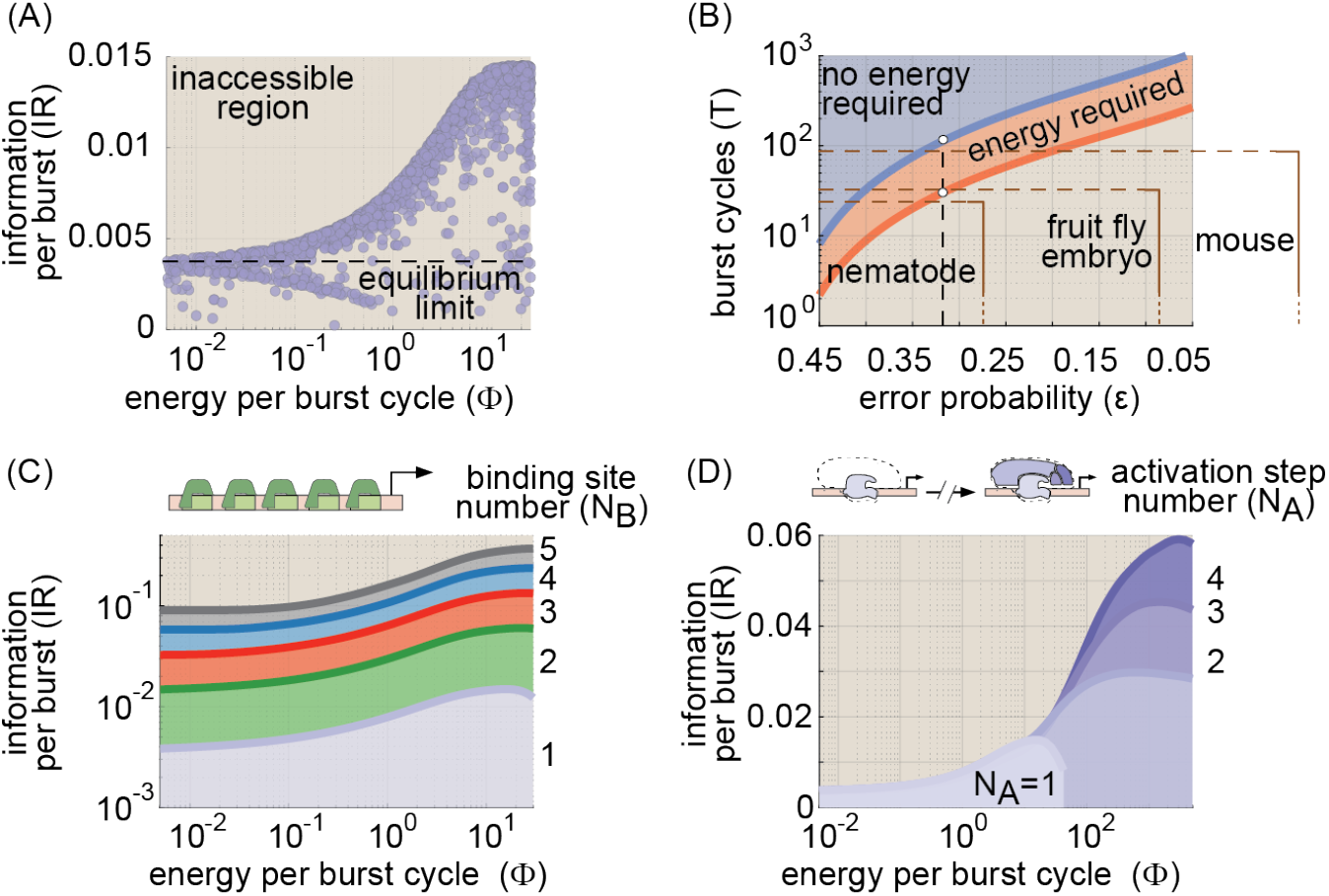
Energy dissipation increases the information transmission rate in gene circuits. **(A)** Information rate (IR from Equation 2) as a function of energy dissipation rate (Φ from Equation 1) for a parameter sweep exploring all possible model realizations. Modest energy dissipation rates can lead to a significant increase in the maximum amount of information that can be transmitted per burst cycle. **(B)** The amount of time needed to distinguish between *c*_0_ and *c*_1_ as a function of the probability of deciding incorrectly for equilibrium and non-equilibrium gene circuits. The decision time is given in terms of the number of transcriptional burst cycles required for a decision to be made. Note that the x-axis is arranged in order of *decreasing* error probability (i.e., increasing accuracy) from left to right. Horizontal lines indicate approximate upper bounds on decision times (in burst cycles) for different biological systems. **(C)** Parameter sweep results for achievable IR and Φ values for gene circuits with 1–5 activator binding sites. Achievable regimes for each molecular architecture are indicated as color-coded shaded regions. **(D)** Sweep results illustrating achievable IR vs. Φ regimes for gene circuits featuring 2–5 locus conformations. (For all parameter sweep results in A-D, transition rate and interaction term magnitudes, *k* and *η*, were constrained such that 10^*−*5^ ≤ *kτ*_*b*_ ≤ 10^5^ and 10^*−*5^ ≤ *η* ≤ 10^5^, where *τ*_*b*_ is the burst cycle time. *η*_*ab*_ and *η*_*ib*_ were further constrained such that *η*_*ab*_ ≥ 1 and *η*_*ib*_ ≤ 1, consistent with our assumption that the transcription factor activates the gene locus.)

Our analysis indicates that energy dissipation opens the door to improved information transmission, leading to a fourfold increase in the upper IR limit from 0.0035 to 0.014 bits per burst cycle (Figure 2A). Moreover, this performance gain is realized at biologically plausible levels of energy consumption: IR reaches its maximum non-equilibrium value at Φ ≈ 20 k_B_T per cycle, which is approximately equivalent to the hydrolysis of one to two ATP molecules (Milo and Phillips, 2015). This corresponds to an energy-dependent decrease in decision time from 110 to 29 burst cycles (red shaded region in Figure 2B). This reduction meets the upper decision limit for mouse cells (Figure 2B). Yet there remains an absolute speed limit that no amount of energy dissipation can overcome, as shown by the empty space below the red non-equilibrium boundary in Figure 2B.

How can gene circuits do better? Real transcriptional systems are typically far more complex than the simple four-state model in Figure 1C; gene enhancers typically feature multiple transcription factor binding sites (Vincent et al., 2016), and transcriptional activation depends on the combined action of multiple molecular components at the gene locus (Lammers et al., 2020). Thus, to overcome this speed limit, we must examine the impact of tuning two molecular “knobs”: the number of specific activator binding sites in our model (N_B_) and the number of molecular steps required to achieve productive transcription (N_A_). For simplicity, we focus on systems in which all binding sites are identical and assume identical kinetics for all molecular transitions between locus conformations. While restrictive, this simple approach gives rise to rich, biologically salient behaviors. While we explore the effects of varying N_B_ and N_A_ separately, these mechanisms are mutually compatible and may act jointly in real biological systems. See Appendix I for details regarding the implementation of these higher-order models.

#### Adding binding sites improves information-energy tradeoffs

We first examined the performance of gene circuit models with multiple binding sites. In these models (as with the four-state model described above), activator binding does not directly dictate transitions into and out of transcriptionally active molecular states, but instead increases or decreases the likelihood of these transitions. Models with multiple binding sites also permit cooperative interactions between activator molecules, encoded by *η*_*ub*_ terms (see Appendix I and Figure A9A). With these assumptions, we employed our parameter sweep algorithm to explore tradeoffs between the rate of energy dissipation (Φ) and the IR for systems with 1–5 activator binding sites. In all cases, we held the number of activation steps constant at N_A_ = 1 (as in Figure 1C).

As illustrated in Figure 2C, adding activator binding sites shifts the IR vs. Φ tradeoff boundary from Figure 2A upwards, allowing for higher information transmission rates for a given energy dissipation rate. This leads to significant IR gains, even in gene circuits operating near the equilibrium limit (vertical dashed line in Figure 2C), with the upper equilibrium limit increasing by approximately a factor of 25 from 0.0035 bits per burst cycle for N_B_ = 1 to 0.090 bits per cycle for N_B_ = 5. As a result, equilibrium gene circuits with 5 binding sites need as little as 5 burst cycles to distinguish between *c*_1_ and *c*_0_, easily satisfying the decision time constraints of the biological systems shown in Figure 2B (Figure S1A). More generally, the lower decision time limit scales as the inverse of the number of binding sites squared (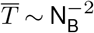, see Figure S1A).

#### Adding molecular activation steps allows gene circuits to harness higher rates of energy dissipation

Next, we expanded the four-state model by changing the number of activation steps (1 ≤ N_A_ ≤ 4) while holding the number of binding sites fixed at N_B_ = 1 (top panel of Figure 2D). To illustrate this model, let us first consider the baseline case, where N_A_ = 1. Here, locus activation depends on the state of a single molecular component (e.g., mediator), which can be disengaged (i.e., the locus is OFF) or engaged (i.e., the locus is ON). Now, consider model in which locus activation also depends on the state of a second molecular component (e.g. PIC assembly) that can, likewise, be either engaged or disengaged. If we stipulate that both components must be engaged to achieve RNA polymerase initiation, then two molecular activation steps are required to reach the ON state and N_A_ = 2. We use the same logic to extend the model to the N_A_ = 3 and N_A_ = 4 cases to capture the impact of the additional molecular components necessary for transcription. See Appendix I and Figure A9B for details.

We conducted parameter sweeps to examine the interplay between energy dissipation and information transmission for these systems. As with adding binding sites, the addition of activation steps leads to increased rates of information transmission. Unlike increasing N_B_, however, these IR gains do not come for free. Instead, the addition of activation steps extends the Φ-IR boundary into higher-energy regimes, allowing non-equilibrium gene circuits to achieve larger gains in IR at the expense of increased energy dissipation rates (Figure 2D).

This increased IR gain means that systems with multiple activation steps can drive decisions between *c*_1_ and *c*_0_ more rapidly than the simple four-state gene circuit. For example, non-equilibrium gene circuits with four activation steps can drive decisions nearly four times as rapidly as systems with a single step (8 vs. 29 burst cycles; see Figure S1B). This 8-burst-cycle limit approaches what can be achieved by an equilibrium gene circuit with 5 activator binding sites (5 burst cycles; compare Figure S1A and B), suggesting a similarity between adding activator binding sites at equilibrium and adding activation steps out of equilibrium. However, this parity has an energetic cost: to approach the performance of the five-binding-site model, the one-binding-site system with five conformations must dissipate at least 180 k_B_T per burst.

### D. Increases in non-equilibrium sharpness improve information transmission

According to Equation 2, the energy-dependent increases in IR uncovered in Figure 2 must result from increased sharpness, increased precision, or some combination thereof. Thus, to uncover how energy reshapes the transcriptional input-output function to increase IR, we used our numerical sweep algorithm to examine the space of achievable sharpness and precision values for our baseline four-state model (Figure 1C) both at and away from thermodynamic equilibrium. One challenge in comparing sharpness and precision levels across different gene circuits is that the upper bounds on both *s* and *p* depend on the fraction of time, *π*_a_, the system spends in the transcriptionally active conformation, which changes as the transition rates vary between different realizations of our gene circuit. Thus, for ease of comparison across different model realizations, we give all results in terms of normalized sharpness and precision measures: S = *s/b* and P = *pb*, where *b* = *π*_a_(1 − *π*_a_) is the binomial variance in the occupancy of the transcriptionally active conformation. These metrics have intuitive interpretations: the S value of a particular gene circuit’s input-output function gives the Hill coefficient of an equivalently sharp Hill function, and P is inversely proportional to the level of intrinsic noise in the transcriptional output. See Appendix J for details.

Figure 3A shows the results of our analysis, with each circle representing the S and P values for a single gene circuit realization. For systems operating at equilibrium (blue dots in Figure 3A), we find that both S and P are bounded by “Hopfield barriers” (dashed lines) (Hopfield, 1974; Estrada et al., 2016) with values of 1 and 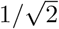, respectively. These bounds place strict limits on information transmission at equilibrium and have a straight-forward interpretation: they are precisely equal to the sharpness and precision of a simple two-state gene circuit with a single activator binding site and no molecular activation step, where the ON rate is concentration-dependent (*k*_on_ ∝ [*c*], see Appendix K for details).

**Figure 3.**
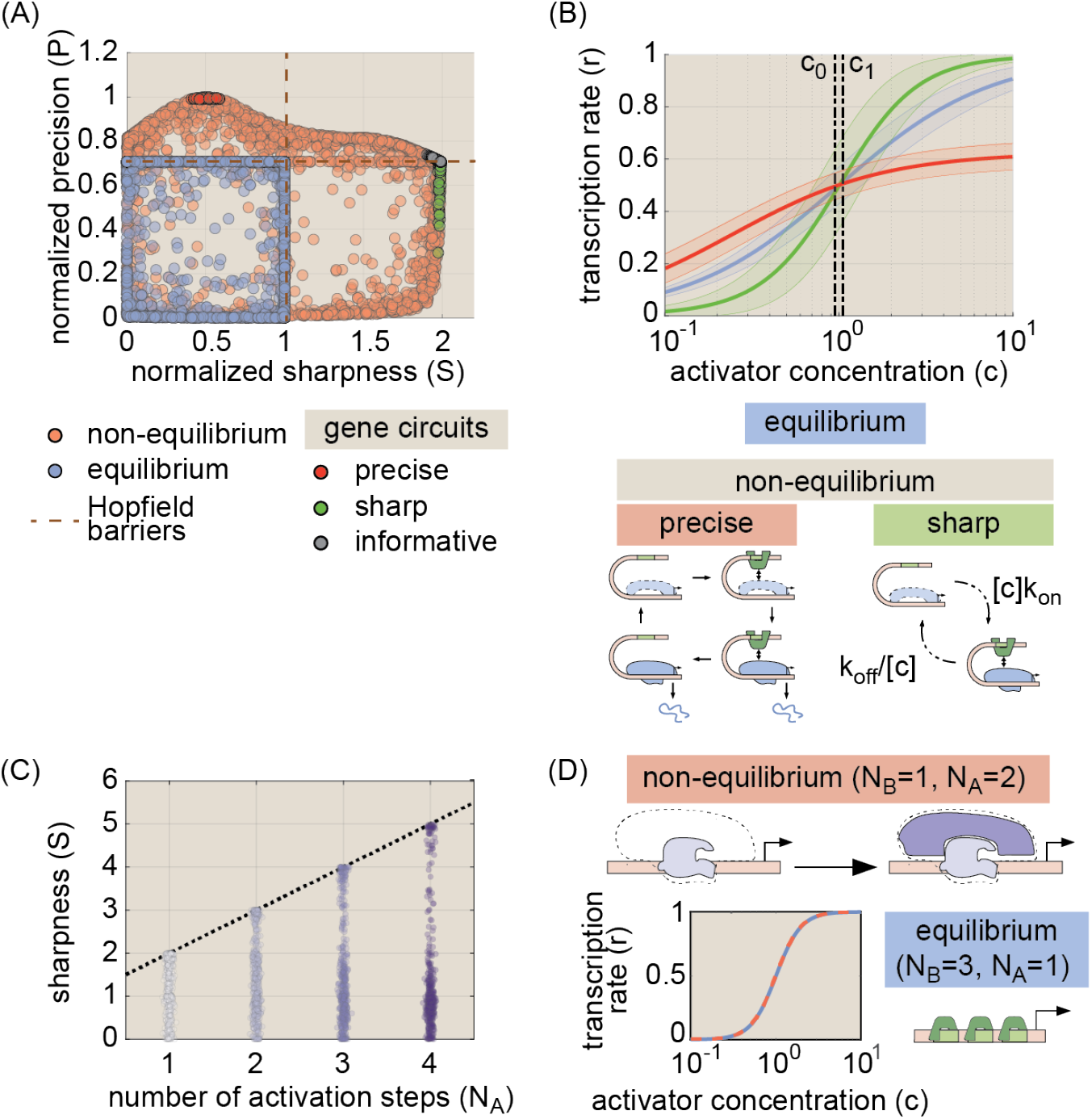
Increased transcriptional sharpness drives increased information transmission away from equilibrium. **(A)** Scatter plot of parameter sweep results showing the normalized sharpness and precision of 3,000 simulated gene circuits with and without equilibrium constraints. Energy expenditure overcomes Hopfield-like barriers, doubling the upper sharpness limit and increasing the precision limit by a factor of 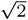. The absence of gene circuits in the upper right quadrant indicates that no circuits can simultaneously maximize sharpness and precision. Calculations indicate that IR-maximizing systems (gray circles) spend energy to maximize non-equilibrium sharpness while maintaining precision at the maximum equilibrium level. **(B)** Illustrative input-output functions for a maximally informative equilibrium gene circuit (blue) from the parameter sweeps shown in (A) and maximally sharp and precise non-equilibrium gene circuits (green and red, respectively). The shaded region indicates predicted noise levels in gene expression patterns after 25 bursting cycles. Cartoons below illustrate molecular motifs for maximally precise and sharp non-equilibrium gene circuits. **(C)** Plot of achievable non-equilibrium sharpness levels for models with 2–5 locus conformations and one activator binding site. Each circle represents a single gene circuit model. Normalized sharpness is bounded by the number of locus conformations. **(D)** Cartoon illustrating functional equivalence between three binding sites at equilibrium and two activation steps out of equilibrium. The plot shows input-output functions for maximally sharp realizations of each case, demonstrating the equivalent sharpness levels driven by the two strategies. (For parameter sweep results in A and C, transition rate and interaction term magnitudes, *k* and *η*, were constrained such that 10^*−*5^ ≤ *kτ*_*b*_ ≤ 10^5^ and 10^*−*5^ ≤ *η* ≤ 10^5^, where *τ*_*b*_ is the burst cycle time. *η*_*ab*_ and *η*_*ib*_ were further constrained such that *η*_*ab*_ ≥ 1 and *η*_*ib*_ ≤ 1, consistent with our assumption that the transcription factor activates the gene locus.)

Energy dissipation permits gene circuits to overcome these equilibrium performance bounds, increasing S by up to a factor of 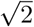 and P by up to a factor of 2 with respect to their equilibrium limits (Figure 3A). Yet, while energy can improve sharpness and precision individually, the absence of realizable gene circuits in the upper-right-hand corner of Figure 3A indicates that genes cannot maximize both simultaneously. This tradeoff places inexorable limits on the degree to which energy can boost IR and—as illustrated in Figure 3B— arises because maximally sharp and maximally precise gene circuits require distinct and incompatible underlying molecular architectures (see Appendix L for details).

Because sharpness and precision cannot be maximized simultaneously, gene circuits that dissipate energy must “choose” which aspect to maximize. From the perspective of IR maximization, the choice is clear: Figure 3A shows the location of 100 gene circuits within 1% of the maximum of 0.014 bits per cycle (Figure 2A) in S − P phase space (gray circles). Thus, the most informative gene circuits maximize transcriptional sharpness (S = 2) at thecost of retaining equilibrium precision levels 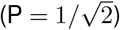, which makes sense given that non-equilibrium systems can boost S by up to a factor of 2 while P is limited to a maximum gain of 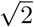. As with the equilibrium case, these S and P values have an intuitive interpretation: they are simply equal to the expected sharpness and precision of a two-state system, one in which both the ON and OFF rates are concentration-dependent (see Appendix M). Thus, although spending energy to overcome the constraints of detailed balance opens up a vast new space of possible regulatory schemes, maximally informative non-equilibrium gene circuits exhibit an emergent simplicity, converging upon architectures in which their many molecular degrees of freedom collapse into a few effective parameters that define system behavior.

#### Non-equilibrium gains in sharpness drive IR increases in more complex regulatory architectures

To assess the generality of our results, we used our parameter sweep algorithm to examine equilibrium and non-equilibrium tradeoffs between sharpness and precision for more complex gene circuits with 2–5 activator binding sites and 2–4 molecular activation steps. In all cases, energy dissipation increases the upper limits of S and P, and as with our simple four-state model, these non-equilibrium performance gains cannot be realized simultaneously (Figure S2A and B). For all models considered, the gains in IR uncovered in Figure 2 are maximized by spending energy to increase sharpness, rather than precision (see Appendix N for further details). For the case of multiple activator binding sites (N_B_ *>* 1), the N_B_-dependent increases in IR shown in Figure 2C arise because increasing the number of binding sites increases the upper sharpness limit both at and away from equilibrium (Figure S2A-C and Appendix N; (Grah et al., 2020; Estrada et al., 2016)).

More surprisingly, we find that increasing the number of molecular conformations (N_A_) while holding the number of activator binding sites can increase transcriptional sharpness in systems operating out of equilibrium. Figure 3C shows the range of achievable S values for non-equilibrium systems as a function of N_A_. The upper S limit scales linearly with N_A_, such that S_neq_ ≤ N_A_ + 1. This linear scaling is identical to the effect of adding activator binding sites at equilibrium, where S_eq_ ≤ N_B_ (Figure S2C), providing intuition for why systems with multiple molecular steps can drive faster decisions: with respect to transcriptional sharpness, the regulation of multiple activation steps by a single binding site in a non-equilibrium gene circuit is functionally equivalent to the effect of having multiple binding sites at equilibrium (Figure 3D).

### E. Energy dissipation is required for rapid cellular decisions at high non-cognate factor concentrations

In real biological settings, cells do not contain only a single species of transcription factor. Therefore, to drive timely biological decisions, a gene circuit must not only sense and respond to its cognate transcription factor, but also efficiently filter out “irrelevant” signals from noncognate factors. This process is inherently challenging in eukaryotes, where short DNA-binding footprints lead to modest energetic differences between specific (correct) and non-specific (incorrect) transcription factor binding events on the order of 4.6 k_B_T (Maerkl and Quake, 2007), meaning that non-cognate transcription factors unbind from gene loci approximately 100-fold faster than cognate factors 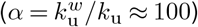.

To understand whether this 100-fold difference in binding kinetics is sufficient to drive decisions in real biological systems, we examined a stripped-down scenario in which cognate and non-cognate activators must compete to bind a single binding site (Figure 4A). We can quantify the severity of non-cognate factor interference by dividing the fraction of time the site is bound by a cognate factor (*π*_*c*_) by the total fraction of time it is bound by *either* the cognate or non-cognate species (*π*_*c*_ + *π*_*w*_). If we assume equal basal binding rates (*k*_b_) for cognate and non-cognate species, then the fraction of time the locus spends bound by a cognate transcription factor is given by

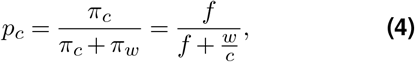

where we introduce a new quantity, the transcriptional specificity (*f*), defined as the (average) ratio of the probability of having cognate and non-cognate factors bound, normalized by the concentration, namely

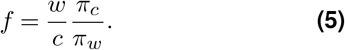

**Figure 4.**
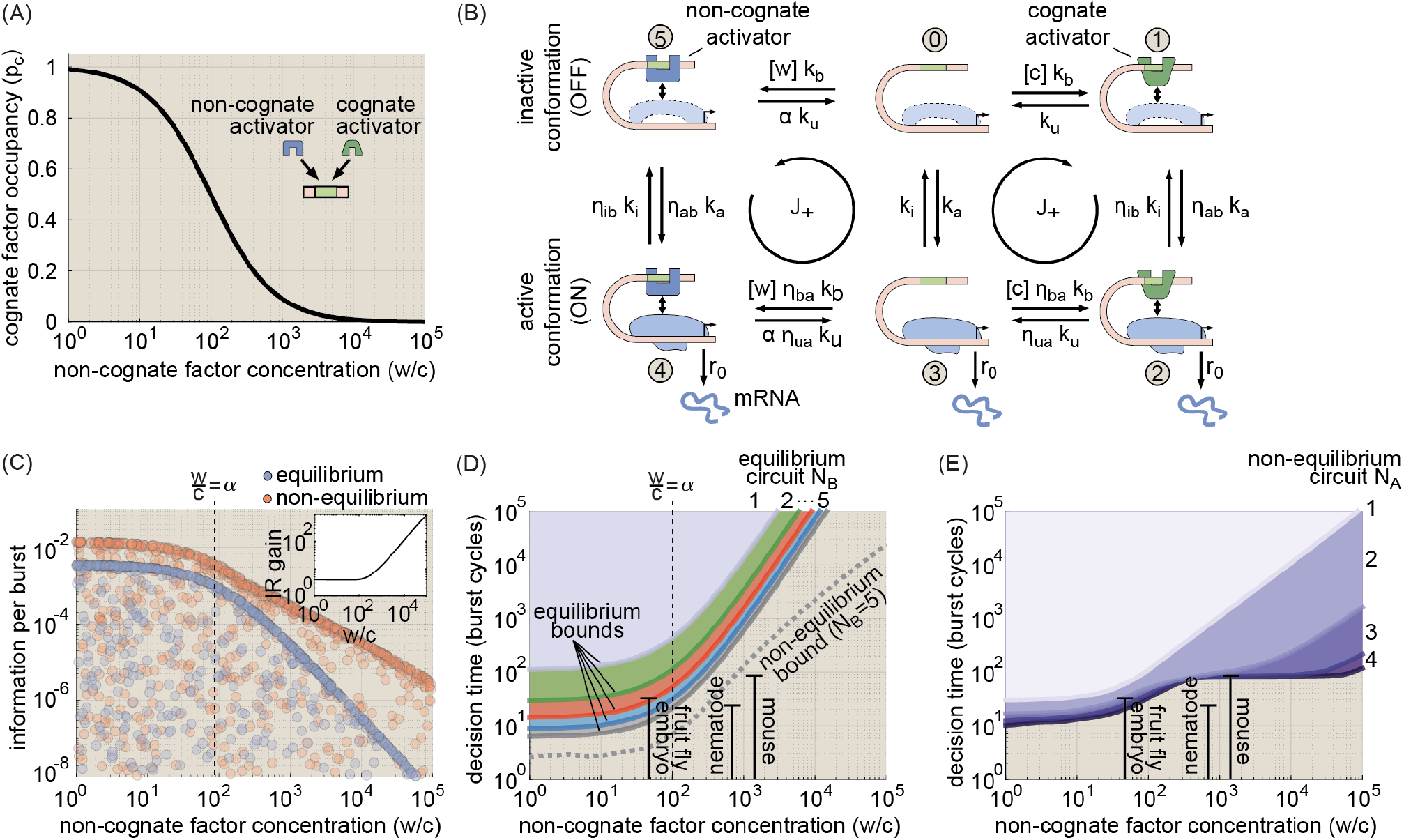
Energy dissipation is key to driving cellular decisions in the presence of non-cognate factor interference. **(A)** Cognate factor occupancy at a single binding site as a function of relative non-cognate factor concentration. **(B)** Incorporating non-cognate activator binding leads to a six-state model that features both a right and a wrong activation pathway. **(C)** Numerical results for the maximum achievable information rate for equilibrium (blue circles) and non-equilibrium (red circles) gene circuits with one activator binding site and one activation step (illustrated in (B)) as a function of the relative concentration of non-cognate activators *w/c*. The blue dashed line indicates the upper IR bound at equilibrium. The red line indicates the predicted non-equilibrium IR bound assuming quadratic scaling with w/c (see main text). The vertical dashed line indicates where the non-cognate factor concentration (*w*) equals the cognate factor concentration multiplied by the affinity factor (*αc*). Note how the optimal non-equilibrium systems begin to exceed the predicted bound beyond this point. The inset panel shows the non-equilibrium performance gain as a function of *w/c*. **(D)** Shaded regions indicate parameter sweep results for the range of achievable decision times for equilibrium gene circuits with 1–5 activator binding sites as a function of *w/c*. The dashed gray line indicates the lower bound for decision times driven by *non-equilibrium* gene circuits with five binding sites and one activation step. See Figure S3B for corresponding information rate ranges. **(E)** Decision times for non-equilibrium gene circuits with 1–4 activation steps. See Figure S3C for corresponding information rate ranges. (All results assume 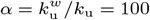. All decision time quantities assume *ε* = 0.32. For parameter sweep results in C-E, transition rate and interaction term magnitudes, *k* and *η*, were constrained such that 10^*−*5^ ≤ *kτ*_*b*_ ≤ 10^5^ and 10^*−*5^ ≤ *η* ≤ 10^5^, where *τ*_*b*_ is the burst cycle time. *η*_*ab*_ and *η*_*ib*_ were further constrained such that *η*_*ab*_ ≥ 1 and *η*_*ib*_ ≤ 1, consistent with our assumption that the transcription factor activates the gene locus.)

We note that Equation 5, which considers competition between two activator species to bind and activate a single gene, is distinct from and complements specificity definitions employed in previous works, which examine the problem for a single activator species that regulates a cognate and a non-cognate locus (Shelansky and Boeger, 2020; Grah et al., 2020) (see Appendix O.1 for details).

From Equation 4, we see that *f* sets the scale for the severity of non-cognate factor interference. At equilibrium, *f* is equal to the affinity factor *α* (see Appendix O.2), such that cognate factor binding dominates when *w/c < α* and non-cognate factors dominate when *w/c* exceeds *α*. For concreteness, we set *α* = 100 throughout the remainder of this work. Where do actual biological systems fall? A recent study pursuing synthetic enhancer design in the early fly embryo cited 47 pertinent regulatory factors that were controlled to avoid off-target binding (Vincent et al., 2016), leading to an estimate of *w/c* = 47 (see also (Estrada et al., 2016)). Inserting this value into Equation 4, we predict that the cognate factor will be bound approximately 2/3 of the time in the fly embryo. At the other end of the spectrum, we can use the genomic abundance of transcription factor proteins to estimate upper bounds on *w/c* values for adult nematode and mouse cells, yielding estimates of *w/c* ≤ 698 and *w/c* ≤ 1, 426, respectively (Charoen-sawan et al., 2010). In this case, Equation 4 predicts that cognate binding accounts for only a small fraction of total binding interactions—as little as 1/8 in worms and 1/15 in mice—suggesting that equilibrium affinity differences alone may be insufficient in these cases. To examine how these high interference levels impact the timescale of biological decisions and to determine whether energy dissipation can improve upon this equilibrium baseline, we must extend our gene circuit model to incorporate interference from non-cognate activator binding.

To do this, we draw inspiration from (Cepeda-Humerez et al., 2015), adding a second “wrong” activation cycle to our original four-state model (Figure 1C), wherein the binding of a non-cognate factor to the gene locus can also induce transitions to the active conformation. This leads to the six-state model shown in Figure 4B, where, for simplicity, we have grouped all non-cognate activators into a single concentration term: *W*. Here, states 5 and 4 are identical to states 1 and 2, except that a non-cognate activator species (blue circle) is bound rather than the cognate activator (green square). For notational convenience, we write the unbinding rates of the non-cognate activator 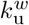 as the unbinding rate of the cognate factor *k*_u_ multiplied by an affinity factor 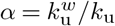, with *α* = 100.

We employed parameter sweeps to examine the upper limits on information transmission as a function of the ratio of wrong-to-right activator concentrations (*w/c*). We held the cognate factor concentration at *C* = *c*^∗^, such that *W* was the only variable concentration parameter. Figure 4C presents the range of achievable information rates as a function of the relative wrong factor concentration. Our results reveal that the rate of information transmission at equilibrium drops precipitously once *w/c* exceeds *α*(blue circles in Figure 4C). Away from equilibrium, the upper information limit likewise decreases with *w/c*; however, we find that non-equilibrium gene circuits are significantly more robust to high non-cognate factor concentrations than equilibrium systems. The relative IR gain from energy dissipation increases from a factor of 4 when *w/c* ≈ 1 to a factor of 1,000 when *w/c* = 10^5^ (Figure 4C, inset). This shift in information gain suggests that a qualitative change occurs in how energy is used once *w/c > α* (vertical dashed line) (see Section F).

We next used Equation 3 to calculate the amount of time required for a cell to decide between concentrations *c*_0_ and *c*_1_ of the cognate activator species for different values of *w/c*, starting with gene circuits constrained to operate at equilibrium. As in Figure 2B, we compared our model’s performance to the decision time limits for different biological systems, this time with each organism placed appropriately along the *w/c* axis. In all organisms considered, gene circuits generally have a few tens of burst cycles over which to transmit information, with no organism exceeding 100 bursts (black error bars in Figure 4D). This decision time limit is significantly shorter than can be achieved by our simple six-state model with one binding site and one activation step at equilibrium, even in the presence of negligible amounts of non-cognate transcription factor (*w/c* = 1, purple shaded region corresponding to N_B_ = 1 in Figure 4D).

Next, we investigated the effect of having equilibrium gene circuits with multiple sites. Figure 4D indicates that equilibrium gene circuits with three or more activator binding sites (red, blue, and gray regions) are sufficient to drive timely decisions in “low-interference” systems such as the early fruit fly embryo. However, we again observe a precipitous decline in performance once *w/c > α*. Indeed, the best equilibrium model (N_B_ = 5) can drive decisions in no fewer than 1,100 burst cycles—the equivalent of at least 550 hours (3 weeks) for mouse cells—when *w/c* ≈ 1, 400 (the upper limit for mice). This finding is over an order of magnitude too slow for the mouse system’s decision time limit of 86 burst cycles (Figure 4D). Moreover, our analysis suggests that at least 17 activator binding sites are needed at equilibrium (see Figure S3A). Such a number is conceivable for eukaryotic enhancers, but this analysis emphasizes that equilibrium systems—even those with biologically salient numbers of binding sites—struggle to achieve realistic decision times in the presence of significant non-cognate factor interference.

How do non-equilibrium gene circuits fare? The dashed gray line in Figure 4D indicates the lower decision time limit for *non-equilibrium* gene circuits with five binding sites and one activation step. We observe a substantial improvement relative to the equilibrium case; however performance nonetheless suffers at large values of *w/c*, falling short of the decision time limit for the mouse system (209 vs. 86 burst cycles). We used our parameter sweep algorithm to examine the impact of increasing the number of molecular activation steps (N_A_ *>* 1) in non-equilibrium gene circuits with a single activator binding site. This revealed substantial improvements, particularly at large *w/c* values. Whereas the N_A_ = 1 system required at least 1,500 burst cycles when *w/c* = 1, 400, gene circuits with two activation steps can drive decisions between *c*_0_ and *c*_1_ in as little as 104 bursts (Figure 4E), a full order of magnitude over equilibrium genes with five binding sites and twice that of non-equilibrium gene circuits with five binding sites and a single activation step (Figure 4D). Adding a third step further improves this bound to 83 burst cycles, below the 86-burst limit for the mouse system. Moreover, this N_A_ = 3 system exhibits remarkable robustness to non-cognate factor interference, sustaining the same level of performance up to *w/c* ≈ 10^4^ (Figure 4E).

These results suggest that, in biological contexts where the ratio of wrong-to-right activator concentrations exceeds the intrinsic binding affinity difference (*α*), energy dissipation increasingly becomes a necessary precondition for driving cellular decisions within biologically salient timescales. Moreover, the presence of multiple molecular activation steps greatly amplifies non-equilibrium performance gains in these high-interference regimes. Yet Figure 4E also reveals that one-binding-site systems have a performance limit. To further improve, non-equilibrium gene circuits likely require multiple molecular steps (N_A_ ≥ 2) *and* multiple activator binding sites (N_B_ ≥ 2).

### F. Non-cognate factor concentration defines performance tradeoffs between sharpness and specificity

Next, we investigated how much sharpness and precision each contribute to the IR gain depicted in the panel inset of Figure 4C. Figure 5A shows the relative non-equilibrium gains in S and P (S*/*S^eq^ and P*/*P^eq^) as a function of *w/c* for information-maximizing realizations of the six-state gene circuit model shown in Figure 4B. The plot reveals that IR-maximizing gene circuits consistently utilize energy to drive sharpness above its equilibrium limit (S*/*S^eq^>1), while precision is maintained at or below its equilibrium limit (P*/*P^eq^ ≲ 1). Moreover, the *degree* to which non-equilibrium gene circuits amplify S increases dramatically as *w/c* increases, from a factor of 2 when *w/c* ≈ 1 to a factor of 100 when *w/c* ≈ 10^4^ (Figure 5A). Thus, the key to understanding how energy increases IR at large *w/c* values lies in understanding transcriptional sharpness.

**Figure 5.**
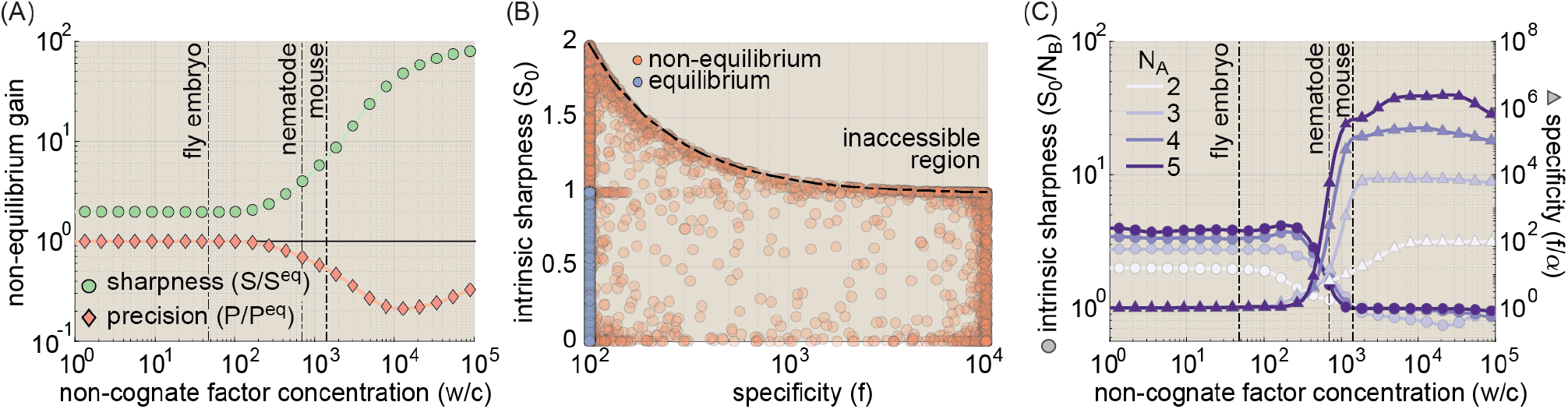
A shifting optimality landscape for information transmission. **(A)** Non-equilibrium gains in sharpness and precision as a function of w/c for six-state (N_B_ = 1, N_A_ = 1; Figure 4B) gene circuits found to drive maximum information rates. IR−maximizing gene circuits are drawn from optimal systems uncovered in the parameter sweeps from Figure 4E. Values above 1 indicate that the system is dissipating energy to enhance performance. The black line indicates a “break-even” point where the non-equilibrium value is equal to the equilibrium maximum. See Figure S4A for results for systems with N_A_ *>* 2. **(B)** Tradeoffs between intrinsic sharpness (S_0_) and specificity (*f*) for equilibrium and non-equilibrium networks (blue and red circles, respectively). Note that equilibrium gene circuits have no horizontal dispersion because all are constrained to have *f* = *α*. The black dashed line indicates the bound predicted by Equation 7. **(C)** Non-equilibrium gains in intrinsic sharpness and specificity for IR-maximizing gene circuits as a function of *w/c*. Values above 1 indicate that the system is dissipating energy to enhance sharpness or specificity. Note that the left and right axes have different scales. (*α* was set to 100 for all plots shown. For all parameter sweep results in A-C, transition rate and interaction term magnitudes, *k* and *η*, were constrained such that 10^*−*5^ ≤ *kτ*_*b*_ ≤ 10^5^ and 10^*−*5^ ≤ *η* ≤ 10^5^, where *τ*_*b*_ is the burst cycle time. *η*_*ab*_ and *η*_*ib*_ were further constrained such that *η*_*ab*_ ≥ 1 and *η*_*ib*_ ≤ 1, consistent with our assumption that the transcription factor activates the gene locus.)

The upper non-equilibrium limit on S can be expressed as a function of the specificity (*f*), such that

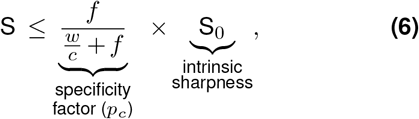

where the observed sharpness (S) bound breaks naturally into two pieces: the specific bound fraction, *p*_*c*_ (defined in Equation 4), and the intrinsic sharpness (S_0_), defined as a gene circuit’s normalized sharpness absent non-cognate factor binding (i.e., *w* = 0).

To probe the interplay between intrinsic sharpness and specificity, we employed parameter sweeps for the six-state system in Figure 4B. At equilibrium, this analysis indicated that intrinsic sharpness is constrained such that S_0_ ≤ 1 (consistent with Figure 3A) and confirmed that specificity is fixed at *α*. Indeed, we find that *f*^*eq*^ = *α* applies for *all* gene circuits operating at equilibrium irrespective of the number of binding sites or activation steps, placing strict limits on information transmission at equilibrium when *w/c* is large (see Appendix O.3).

Away from equilibrium, systems can overcome these constraints, achieving up to a two-fold increase in S_0_ and increasing specificity by up to an additional factor of *α* to reach an upper limit of *α*^2^ (Figure 5B). The observed 100-fold increase in *f* is comparable to the gain in the observed sharpness (S) in Figure 5A, suggesting that the sharpness gain at high *w/c* arises from non-equilibrium increases in specificity. Why not spend energy to *simultaneously* increase intrinsic sharpness by two-fold and specificity by 100-fold to achieve S*/*S^*eq*^ = 2 × *α* = 200? The simple answer is that non-equilibrium gains in intrinsic sharpness and specificity cannot be realized simultaneously. Instead, our analysis reveals a steep tradeoff between specificity and intrinsic sharpness away from equilibrium, with the maximum value of S_0_ = 2 only realizable when specificity is at its equilibrium level (*f* = *α*) and vice versa (Figure 5B). We find that the bound describing this tradeoff (black dashed line in Figure 5B) follows a simple analytic form, allowing us to express S as a function of the specificity, *f*, such that

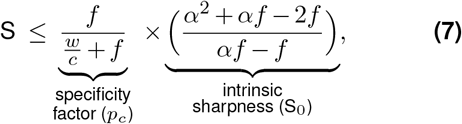

where we assume that *α* ≤ *f* ≤ *α*^2^. See Appendix P for a derivation of Equation 7. As with the non-equilibrium tradeoffs between sharpness and precision, this incompatibility stems from the fact that sharpness and specificity require distinct and incompatible underlying molecular architectures. Although we focused on the simple model shown in Figure 4B, we find similar non-equilibrium tradeoffs between *f* and S_0_ for more complex molecular architectures (Figure S4B). Thus, we conclude that these specificity gains come at the cost of diminished intrinsic sharpness.

The inexorable tradeoff between the intrinsic sharpness S_0_ and specificity *f* illustrated in Figure 5B means that gene loci must “choose” between allocating energy to maximize intrinsic sharpness and allocating energy to maximize specificity. To examine how the concentration of non-cognate factors shapes this tradeoff, we took IR-maximizing non-equilibrium gene circuits spanning the relevant range of *w/c* values for systems with 1–4 activation steps and calculated S_0_ and *f*. Figure 5C illustrates the relative non-equilibrium gains in intrinsic sharpness and specificity, respectively, for these circuits as a function of *w/c*.

Figure 5C reveals that the relative non-cognate factor concentration, *w/c*, defines a shifting optimality landscape. At low non-cognate factor concentrations, maximally informative gene circuits spend energy exclusively to maximize intrinsic sharpness (S_0_*/*N_B_ *>* 1 for all systems on the left-hand side of Figure 5C) at the cost of equilibrium specificity levels (*f/α* = 1). Thus, our model predicts that at low levels of non-cognate factor interference—as would be experienced, for instance, in developing fruit fly embryos—non-equilibrium mechanisms are not required to buffer against non-cognate factor interference, and allocating energy to maximize intrinsic sharpness constitutes the optimal regulatory strategy. However, once *w/c* surpasses the affinity factor *α*, IR maximization starts to disfavor sharpness (see decreasing S_0_ near *w/c* = 10^2^ in Figure 5C) and increasingly depends on enhancing specificity to non-equilibrium levels. Moreover, the presence of multiple activation steps dramatically increases the upper limit for non-equilibrium specificity, such that 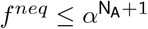 (Figure S4B). Together, these results indicate that the optimal molecular strategy for transmitting information is not fixed, but changes according to a scale set by the relative amount of non-cognate factor interference, *w/c*, and the kinetic binding differences between cognate and non-cognate factors, *α*.

### G. Predicting experimental signatures of non-equilibrium processes in transcriptional regulation

So far, we have demonstrated that energy dissipation can, in principle, increase the rate of information transmission in gene circuits. However, determining whether gene circuits mediating cellular decision making actually leverage energy dissipation to do so remains, to a large degree, an open challenge. Thus, we examined how simple experiments can identify signatures of non-equilibrium performance in real biological systems. For simplicity, we focused on the simple gene circuit in Figure 4B with one binding site and one molecular activation step, illustrating a broadly applicable set of experimental and analytical approaches that can be used to assess whether energy is harnessed to enhance transcriptional performance in real biological systems.

Recent works have shown that strict equilibrium limits on transcriptional sharpness can be calculated if the number of activator binding sites is known, suggesting that sharpness might serve as an accessible signature of non-equilibrium regulatory mechanisms (Estrada et al., 2016; Park et al., 2019). However, these studies did not consider off-target activation from non-cognate activator species. What happens when we account for the impact of such non-cognate factor binding? Equation 7 predicts that the upper S limit should decrease as *w/c* increases (blue and red dashed lines in Figure 6A), as confirmed by numerical parameter sweeps of S vs. *w/c* (blue and red circles). Thus, the upper sharpness limit is not absolute, but instead depends on the concentration of non-cognate factors in the cellular environment. This *w* dependence must be considered to accurately interpret experimental measurements.

**Figure 6.**
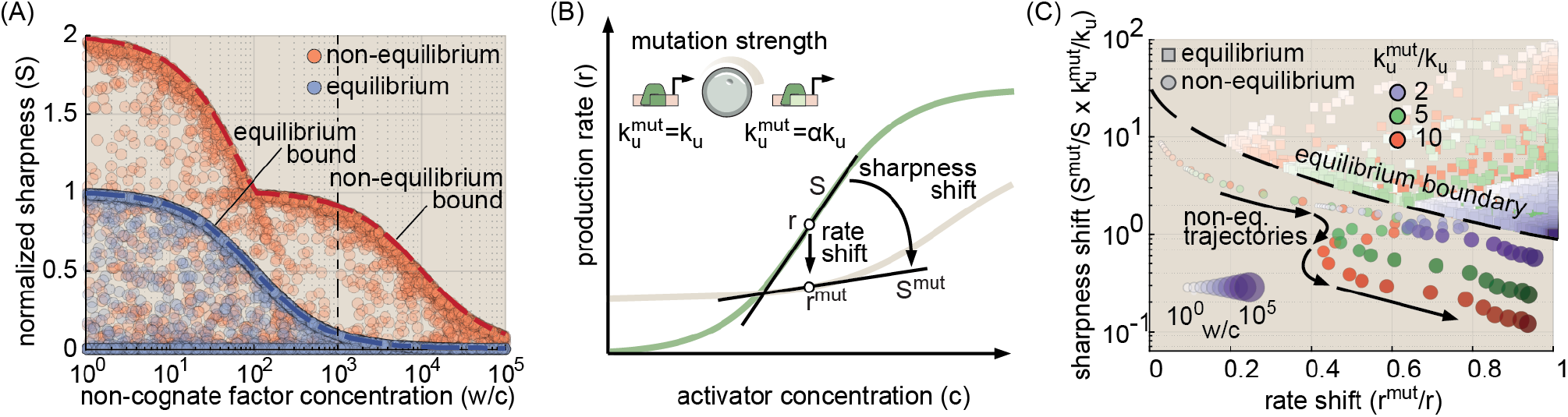
Experimental signatures of non-equilibrium processes in transcriptional regulation. **(A)** Observed sharpness as a function of *w/c* for equilibrium (blue circles) and non-equilibrium (red) gene circuits. The black dashed line indicates the point where *w/c* = 10^3^. **(B)** Illustration of proposed binding site perturbation experiments. Reducing site specificity is predicted to reduce both the observed sharpness, S, and the mRNA production rate 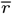. The strongest possible perturbation would entail a conversion from cognate specificity (*k*_u_) to non-cognate specificity (*αk*_u_). **(C)** Phase-space plot of predicted sharpness shift (normalized by 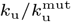) versus rate shift for equilibrium (squares) and non-equilibrium (circles) gene circuits at three binding site perturbation strengths. Note that we normalize the sharpness fold change by 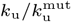, which allows us to plot results for different mutation strengths on the same y-axis. Shading indicates the *w/c* value (darker shades correspond to higher values). Additionally, the circle size indicates the *w/c* magnitude for non-equilibrium circuits. We see that, regardless of non-cognate concentration and perturbation strength, non-equilibrium systems do not cross the equilibrium boundary (dashed line). Results assume the initial transcription rate of the wild-type gene is at half-maximum 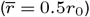. (For all parameter sweep results in A and C, transition rate and interaction term magnitudes, *k* and *η*, were constrained such that 10^*−*5^ ≤ *kτ*_*b*_ ≤ 10^5^ and 10^*−*5^ ≤ *η* ≤ 10^5^, where *τ*_*b*_ is the burst cycle time. *η*_*ab*_ and *η*_*ib*_ were further constrained such that *η*_*ab*_ ≥ 1 and *η*_*ib*_ ≤ 1, consistent with our assumption that the transcription factor activates the gene locus.)

For instance, consider the case where *w/c* = 10^3^ (black dashed vertical line in Figure 6A), a plausible value for mammalian systems (Friedlander et al., 2016; Cepeda-Humerez et al., 2015; Charoensawan et al., 2010). Our model predicts that the maximum achievable S for non-equilibrium gene circuits is 0.91, far exceeding the *true* equilibrium sharpness limit of 0.09 when accounting for the effects of non-cognate factor interference (blue dashed line in Figure 6A). However, *S* = 0.91 falls *below* the “naive” equilibrium bound of S = 1 that one would predict if *w* were not accounted for (see blue bound on far-left-hand side of Figure 6A, see also Figure S5A). Thus, failing to account for non-cognate factor interference could mask strong non-equilibrium signatures, highlighting the importance of incorporating regulatory cross-talk into transcription models. However, accurately measuring *w/c* may be challenging in many experimental settings, since *w* comprises the aggregate activity of all non-cognate activator species.

In light of this challenge, we propose a complementary experimental approach to search for signatures of non-equilibrium gene regulation that is more robust to uncertainty regarding the precise value of *w/c*. As illustrated in Figure 6B, this method involves measuring changes in gene expression at *C* = *c*^∗^ that result from point mutations to the activator binding site, which thereby lead to a higher unbinding rate, 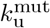, for cognate activators 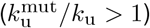. Whereas may be *w/c* difficult to estimate in many biological contexts, robust algorithms can predict changes in binding energies from the DNA sequence of transcription factor binding sites (Le et al., 2018), allowing for accurate predictions of how much a particular mutation will perturb the relative binding kinetics of a specific activator species. We employ two metrics to quantify the resulting change in gene expression: fold changes in the mRNA production rate 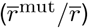 and in the normalized sharpness (S^mut^*/*S), each defined as the quantity corresponding to the mutated binding site divided by its corresponding wild-type value (Figure 6B).

To illustrate the method, we used our model to predict outcomes for the case where the wild-type gene circuit is expressing at half its maximum rate 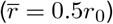. Overall, we find that IR-optimized non-equilibrium gene circuits are highly sensitive to changes in cognate activator specificity and that this sensitivity can be used to probe for non-equilibrium behavior. At low *w/c* levels (*w/c* ≲ 10^3^), mutated non-equilibrium circuits exhibit larger shifts in their transcription rate than can be achieved at equilibrium (Figure S5B). Meanwhile, when *w/c >* 10^3^, IR-optimized non-equilibrium systems experience a substantially larger sharpness decrease than even maximally sensitive equilibrium circuits (Figure S5C). Consequently, when combined, S^mut^*/*S and 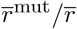 define a perturbation response space in which non-equilibrium gene circuits that transmit information at optimal (or near-optimal) levels are completely disjoint from equilibrium systems. This is illustrated in Figure 6C, which plots our model’s predictions for the sharpness fold change (S^mut^*/*S) vs. 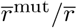 for three binding site perturbation strengths for equilibrium and non-equilibrium gene circuits (squares and circles, respectively). Despite the wide range of perturbation strengths and non-cognate factor concentrations examined, optimal non-equilibrium systems never cross the equilibrium boundary (dashed line). Thus, by measuring S^mut^*/*S and 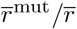, we can obtain clear-cut signatures on non-equilibrium regulation, even when *w/c* is unknown.

## Discussion

Gene regulation is central to cellular function. Yet, despite decades of biochemical and genetic studies that have established a reasonably complete “parts list” of the molecular components driving eukaryotic transcription (Kornberg, 2007), and despite recent advances in our ability to track how these pieces assemble in space (Nogales et al., 2017) and time (Lammers et al., 2020; Coulon et al., 2013; Lenstra et al., 2016), we nonetheless lack quantitative models that can predict how transcriptional control emerges from molecular interactions at the gene locus. Thermodynamic models of transcription, which assume that gene circuits operate at equilibrium, have been employed with considerable success to predict transcriptional control in the context of bacteria (Phillips et al., 2019). However, the presence of ATP-dependent processes—such as chromatin remodeling (Zhou et al., 2016), pre-initiation complex assembly (Taatjes, 2017), and Pol II initiation (Yan and Gralla, 1997)—within the eukaryotic transcriptional cycle suggests that equilibrium models may be insufficient to capture how eukaryotic gene circuits sense and respond to input transcription factor concentrations. Thus, there is an urgent need for theoretical frameworks that can probe how non-equilibrium mechanisms reshape the transcriptional input-output function and, ultimately, redefine the limits of transcriptional control.

Here, we employed simple kinetic models of transcription to investigate how energy dissipation within the transcriptional cycle impacts the rate at which a gene circuit drives cellular decisions. We found that biologically plausible rates of energy dissipation can drive significant gains in the information transmission rate and discovered that the regulatory mechanisms underlying these non-equilibrium gains change from increased sharpness to increased specificity depending on the level of interference in the cellular environment from non-cognate factor binding.

### Performance tradeoffs dictate limits of information transmission away from equilibrium

This work has established that, although energy dissipation can increase transcriptional sharpness, precision, and specificity *individually*, these gains cannot be realized simultaneously. For negligible non-cognate factor binding, we showed that IR is dictated by a tradeoff between sharpness (S) and precision (P). Although previous works have established that energy expenditure can boost sharpness (Estrada et al., 2016; Park et al., 2019) and, to a lesser extent, suppress transcriptional noise (Rieckh and Tkačik, 2014). As a result of this tradeoff, gene circuits must “choose” whether to spend energy to enhance sharpness or precision. For all models considered, we discovered that the information rate was maximized by systems that boosted transcriptional sharpness (not precision) above its equilibrium limit (Figure 3A, Figure S2A and B).

Similarly, our analysis revealed that non-equilibrium gains in specificity and sharpness cannot occur simultaneously (Figure 5B and Figure S4B). This incompatibility arises from the fact that intrinsically sharp systems are tuned to amplify concentration-dependent activator binding rates, whereas specific systems amplify differences in *unbinding* rates between cognate and non-cognate activator species. Our model predicts that *w/c* defines a shifting optimality landscape, wherein non-equilibrium gene circuits that maximize intrinsic sharpness drive the fastest decisions when *w/c* ≤ *α*, but the optimal strategy begins to shift from increasing sharpness to activator proofreading when *w/c > α* (Figure 5C). A recent study reported the potential for this kind of context-dependent shift from sharp to specific gene circuits (Grah et al., 2020), although sharpness was only investigated at its equilibrium limit. Here, we provide quantitative predictions for how IR-maximizing gene circuits navigate this sharpness-specificity tradeoff far from equilibrium.

### Activation steps amplify non-equilibrium performance gains

Another key finding of this work is that the presence of multiple activation steps, wherein multiple molecular components must engage to achieve transcription, can amplify non-equilibrium gains in transcriptional sharpness (Figure 3C). Our result is evocative of a recent study (Biddle et al., 2020) demonstrating that systems with multiple conformational degrees of freedom can achieve sharper, more flexible transcriptional input-output functions, although these systems still adhere to the fundamental equilibrium limitation that sharpness cannot exceed the number of activator binding sites (S ≤ N_B_). Thus, our findings further emphasize potential benefits of the conformational complexity of the eukaryotic gene cycle.

Consistent with previous results in the kinetic proofreading literature (Murugan et al., 2012), we also found that gene circuits with multiple activation steps can realize dramatic increases in transcriptional specificity when driven out of equilibrium (*f*), such that 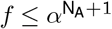 (Figure S4B). This result extends the findings of a recent work examining transcriptional specificity in systems with up to two activation steps (Shelansky and Boeger, 2020). Yet there exists an important asymmetry between sharpness and specificity: whereas the addition of activator binding sites can increase the sharpness *S* at equilibrium, energy dissipation constitutes the *only* route (short of altering activator binding sequences) for increasing specificity *f* above the intrinsic affinity factor *α*. Thus, for large *w/c*, energy dissipation overcomes a fundamental limitation of eukaryotic gene circuits—the lack of binding specificity—that no equilibrium mechanism can address.

### Equilibrium regulatory schemes may be sufficient in many real biological systems

While activator proofreading may be critical when *w/c* is large, our analysis suggests that it is unlikely to constitute a universal constraint on gene regulatory architectures. Indeed, even relatively simple equilibrium architectures with 3–5 binding sites should suffice to drive timely cellular decisions in “low-interference” systems such as the fruit fly embryo (Figure 4D). Moreover, while simple estimates based on genomic transcription factor abundances suggest that many eukaryotic systems can exceed the *w/c* = *α* interference limit, these estimates likely represent upper bounds on *w/c*, since different cell types selectively express distinct subsets of transcription factors (Choudhury and Ramsey, 2016; Lee et al., 2012; Henry et al., 2012). In addition, we note that the relative size of the concentration difference between *c*_1_ and *c*_0_ (*δc/c*) plays a key role in dictating the information transmission rate (Equation 2) and varies across different biological contexts. Thus, it would be interesting to use the quantitative tools presented in this work to enumerate the space of viable equilibrium and non-equilibrium gene circuit architectures for specific biological systems in which the relative magnitudes of *w/c* and *δc/c* are well established.

### Different frameworks for examining the impact of non-cognate factor binding

In considering the impact of non-cognate factor binding, we drew inspiration from a previous study examining competition between cognate and non-cognate transcription factors to bind and activate a single gene locus (Cepeda-Humerez et al., 2015). This formulation of the problem is distinct from the approach taken in two recent works, which addressed the problem of specificity from the perspective of a single activator species that interacts with two different gene loci: a cognate (with specific binding sites) and a non-cognate locus (without specific binding sites) (Shelansky and Boeger, 2020; Grah et al., 2020). While both approaches have proven fruitful, we favor the “single-locus” approach, since it captures the effects of competitive binding between different species, which are an unavoidable reality of crowded cellular environments.

Moreover, this shift in perspectives has meaningful consequences for our understanding of how off-target binding impacts gene regulation. A previous study found that the equilibrium limit of *f* = *α* could only be achieved at the cost of high levels of transcriptional noise (Shelansky and Boeger, 2020). Yet, we find that this tradeoff evaporates once competitive binding between cognate and non-cognate factors is considered, since *f* is fixed at *α* in this case (Figure 5B). The upper limits of transcriptional sharpness also decrease as *w/c* increases (Equation 7 and Figure 6A). Previous studies have reported transcriptional sharpness as a key potential indicator of non-equilibrium optimization (Estrada et al., 2016; Park et al., 2019). Our analysis reaffirms this idea but, crucially, reveals that one must consider the relative concentration of non-cognate factors (*w/c*) to accurately assess whether a particular system is performing above the equilibrium limit (Figure 6A and B). For instance, a sharpness of 0.9 falls below the equilibrium limit for the six-state gene circuit shown in Figure 4B when *w/c* ≈ 1, but is an order of magnitude above the limit when *w/c* ≈ 10^3^ (Figure 6A).

### Future directions

While we have considered gene loci with varying numbers of *specific* activator binding sites, real enhancers also contain significant stretches of “neutral” DNA with no binding sites, as well as weak activator sites that fall below typical thresholds used to identify specific sites (Vincent et al., 2016; Shahein et al., 2021). This focus on specific sites is widespread in theoretical studies of transcription (Estrada et al., 2016; Park et al., 2019; Cepeda-Humerez et al., 2015; Lammers et al., 2020), despite the well-established importance of weak binding sites in the context of certain genes (Shahein et al., 2021; Crocker et al., 2015; Farley et al., 2015). Moreover, recent efforts on synthetic enhancer reconstitution have pointed to the importance of supposedly neutral stretches of regulatory DNA (Vincent et al., 2016), and it seems theoretically plausible that these stretches, where cognate and non-cognate activator species bind with equal affinity, could have important effects on the input-output function in systems when *w/c > α*. We propose that the kinetic models utilized herein could readily be extended to feature some combination of specific and neutral sites. More ambitiously, the field would benefit from the introduction of continuous, rather than discrete, theoretical models that admit non-equilibrium phenomena while accounting for the reality that transcription factors interact with a continuum of sites along enhancer DNA.

Ultimately, the key to unraveling the molecular mechanisms by which genes sense and respond to transcription factor concentrations lies in the coupling of theoretical models with careful experimental measurements. To this end, we advocate for the expanded use of theoretically tractable synthetic enhancer systems in which the number and identity of binding sites are well established and intervening DNA sequences are carefully engineered to minimize binding specificity (e.g., using SiteOut (Estrada et al., 2016)). Several recent studies constitute promising initial steps in this direction (Reimer et al., 2021; Park et al., 2019; Vincent et al., 2016; Kim et al., 2021). Additionally, synthetic transcription factor systems, which can act orthogonally to endogenous regulatory networks, represent an intriguing experimental platform for investigating questions relating to transcriptional specificity (Kabadi and Gersbach, 2014; Crocker and Stern, 2013). Lastly, statistical methods that infer how transcription factor concentrations impact the kinetics of the transcriptional cycle (Zoller et al., 2018; Lammers et al., 2020; Corrigan et al., 2016; Bowles et al., 2022) hold promise for connecting macroscopic experimental measurements to theoretical models of the microscopic processes driving transcription. Looking ahead, holistic research efforts that integrate cutting-edge experiments, statistical methods, and theory will be key to bridging the as yet yawning gap between enhancer sequence and gene regulatory function.

## ACKNOWLEDGEMENTS

We are grateful to Jane Kondev, Sara Mahdavi, and Vahe Galstyan for substantial comments and discussion on the manuscript. Thanks also to Rob Phillips, Muir Morrison, and Ben Kuznets-Speck for their helpful discussion and insights at various stages of this project’s development. NCL was supported by NIH Genomics and Computational Biology training grant 5T32HG000047-18, the Howard Hughes Medical Institute, and by DARPA under award number N66001-20-2-4033. AIF was supported in part by an NSF Graduate Research Fellowship, NSF Grant No. PHY-1748958, the Gordon and Betty Moore Foundation Grant No. 2919.02, the Kavli Foundation, and by a Postdoctoral Fellowship from the Jane Coffin Childs Memorial Fund for Medical Research. HGG was supported by the Burroughs Wellcome Fund Career Award at the Scienti1c Interface, the Sloan Research Foundation, the Human Frontiers Science Program, the Searle Scholars Program, the Shurl and Kay Curci Foundation, the Hellman Foundation, the NIH Director’s New Innovator Award (DP2 OD024541-01) and NSF CAREER Award (1652236), an NIH R01 Award (R01GM139913) and the Koret-UC Berkeley-Tel Aviv University Initiative in Computational Biology and Bioinformatics. HGG is also a Chan Zuckerberg Biohub Investigator.

## Supplementary Figures

**Fig. S1.**
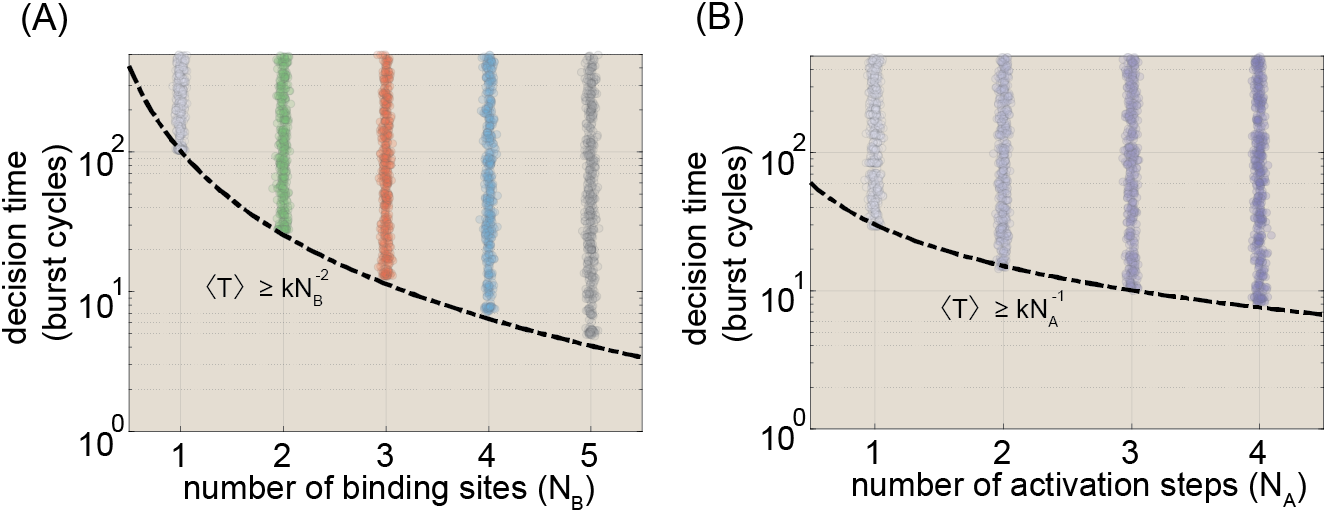
Decision times for different gene circuit architectures. **(A)** Parameter sweep results for equilibrium gene circuits with different numbers of activator binding sites. Black dashed line indicates lower limit of the decision time and is a function of the form 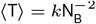, where k is a proportionality constant. **(B)** Plot of range of achievable decision times for non-equilibrium gene circuits with a single activator binding site (N_B_ = 1) as a function of the number of activation steps, N_A_. The dashed line indicates the lower decision time bound, and is a function of the form 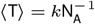. (All results shown assume an error probability of 32%. For parameter sweep results in A and B, transition rate and interaction term magnitudes, *k* and *η*, were constrained such that 10^*−*5^ ≤ *kτ*_*b*_ ≤ 10^5^ and 10^*−*5^ ≤ *η* ≤ 10^5^, where *τ*_*b*_ is the burst cycle time. *η*_*ab*_ and *η*_*ib*_ were further constrained such that *η*_*ab*_ ≥ 1 and *η*_*ib*_ ≤ 1, consistent with our assumption that the transcription factor activates the gene locus.)

**Fig. S2.**
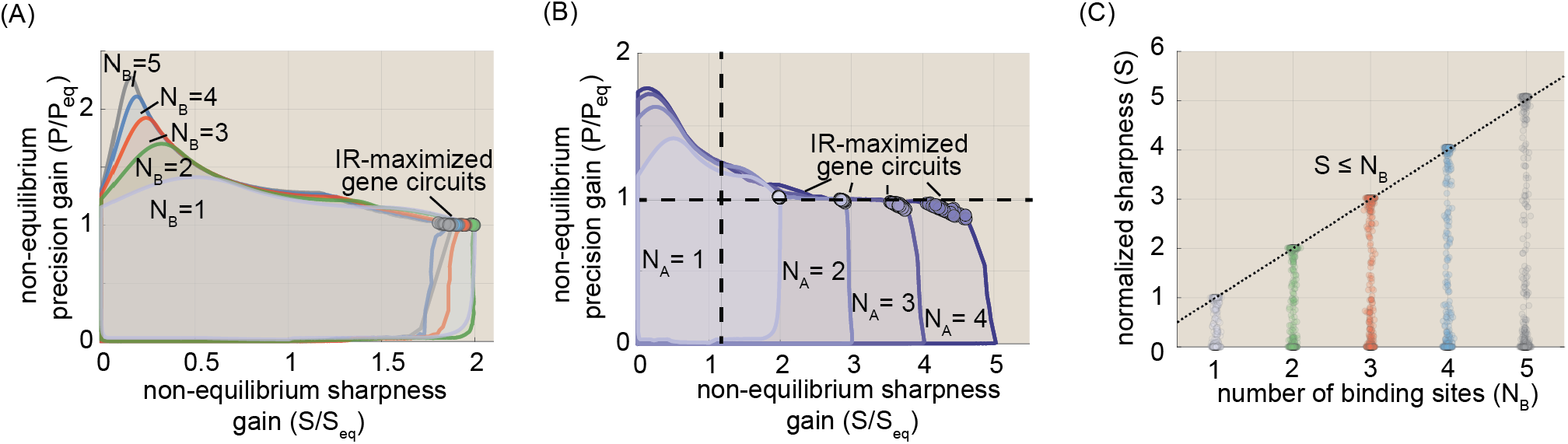
Tradeoffs between sharpness and precision persist for more complex gene regulatory architectures. **(A)** Non-equilibrium gains in sharpness and precision for gene circuits with different numbers of activator binding sites (N_B_) and one activation step. Shaded regions indicate achievable regimes for each system, as determined by no fewer than 10,000 unique simulated gene circuits. **(B)** Non-equilibrium gains in sharpness and precision for gene circuits with different numbers of activation steps (N_A_) and one activator binding site. **(C)** Scatter plots indicate sharpness levels for equilibrium gene circuits as a function of the number of binding sites. Bounding line is for a function of the form S = N_B_. (For parameter sweep results in A-C, transition rate and interaction term magnitudes, *k* and *η*, were constrained such that 10^*−*5^ ≤ *kτ*_*b*_ ≤ 10^5^ and 10^*−*5^ ≤ *η* ≤ 10^5^, where *τ*_*b*_ is the burst cycle time. *η*_*ab*_ and *η*_*ib*_ were further constrained such that *η*_*ab*_ ≥ 1 and *η*_*ib*_ ≤ 1, consistent with our assumption that the transcription factor activates the gene locus.)

**Fig. S3.**
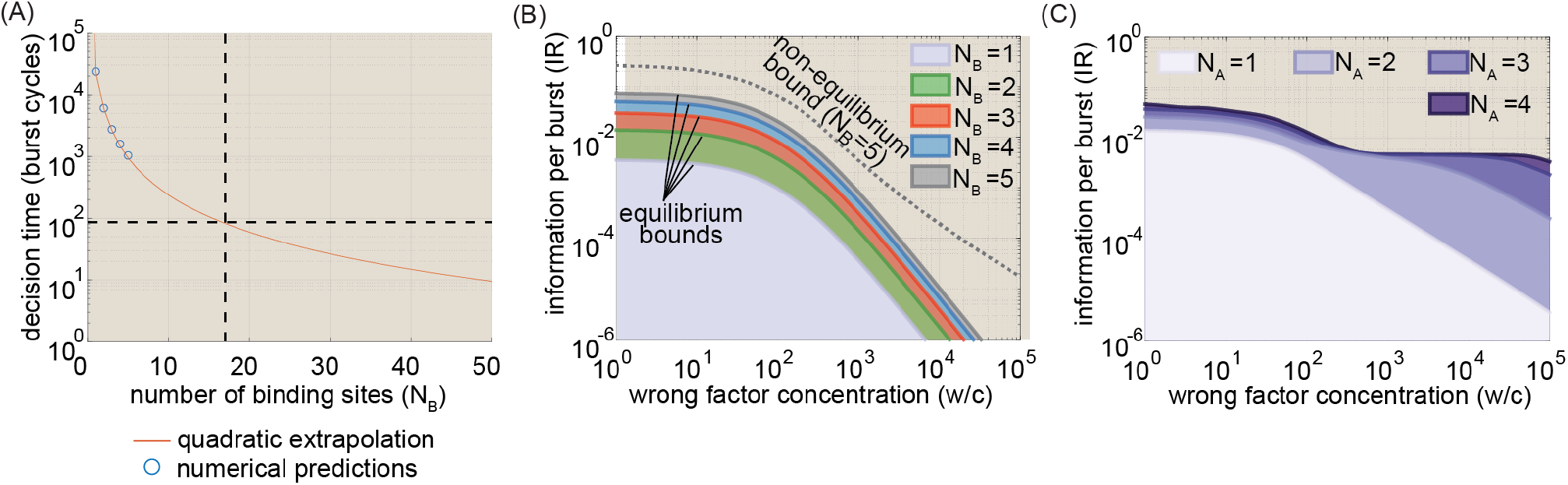
Supplemental analyses for the dependence of IR with non-cognate transcription factor interference. **(A)** Parameter sweep results showing the range of achievable information rates as a function of *w/c* for equilibrium gene circuits with 1-5 activator binding sites and one molecular activation step. **(B)** Sweep results for non-equilibrium gene circuits with 1-4 activation steps and a single activator binding site. **(C)** Extrapolation of minimum decision times for equilibrium gene circuits as a function of number of activator binding sites based on numerical results for circuits with 1-5 binding sites. Analysis indicates that at least 17 sites would be required to achieve plausible decision ties in the context of the mouse system. (For parameter sweep results in B and C, transition rate and interaction term magnitudes, *k* and *η*, were constrained such that 10^*−*5^ ≤ *kτ*_*b*_ ≤ 10^5^ and 10^*−*5^ ≤ *η* ≤ 10^5^, where *τ*_*b*_ is the burst cycle time. *η*_*ab*_ and *η*_*ib*_ were further constrained such that *η*_*ab*_ ≥ 1 and *η*_*ib*_ ≤ 1, consistent with our assumption that the transcription factor activates the gene locus.)

**Fig. S4.**
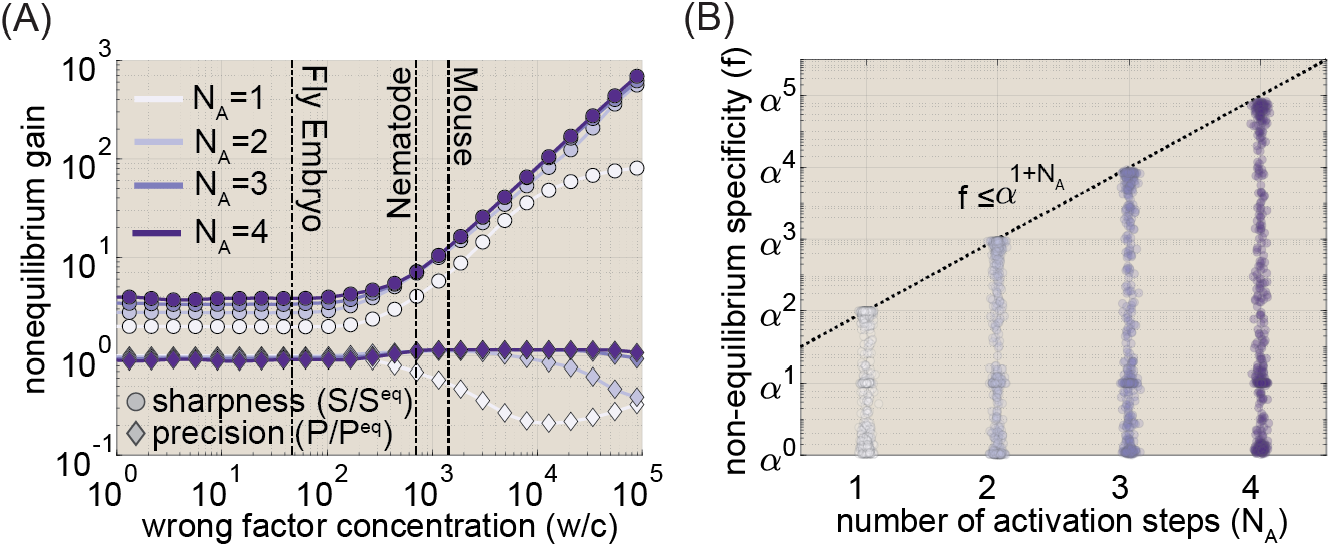
Supplemental results for main text Figure 5. **(A)** Non-equilibrium sharpness and precision gains for IR-maximizing gene circuits with 1-4 activation steps. **(B)** Range of achievable specificity values for non-equilibrium gene circuits with with 1-4 activation steps. (Transition rate and interaction term magnitudes, *k* and *η*, were constrained such that 10^*−*5^ ≤ *kτ*_*b*_ ≤ 10^5^ and 10^*−*5^ ≤ *η* ≤ 10^5^, where *τ*_*b*_ is the burst cycle time. *η*_*ab*_ and *η*_*ib*_ were further constrained such that *η*_*ab*_ ≥ 1 and *η*_*ib*_ ≤ 1, consistent with our assumption that the transcription factor activates the gene locus.)

**Fig. S5.**
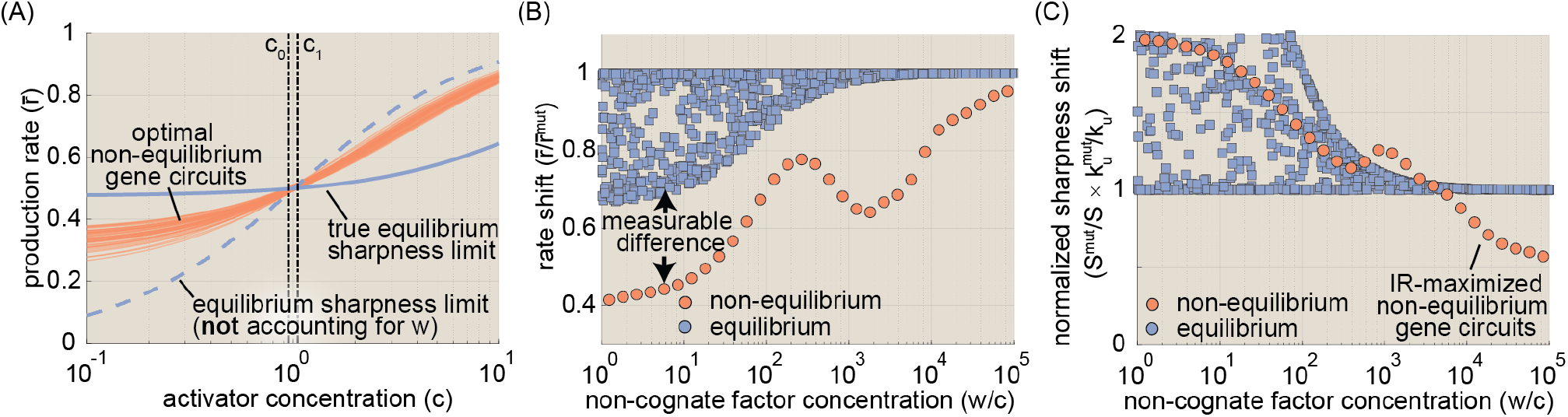
Experimental signature of energy expenditure. **(A)** Predicted induction curves for 50 near-optimal non-equilibrium gene circuits when *w* = 10^3^*c*^*∗*^, as well as the *actual* induction curves for the sharpest achievable equilibrium curve (solid blue line) and the (incorrect) limit when that would be predicted if *w* was not accounted for (dashed line). Note that red curves fall above the true equilibrium limit but below the naive limit. **(B)** Predicted shift in the production rate resulting from a binding site perturbation that doubles the unbinding rate 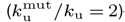 —equivalent to a energetic difference of 0.7 k_B_T—for equilibrium gene circuits (blue squares) and IR-maximizing non-equilibrium circuits (red circles). Note that non-equilibrium circuits are far more sensitive than equilibrium circuits when *w/c <* 10^4^ The shift becomes negligible at higher values, thus providing a clear signature of energy dissipation. **(C)** Predicted sharpness shift upon perturbing the activator binding site. The non-equilibrium shift becomes markedly larger than equilibrium limit when *w/c >* 10^3^. (For parameter sweep results in B and C, transition rate and interaction term magnitudes, *k* and *η*, were constrained such that 10^*−*5^ ≤ *kτ*_*b*_ ≤ 10^5^ and 10^*−*5^ ≤ *η* ≤ 10^5^, where *τ*_*b*_ is the burst cycle time. *η*_*ab*_ and *η*_*ib*_ were further constrained such that *η*_*ab*_ ≥ 1 and *η*_*ib*_ ≤ 1, consistent with our assumption that the transcription factor activates the gene locus.)

## Appendices

### A. Analytic expressions for key gene circuit characteristics

This section lays out analytic expressions for key quantities that play a central role in the investigations undertaken over the course of the main text. We do not repeat derivations for expressions that are treated separately elsewhere in these Appendices, and avoid re-deriving expressions from scratch, unless they are novel to this work.

#### A.1. The transition rate matrix and activity vector

Consider a gene circuit *g* that has *K* different microscopic states. We assume that microscopic transitions between the molecular states that make up *g* are Markovian, such that our system can be modeled as a continuous time Markov chain (CTMC). It follows that the steady-state behavior of *g* is fully determined by two quantities: the transition rate matrix, ***Q*** and the state activity vector, ***a. Q*** is a *K* × *K* matrix with off-diagonal elements that encode the rates with which the system switches between microscopic rates. For instance, *q*_*mn*_—the element in the mth row and nth column of ***Q***—gives the transition rate going from state n to state m. The diagonal elements of ***Q*** are negative, and are scaled such that each column of ***Q*** sums to 0. The activity vector ***a*** is a binary vector of length *K* that contains a “1” for each state that is transcriptionally active, and a “0” for inactive states. We assume that both ***Q*** and ***a*** are fixed in time.

#### A.2. State probabilities, transcription rate, and transcriptional noise

A first step to calculating virtually all gene circuit characteristics of interest is to obtain the steady-state vector, ***π***, which is a vector of length *K* that gives the steady state probability of finding the gene circuit of any one of the *K* microscopic states. We can obtain ***π*** by finding the right eigenvector (***v***_*R*_) of ***Q*** with an eigenvalue of 0,

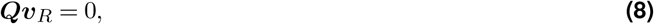

and imposing the additional constraint that the elements of ***π*** sum to 1, such that

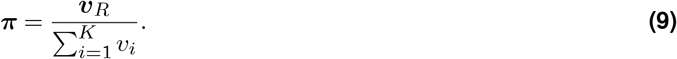

With this the steady state probability vector in hand, we can calculate the average transcription rate by taking the dot product of ***a*** and ***π***:

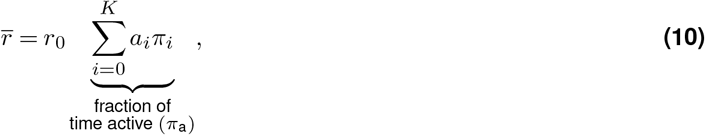

where we define the quantity indicated by the underbrace as the average fraction of time, *π*_a_, that the system spends in the active state. Throughout the course of this work, we assume that *r*_0_ is held fixed, such that the transcriptional activator may only impact transcription by modulating microscopic transition rates in ***Q*** to alter ***π***. Further, since we take Poisson noise from mRNA synthesis to be negligible (see Appendix D), the absolute magnitude of *r*_0_ is unimportant, and we set it to 1 for simplicity.

Next, we turn to obtaining an expression for the variance (noise) in gene expression. From Whitt 1992 (Whitt, 1992), we have that

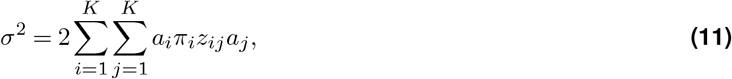

where *z*_*ij*_ is the element from ith row and jth column of what is known as the fundamental matrix, ***Z*** of our transition rate matrix, ***Q. Z*** is a *K* × *K* matrix that plays an integral role in the calculation of many key behaviors of a Markov chain. Once again drawing from Whitt, we can calculate ***Z*** using the formula

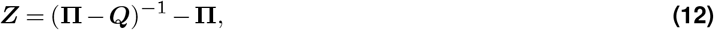

where **Π** is a *K* × *K* matrix with each row equal to *π*.

#### A.3. Using the fundamental matrix to calculate first passage times

First passage times provide a useful conceptual tool for connecting microscopic fluctuations, which often are unobservable, with emergent dynamical behaviors, such as transcriptional bursting. The fundamental matrix provides an invaluable tool for doing this in the context of arbitrarily complex transcriptional systems. Once again, we start with an expression from Whitt 1992 (Whitt, 1992) that relates off-diagonal elements of ***Z*** to first passage times between microscopic states:

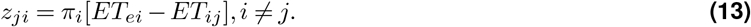

Here, *ET*_*ij*_ is the mean expected first passage time from state *j* to state *i* and *ET*_*ei*_ the first passage time to state *i* at equilibrium, defined as

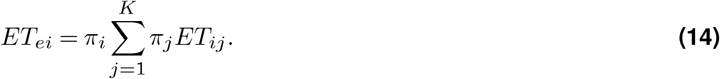

Now, from (Whitt, 1992) we also have that the diagonal elements of ***Z*** can be expressed as

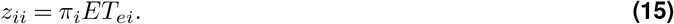

We can now combine Equations 13 and 15 to solve for the first passage time from state i to state j:

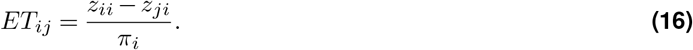

#### A.4. Calculating the burst cycle time

First passage times are intimately related to a quantity of central importance throughout the text: the burst cycle time, *τ*_*b*_, defined as the average time required for a system to complete one ON → OFF → ON cycle (Figure 1D). This is trivial in the case of a simple two state system with a single OFF and ON state and rates *k*_on_ and *k*_off_ (Figure A11). In this case, the burst cycle time is simply

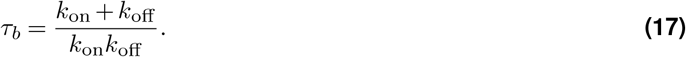

The calculation becomes less trivial for systems with larger numbers of states, however. Fortunately, the concepts outlined above provide us with the tools necessary to derive a generic expression for *τ*_*b*_ that applies to systems of arbitrary complexity.

The essence of the procedure lies in calculating effective off and on rates (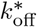 and 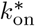) from ***Q*** using first passage times. We go through this procedure in detail for 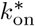 and note that the same approach applies for 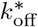. The activity vector ***a*** partitions our system into *M* OFF states and *N* ON states. To calculate 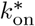, the first step is to estimate the expected amount of time it will take for the system to reach an ON state (any ON state) from each OFF state. We can do this by defining a new transition rate matrix, ***Q***^OFF^, that has dimensions *M* + 1 × *M* + 1. The off-diagonal elements of the first M rows an M columns of ***Q***^OFF^ are simply equal to the microscopic rates from ***Q*** that lead from one of the M OFF states to another OFF state. Together, these molecular states constitute a single coarse-grained OFF state.

The final row and column, however, are different and contain total fluxes into and out of all ON states from each OFF state. An element in the final row of ***Q***^OFF^ is given by

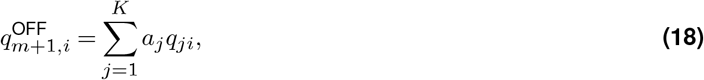

where *a*_*j*_ is the jth element of the activity vector, *q*_*ij*_ is a microscopic rate from the original transition rate matrix, and we assume the state i is in the set of OFF states. Thus, we see that each element of the last row of ***Q***^OFF^ gives the total flux from *all* OFF state into the ON conformation. The elements of the final column have a complementary definition:

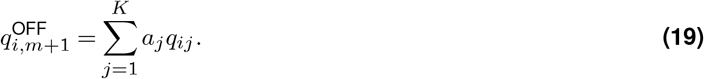

With our condensed transition rate matrix thus defined, we can use Equations 8 and 9 to calculate ***π***^OFF^ and Equation 12 to calculate ***Z***^OFF^. Then, we can use Equation 16 to obtain a vector ***et***^*ON*^ of length *M*, where each element *i* is defined as the expected first passage time from OFF state i back into *any* of the ON states. Specifically, we have that each element, *i*, is given by

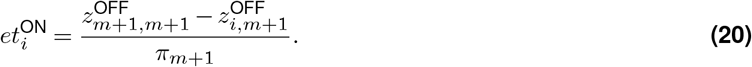

Thus, we have obtained a vector, ***et***^ON^, of expected mean first passage times out of each OFF state into the set of *N* active transcriptional states. But how do we weight the different passage times in this vector to arrive at an overall average expectation for the amount of time required for the system to turn back ON following a transition into an OFF state? It’s tempting here to use the stead-state probabilities of each OFF state given by ***π***, but this is actually not correct.

Instead, the key is to recognize that each OFF state should be weighted by the rate at which ON states switch into it. In other words, we weight OFF states by the probability that they are the initial state the system reaches upon switching out of the ON conformation; the gateway into the OFF states. Mathematically, we encode these weights using the flux vector ***f*** ^OFF^, which has *M* elements, each defined as

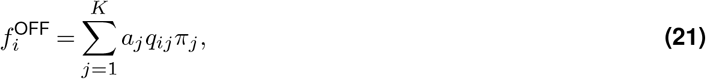

where *a*_*j*_ is the *jth* element of the activity vector ***a*** (1 for ON states and 0 otherwise), *q*_*ij*_ is the transition rate from state *j* to state *i*, and *π*_*j*_ is the steady-state probability of state *j*.

Finally, we combine this expression with Equation 20 to obtain an expression for the average reactivation time as a flux-weighted average of the first passage times out of each OFF state:

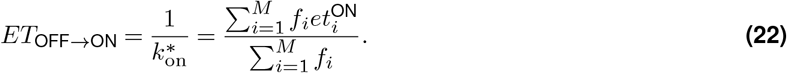

As noted above, the calculations for 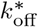 follow precisely the same logic, with the roles of the OFF and ON states switched. After this is done, the total burst cycle time, *τ*_*b*_, is simply

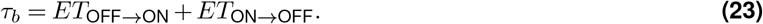

Equation 22 is useful because it allows us to relate the (potentially quite complex) microscopic dynamics of a transcriptional system to emergent bursting timescales observed in live imaging experiments (Lammers et al., 2020). To our knowledge, this is the first time that take this flux-weighted first passage time approach is applied to the modeling of burst dynamics. We hope that the expressions provided here will prove useful to others seeking to pursue similar projects in the future.

Finally, a useful feature implied by Equation 22 and Equation 23 is that the absolute size of *τ*_*b*_ scales inversely with the microscopic rates in ***Q***, such that we can decrease *τ*_*b*_ by some scaling factor *λ* by simply multiplying ***Q*** by *λ*. We use this trick to renormalize all time-dependent metrics calculated over the course of our parameter sweeps to have units of burst cycle time. This is done by calculating *τ*_*b*_ for each new model realization we generate, and then multiplying its transition rate matrix by this quantity to generate a normalized rate matrix, namely

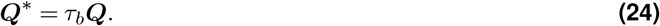

The adjusted matrix, ***Q***^∗^, is then used to calculate all relevant gene circuit characteristics.

#### A.5. A generic expression for the rate of energy dissipation

Equation 1 gives an expression for the rate of energy dissipation (also termed entropy production), Φ, in the context of the four-state model shown in Figure 1C. This is a special case of a more general formula for Φ that applies to arbitrary molecular architectures. From (Lang et al., 2014; Lebowitz and Spohn, 1999), we have

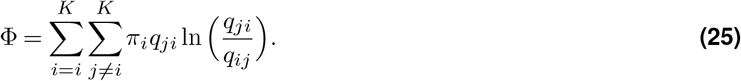

We use Equation 25 to calculate all energy dissipation rates given throughout the main text. In the case of the simple four-state system shown in Figure 1B, we have from (Lang et al., 2014) that Equation 25 simplifies to

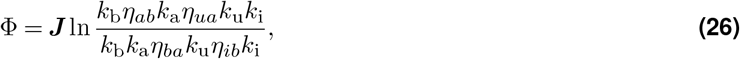

which further simplifies to

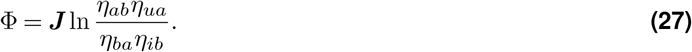

Here ***J*** is the net cycle flux, a quantity with units of inverse time which encodes the rate at which the system completes extra cycles in the clockwise (***J*** *>* 0) or counterclockwise (***J*** *<* 0) directions. Mathematically, ***J*** is given by

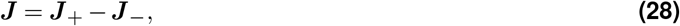

where ***J*** _+_ gives the average rate at which the system completes one full cycle in the clockwise direction (i.e., setting out from state 0 to state 1, reaches state 0 from state 4), and ***J*** ___ is defined analogously. In terms of microscopic quantities, for any system with a single loop we can define ***J*** as

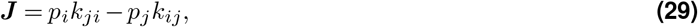

where *k*_*ji*_ denotes the transition rate from state *i* to state *j* and *J*_*ij*_ corresponds to the net transition flux between the two states.

### B. Gaussian noise approximation

Throughout this work, we make the simplifying assumption that the intrinsic noise in accumulated mRNA levels due to transcriptional bursting is approximately Gaussian. In this section, we use stochastic simulations to put this assumption to the quantitative test. The Markov chain central limit theorem states that the distribution of a quantity that is a function of a Markov chain (such as the transcription rate, 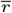), will become approximately Gaussian as the number of iterations becomes large (Geyer, 2011).

The question, then, is whether can expect the accumulated transcriptional output to approach this limiting Gaussian distribution within timescales that are relevant to the decision times discussed in this work. To determine this, we used stochastic simulations (Gillespie, 1977) to track the distribution of the accumulated output of 500 random realizations of the four-state system shown in Figure 1C for 5,000 burst cycles. Each realization had a unique set of transition rates and, correspondingly, a unique average rate of transcription, 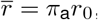, where *π*_*a*_ indicates the fraction of time the system spends in a transcriptionally active molecular state and *r*_0_ is the rate of transcript initiation when active. For each model realization, we ran 100 stochastic simulations. We used these simulations to track the distribution of the apparent average transcription rate for each model realization as function of accumulation time. Figure A1A shows the apparent mean rate across 100 simulations for a single illustrative gene circuit realization. Inset histograms indicate distribution of apparent transcription rates at different time points. As expected, we see that the apparent rates are initially highly dispersed; however, even after 25 burst cycles, we see that 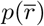 has become a much narrower, roughly symmetrical distribution that appears approximately Gaussian.

To systematically assess the rate of convergence to normality, we utilized the simple One-sample KolmogrovSmirnov test (“kstest”, (Massey, 1951)), which tests the null hypothesis that a vector of transcription outputs from realization *i* at time *t*, ***r***_*i*_(*t*), is drawn from a normal distribution. The test returns a *p* value corresponding to the probability of observing ***r***_*i*_(*t*) if the transcriptional output were truly Gaussian. In standard implementations *p* ≲ 0.05 is taken to constitute strong evidence that the output is *not* Gaussian. Thus, to assess convergence to normality, we tracked this *p* value over time for each of the 500 gene circuit realizations.

Figure A1B shows the average kstest p-values across 10 different sets of gene circuits, grouped by their average rate of transcription. In all cases, we see that noise profiles rapidly converge towards normality, such that all systems cross the (relatively conservative) threshold of *p* = 0.1 within 5 burst cycles (dashed line in Figure A1B). Gene circuits near the tail ends of the induction curve (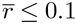 and 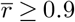) take the longest to converge, which is likely because it takes longer for distributions near the boundaries to become symmetric about their mean; yet even these converge rapidly.

The fastest decisions discussed in the main text (Figure 4D and E), and most decision times considered are significantly longer than the time for Gaussian convergence revealed by Figure A1B). Thus, we conclude that the Gaussian noise approximation invoked throughout this work is justified.

**Fig. A1.**
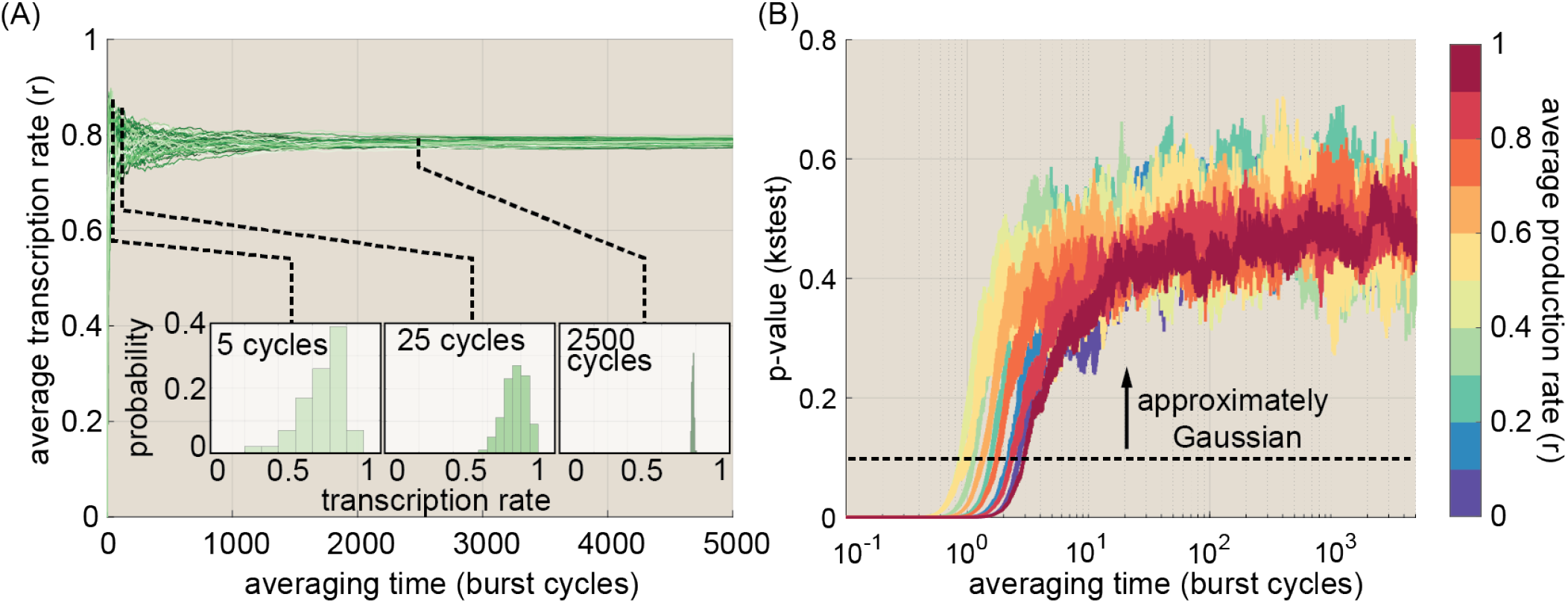
Testing the validity of the Gaussian noise approximation. **(A)** Illustrative plot showing average transcription rate as a function of the averaging time across 100 stochastic simulations of one illustrative realization of the four-state model gene circuit. Inset histograms show distribution of apparent rates at three different time points. We see that, as the accumulation time increases, the distributions get tighter and appear more Gaussian in shape. **(B)** Plot showing p-values of one-sample Kolmograv-Smirnov test. Different colors indicate average trends for systems with different average transcription rates. We see that systems near the low and high ends of the induction curve converge to Guassian form most slowly, but even these cross the *p* = 0.1 line within a handful of burst cycles. Error bars indicate bootstrap estimates of standard error calculated for each group. (For stocahstic simulations shown in A and B, transition rate and interaction term magnitudes, *k* and *η*, were constrained such that 10^*−*2^ ≤ *kτ*_*b*_ ≤ 10^2^ and 10^*−*2^ ≤ *η* ≤ 10^2^, where *τ*_*b*_ is the burst cycle time. *η*_*ab*_ and *η*_*ib*_ were further constrained such that *η*_*ab*_ ≥ 1 and *η*_*ib*_ ≤ 1, consistent with our assumption that the transcription factor activates the gene locus.)

### C. Deriving the rate of information transmission for a gene locus

Motivated by (Siggia and Vergassola, 2013), we define the rate of information transmission as the time derivative of the expected Kullback-Leibler (KL) divergence between the two hypotheses (*C* = *c*_0_ and *C* = *c*_1_), given some accumulated mRNA level *m*, such that

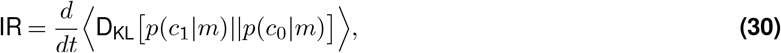

where P(*c*_0_ | *m*) and P(*c*_1_ | *m*) indicate (respectively) the conditional likelihood that the true value of *C* is *c*_0_ and *c*_1_ given the observed output *m*, and where the angled brackets indicate that we are dealing with the expected value of D_KL_ across many replicates. We refer readers to information theory reference materials for a formal definition of D_KL_ (see, e.g., (Cover and Thomas, 2006)); however, at an intuitive level it can be regarded as measuring how different two probability distributions are from one another. Thus, with Equation 30, we define the rate of information production as the rate at which the two possibilities (*c*_1_ or *c*_0_?) become distinguishable from one another given the observed “evidence” (*m*).

We can write out the expected KL divergence from Equation 30 more explicitly as the weighted sum of log probability ratios:

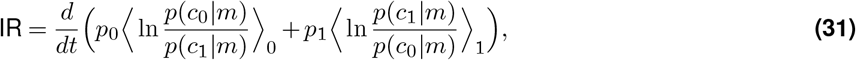

Where ⟨…⟩ _*i*_ indicates the expectation taken assuming the true value of *C* to be *c*_*i*_ and where *p*_0_ and *p*_1_ indicate the priors on the true value of *C*, taken to be equal moving forward (*p*_1_ = *p*_0_ = 1*/*2). This formulation provides intuition for the sense in which IR is the information rate: as the conditional probabilities of the observed output given the true (numerators) and false (denominators) hypotheses about *C* diverge in favor of the true hypothesis, the log ratio terms will become large and positive. Thus a positive derivative corresponds to positive information production.

However, here we must recall that our focus here is to understand how the molecular architecture of gene loci impacts the transcriptional response and, ultimately, IR. Thus we wish to work in terms of *p*(*m* | *c*)—the conditional distribution of observed mRNA outputs given some input—rather than *p*(*c* | *m*). To do this, we make use of Bayes’ Theorem. We have:

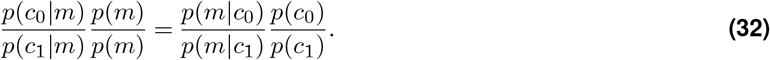

This expression becomes an equality if we assumed equal prior probabilities for our two hypotheses (*p*(*c*_0_) = *p*(*c*_1_)):

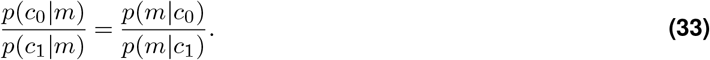

Thus, we can use Equation 33 to rewrite Equation 31 as:

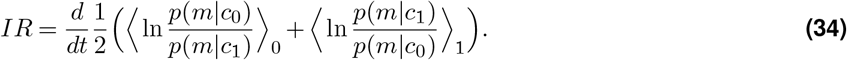

We can think of the conditional probabilities, *p*(*m* | *c*_*i*_), in Equation 34 as representing the full *stochastic* transcriptional response to some input activator concentration *c*_*i*_. When these are approximately Gaussian (a condition discussed above in Appendix B), it becomes a straightforward exercise to solve for the expected log ratios in Equation 34. We will solve for the case when *C* = *c*_1_ in full. The *c*_0_ case proceeds in precisely the same fashion. To start, we have

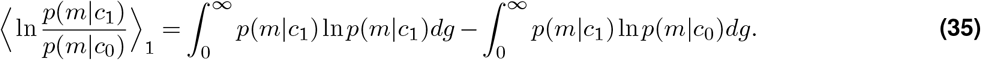

Recall that *m* = *rt* is Gaussian with probability density function:

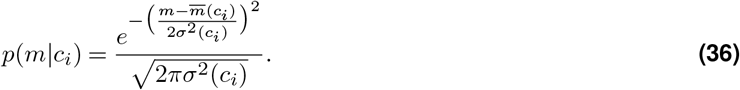

Plugging Equation 36 in for ln *p*(*m*|*c*_1_) yields

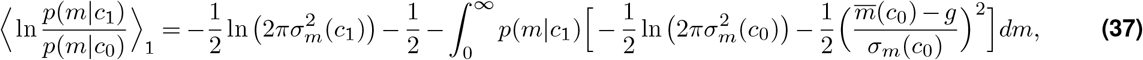

Where we’ve recognized that the first integral will simply yield the standard expression for the entropy of a Gaussian random variable. Pulling constant factors out of the second integral leads to

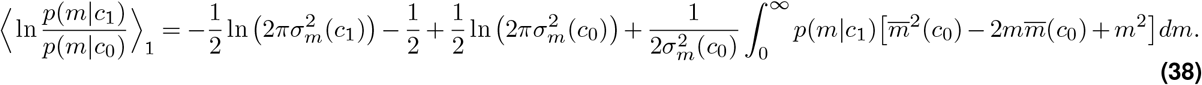

Simplifying and recognizing that 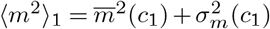 leads to:

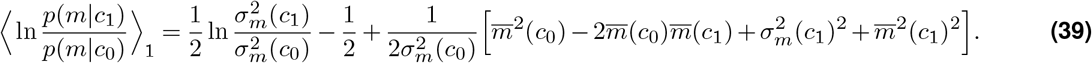

Finally, we recall that *m* = *rt* and 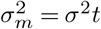, obtaining

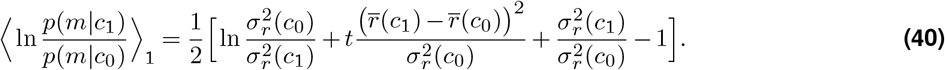

Performing the same procedure for the case where *c* = *c*_0_ yields:

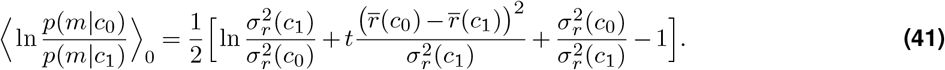

Plugging Equation 40 and Equation 41 into Equation 34 and taking the derivative with respect to time yields

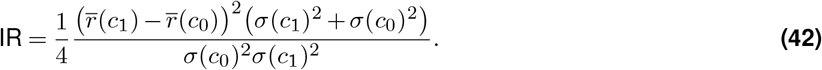

Next, if we assume that the difference between *c*_0_ and *c*_1_ is small (as stipulated in the main text), then *σ*(*c*_0_) ≈ *σ*(*c*_1_) ≈ *σ*^2^(*c*^∗^) and 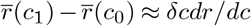, leading to

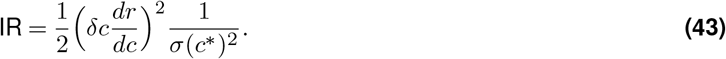

Finally, we invoke the definitions of sharpness and precision given in Figure 1B, which leads to Equation 2 from the main text:

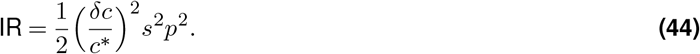

### D. Poisson noise from mRNA synthesis is negligible relative to noise from bursting

In this section, we provide support for the claim, made in Main Text Section B, that Poisson noise due to mRNA synthesis is negligible relative to noise from transcriptional bursting. We take as our starting point Equation 68 from Appendix J,

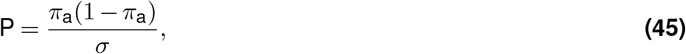

Which relates the normalized precision, P, to the bursting noise, *σ*, and the fraction of time a gene circuit spends in transcriptionally active states, *π*_a_. From Figure 3A, we see that *P* ≤ 1 for the four-state gene circuit shown in Figure 1C when the system is out of equilibrium, which, from Equation 45, implies that

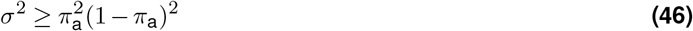

4 state system.

Thus, Equation 47 gives a lower bound for the intrinsic variance in gene expression that arises due to transcriptional burst fluctuations at the gene locus. To see how to relate this to noise from mRNA synthesis, we need to take two more steps. First, we must recall that we are working in units of the burst cycle time, *τ*_*b*_. Second, we must further recall that we set the actual rate of mRNA synthesis, *r*_0_, equal to 1 throughout the main text. We must do away with these simplifications in order to relate *σ*^2^ to synthesis noise. Accounting for these simplifications, the full expression for the noise floor, in “real” time units and accounting for the true rate of mRNA synthesis is

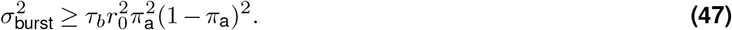

Now, if we assume mRNA synthesis to be a Poisson process (following, e.g., (Shelansky and Boeger, 2020)), we t this component of the variance is simply equal to

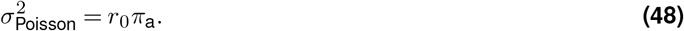

The key thing to notice about Equation 48 is that mRNA synthesis noise is *independent* of the bursting timescale *τ*_*b*_. Thus, as *τ*_*b*_ increases, 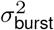 will increase in magnitude relative to 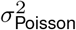. Figure A2A and B illustrate this fact, showing predicted bursting and mRNA synthesis variance components, respectively, as a function of the bursting time scale *τ*_*b*_ and the activity level (*π*_a_). All calculations assume an mRNA synthesis rate of 20 mRNA per minute,a rate based off of estimates from the fruit fly (Lammers et al., 2020) and that is consistent with measurements from other systems (Tantale et al., 2016). From Figure A2A, we see that 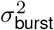 peaks at *π*_a_ = 0.5 and increases dramatically as we move rightward along the x-axis and the burst cycle time increases. We emphasize that this represents a *lower* bound for maximally precise non-equilibrium gene circuits; most systems (including IR-optimized systems) will lie above this bound. In contrast Figure A2B shows that noise from mRNA synthesis scales linearly with *π*_a_, and is constant in *τ*_*b*_.

**Fig. A2.**
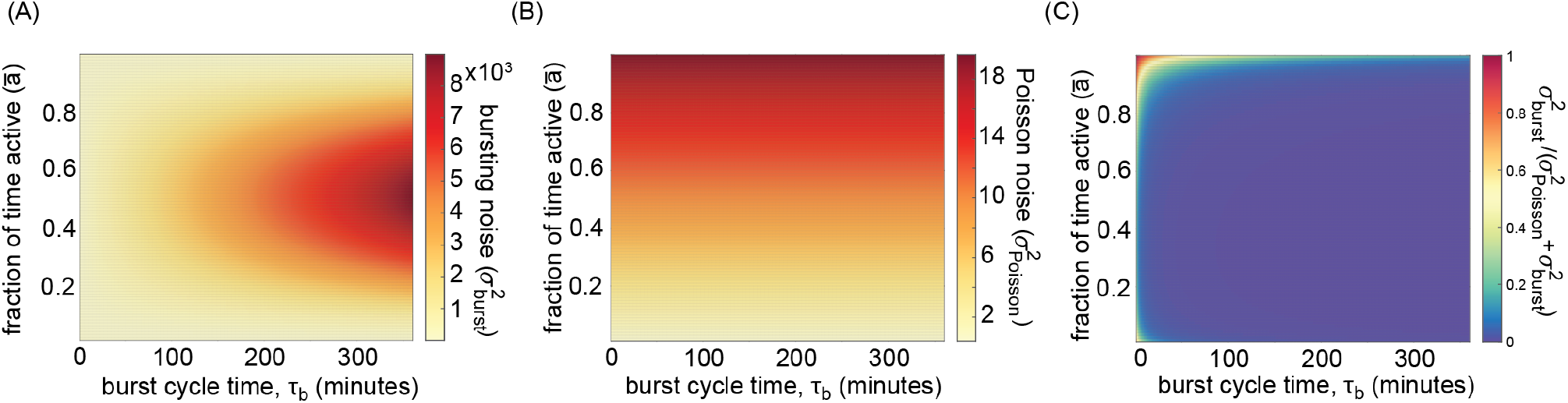
Determining the contribution from mRNA synthesis noise. **(A)** Heatmap showing lower bound of bursting component of variance for the non-equilibrium four-state model shown in Figure 1C as a function of the fraction of time spent in the active state (*π*_a_) and the burst cycle time (*τ*_*b*_). **(B)** Heatmap showing predicted variance component arising from mRNA synthesis. **(C)** Predicted relative contribution of mRNA synthesis noise to total intrinsic noise levels in gene expression. Note that contribution is only significant for rapidly bursting systems near the saturation point. (All calculations assume an mRNA synthesis rate of 20 per minute, in keeping with estimates from (Lammers et al., 2020).)

The total gene expression noise level is given by

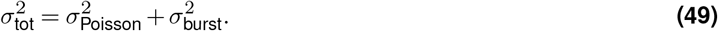

We can use this expression to calculate a lower bound on the relative contribution of mRNA synthesis noise to the overall intrinsic variance in gene expression. Figure A2C shows the results of this calculation. We see that, with the exception of rapidly bursting systems near the saturation (*π*_a_ ≈ 1), the contribution from Poisson noise due to mRNA synthesis is negligible. Thus, we conclude that noise from transcriptional bursting constitutes the dominant source of gene expression noise for the vast majority of the parameter regimes relevant for the investigations in this paper, and that our decision to neglect Poisson noise from mRNA synthesis is reasonable.

### E. The Sequential Probability Ratio Test

Over half a century ago, Wald conceived of the Sequential Probability Ratio Test (SPRT) as a solution to the problem of making accurate decisions between two hypotheses, *H*_1_ and *H*_0_ in “real time” as relevant data is accruing (Wald, 1945). Shortly thereafter, it was established that SPRT represents the optimal approach to sequential decision problems involving binary decisions (Wald and Wolfowitz, 1948), meaning that it requires the fewest observations to achieve a desired level of accuracy. In this framework, a downstream receiver (in our case, downstream genes or other cellular processes) tracks the accrual of some signal (mRNA, and eventually protein) over time and compares how likely this accrued signal is under the two hypotheses to be distinguished (e.g., high or low activator concentration). In this work, we use the optimal nature of SPRT to set lower bounds on decision times that could be achieved given the transcriptional output of model gene loci. The essence of the test lies in tracking the relative likelihoods of our two hypotheses (*C* = *c*_1_ and *C* = *c*_0_) over time as more and more transcriptional output, *m*, accrues:

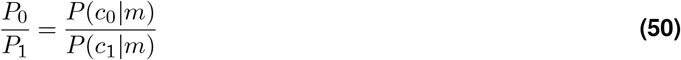

Figure A3A shows a stochastic simulation of how this ratio evolves over time for the output of a single model gene circuit. Although the true concentration in this case is *c*_0_, we see that the two hypotheses are essentially indistinguishable early on. This is because the range of possible outputs given high and low activator concentrations overlap significantly early on (leftmost panel of Figure A3B). However, as more and more time passes, the expected outputs (*m*) given the two possible inputs (*c*_1_ and *c*_0_) start to separate. We see that the ratio in their likelihoods diverges more and more in favor of *c*_0_ (*P*_0_*/P*_1_ *>>* 1), corresponding to a higher and higher degree of certainty that *c*_0_ is the correct choice.

This divergence, however, is non-monotonic and noisy, which reflects the stochastic nature of protein production at a single gene locus. It has been shown that the noisy divergence of the log of the probability ratio (which we will call *ℒ*) can modeled as a 1-D diffusive process with average drift IR (Siggia and Vergassola, 2013) given by

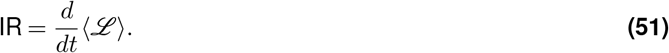

In this framework, a “decision” is made when *ℒ* crosses a so-called “decision boundary” (horizontal dashed lines in Figure A3A). Siggia et al showed that the Gaussian diffusion approximation could be used to obtain an analytic expression for the expected time needed to make a decision. From Equation 15 in the supplement of (Siggia and Vergassola, 2013), we have that:

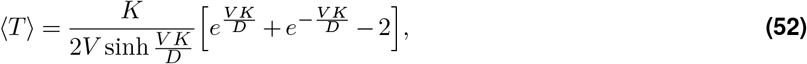

where *V* is the same as IR from above (and in the main text), *D* encodes the diffusivity of decision process (essentially, how large the fluctuations are about its mean drift trajectory), and *K* is related to the log of the error tolerance parameter *ε*, such that

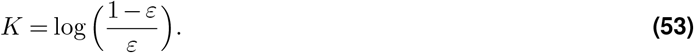

We note that Equation 52 assumes equal priors regarding the likelihood of *c*_1_ and *c*_0_, and also assumes equal error erances for choosing incorrectly in either case (Desponds et al., 2020).

If we take the accumulated transcriptional output of our gene circuit, *m* = *rt*, to be approximately Gaussian (see pendix B), then it can be shown that *D* has the form:

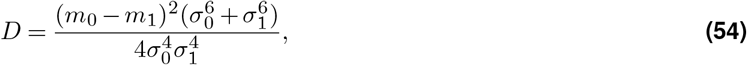

where *m*_*i*_ and *σ*_*i*_ give the mean and variance in the accumulated transcriptional output, given that *C* = *c*_*i*_. From Equation 42 in Appendix C, we also have that

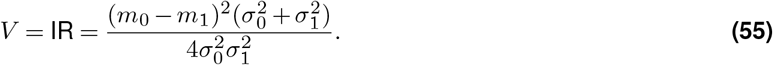

In a different context (exponential distributions, rather than Gaussian), Desponds and colleagues (Desponds et al., 2020) demonstrated that *D* ≈ *V* when the difference between hypothesis—*δc/c*^∗^ in our case—is small. From Equations 54 and 55, we see that this also holds for the Gaussian case: when *c*_1_ and *c*_0_ are sufficiently close, *σ*_1_ and *σ*_0_ will be approximately equal, such that:

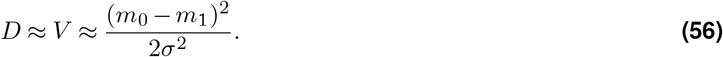

As demonstrated by (Desponds et al., 2020), when *D* ≈ *V*, Equation 52 simplifies dramatically, yielding

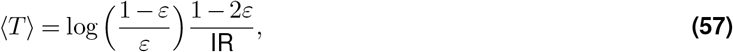

which is Equation 3 from the main text. For correctness, we use the full expression (Equation 52) to calculate all decision time quantities shown in the main text. However, since Equation 57 holds quite well for the 10% concentration difference considered here, we give the simpler expression in the main text to aid the reader’s intuition.

### F. Implementation of parameter sweep algorithm

In this section, we describe the parameter sweep algorithm employed throughout this work to enumerate the performance bounds of gene circuit models. We note that this approach is based off of an algorithm previously employed by Eck & Liu et al. (Eck et al., 2020) to explore the behavior of non-equilibrium models of transcription (see also, (Estrada et al., 2016)). Figure A4A illustrates the key steps in this numerical procedure. First, an initial set of gene circuit realizations (typically comprised of 1,000 variants) is generated by sampling random values for each transition rate in the system. We then calculate the performance metrics of interest (S and P for the example in Figure A4A) for each gene circuit realization. This defines an initial set of points (Figure A4A, Panel i) that collectively span some region in 2D parameter space with area *a*_1_.

Next (Panel ii), we subdivide parameter space into *N* different bins along the X and Y axes, with N dictated by the total number of points (10 ≤ *N* ≤ 50). We subsequently calculate the maximum and minimum point in each X and Y slice (Panel iii). Finally, we randomly select candidate gene circuit models from these boundary points and apply small perturbations to each transition rate to generate a new set of random variants (iv). In general, these variants will lie close to the original model in 2D parameter space and, thus, close to the current outer boundary of parameter space. The key to the algorithm’s success is that some of these variants will lie *beyond* the current boundary (blue points in Figure A4A, Panel iv). This has the effect of extending the boundary outward, leading to an increase in the surface area spanned by our sample points (panel iv). As a result, cycling through steps ii-iv amounts to a stochastic edge-finding algorithm that will iteratively expand the boundary spanned by sample points outward in 2D parameter space until some analytic boundary is reached.

**Fig. A3.**
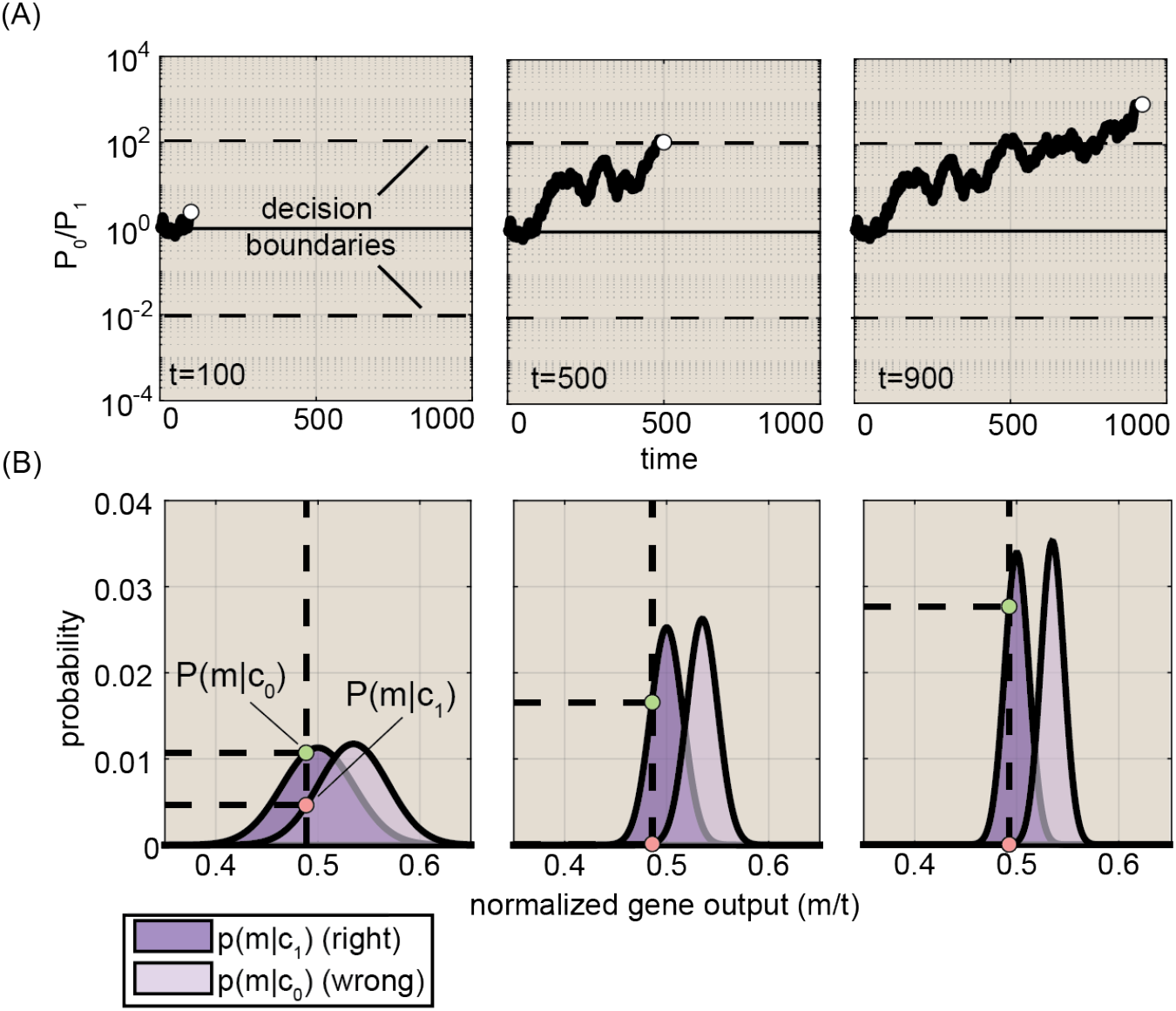
The Sequential Probability Ratio Test. **(A)** Panels show stochastic trajectory of the relative probabilities of *c*_0_ and *c*_1_ over time, given the observed output of some gene circuit (illustrated in Figure 1E). **(B)** Panels illustrating expected distributions of transcriptional outputs, *m*, for each concentration at different time points normalized by the total time over which the gene circuit has been active, *t*. Note how the distributions narrow and separate as time progresses.

The panels in Figure A4B show snapshots of the sweep algorithm’s progress exploring sharpness vs. precision parameter space for non-equilibrium realizations of the four-state gene circuit (Figure 1C). Figure A4C shows the total area spanned by the sample points for this run as a function of sweep iteration. By eye it appears that most of salient parameter space has been explored by step 10 of the algorithm, but we are quite strict with our convergence criteria. We will only terminate a sweep at step *t* if (*a*_*t*_ − *a*_*t*−2_)*/a*_*t*−2_ ≤ 0.001 and (*a*_*t*−1_ − *a*_*t*−3_)*/a*_*t*−3_ ≤ 0.001. In this case, this convergence criterion is met following step 25, leading to the final set of sample points shown in Figure A4D. In general, we run all sweeps until the above criterion is met or some pre-specified maximum number of iterations (usually 50) is reached.

#### F.1. Numeric vs. symbolic metric calculations

The algorithm outlined in Figure A4A is predicated upon the ability to rapidly calculate performance metric quantities (e.g., S and IR) given a set of transition rate magnitudes. Wherever possible, we use symbolic expressions to perform these calculations; however, this is only feasible for the simple four and six state systems depicted in Figure 1C and Figure 4B. For more complex models, it is infeasible to perform the symbolic operations required to obtain closed-form symbolic expressions. As a results, we use numerical calculations to arrive at performance metrics for all higher-order models.

#### F.2. Enforcing equilibrium constraints

In this work, we make frequent use of comparisons between equilibrium and non-equilibrium gene circuits in order to elucidate how energy expenditure alters gene-regulatory performance. A key step in performing parameter sweeps for equilibrium gene circuits is ensuring that transition rates adhere to the constraints imposed by detailed balance. For the simple four state model shown in Figure 1C, this process boils down to ensuring that the product of the four transition rates moving in a clockwise direction about the square is equal to the product of the four counterclockwise rates. As shown in Appendix A.5, this amounts to enforcing the constraint that

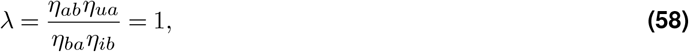

where the *η* factors on the top and bottom of the left-hand-side expression correspond to regulatory interaction terms that modify transition rates in the clockwise and counterclockwise directions, respectively, and where *λ* is the flux factor that captures the relative magnitudes of clockwise and counterclockwise transitions.

To enforce this constraint during the course of a parameter sweep, we add a step to the process outlined above. New gene circuit realizations are generated as before, but now, following its generation, we calculate the initial flux factor, *λ*_0_ for each new realization using Equation 58. In general this quantity will not equal one for the new realizations (*λ*^∗^≠ 1). To fix this, we then multiply *η*_*ba*_ and *η*_*ib*_ each by a factor of 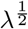, which leads to a modified system that adheres to the constraint laid out in Equation 58. Next, we check the modified terms to ensure that they adhere to magnitude constraints (typically 10^−5^*/τ*_*b*_ ≥ *η*_*i*_ 10^−5^*/τ*_*b*_) and pass all qualifying rates along to the next step in the sweep iteration (step ii in Figure A4A). Finally, we note that, although we have focused on the simple four state system, our assumption that all binding and activation reactions are identical (see Appendix I) ensures that the exact same approach holds for all higher-order models (N_A_ *>* 1 or N_B_ *>* 1) considered in this work.

**Fig. A4.**
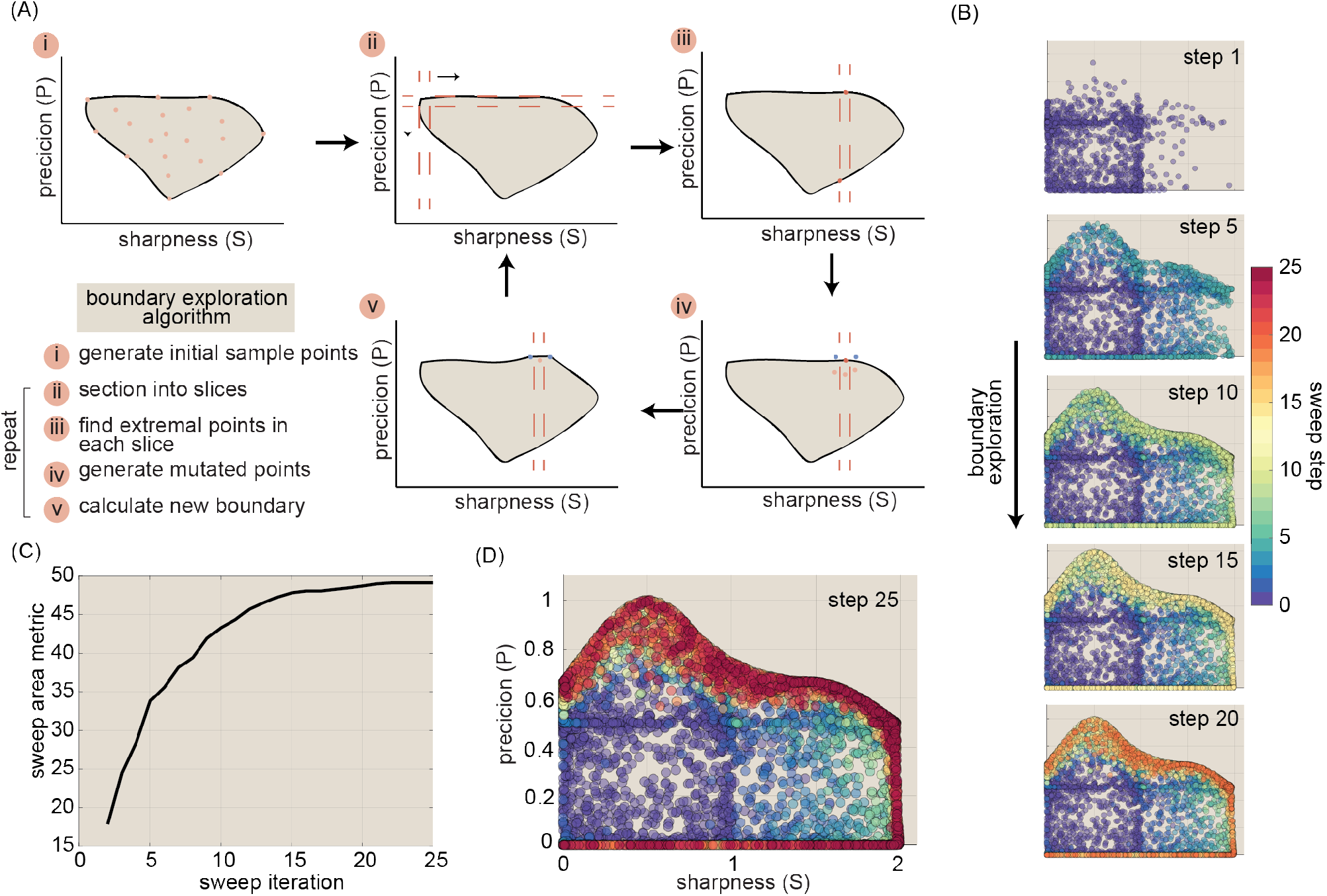
A simple stochastic edge-finding algorithm for numerical parameter sweeps. **(A)** Schematic illustrating key steps in our parameter sweep approach (see text for details). This panel has been adapted with permission from (Eck et al., 2020). **(B)** Sequence of snapshots showing progress of sweep algorithm across a single run for the case of normalized sharpness (S) versus normalized precision (P). Circle color indicates the sweep step on which it was generated. **(C)** Plot showing 2D surface area spanned by sample points over time. **(E)** Plot showing final set of sample points obtained by the sweep algorithm.

### G. Testing the convergence characteristics of the parameter sweep algorithm

Here, we discuss results from a series of tests designed to assess the convergence of our sweep algorithm for key scenarios examined in the main text. This task is the most straight-forward when the algorithm is employed for “two-boundary” sweeps, such as S vs. P (Figure 3A) and *S*_0_ vs *f* (Figure 5B), where both parameters examined adhere to finite performance bounds and, thus, where the 2D region of accessible parameter space has a finite area. In this case, our general approach will be to assess whether independent runs of the algorithm (i) converge prior to the 50 run limit and (ii) reach a consistent final estimate for the area of 2D space that is attainable for different model architectures. The task becomes more complicated for “one-boundary” sweeps, such as IR vs. Φ (Figure 2A, C, and D) and IR vs. *w/c*, where only a single parameter (IR in each case) has a finite upper bound and the other (Φ and *w/c*) is limited only by bounds imposed externally as a part of sweep specification. We will begin by assessing convergence for the simpler two-sided case, and will turn thereafter to examining one-sided cases.

#### G.1. Sharpness vs. Precision sweeps

Figure 3A and Figure S2A and B show results for parameter sweeps examining tradeoffs between normalized sharpness (S) and normalized precision (P) for systems with 1-5 binding sites and 1-4 activation steps. We note that Figure 3C and Figure S2C also derive from these parameter sweep results. Across the board, we find that nearly all independent runs of the sweep algorithm converge according to the definition laid out above (Figure A5A and B). Moreover, for simpler architectures, we find that all independent sweep runs converge to essentially the same total area. For instance, Figure A5B shows normalized area as a function of sweep step for 500 non-equilibrium realizations of the baseline four-state model, indicating that all runs terminate near the global maximum found across all runs (dashed line). We take this as strong evidence that the algorithm is consistently exploring the full extend of 2D parameter space.

**Fig. A5.**
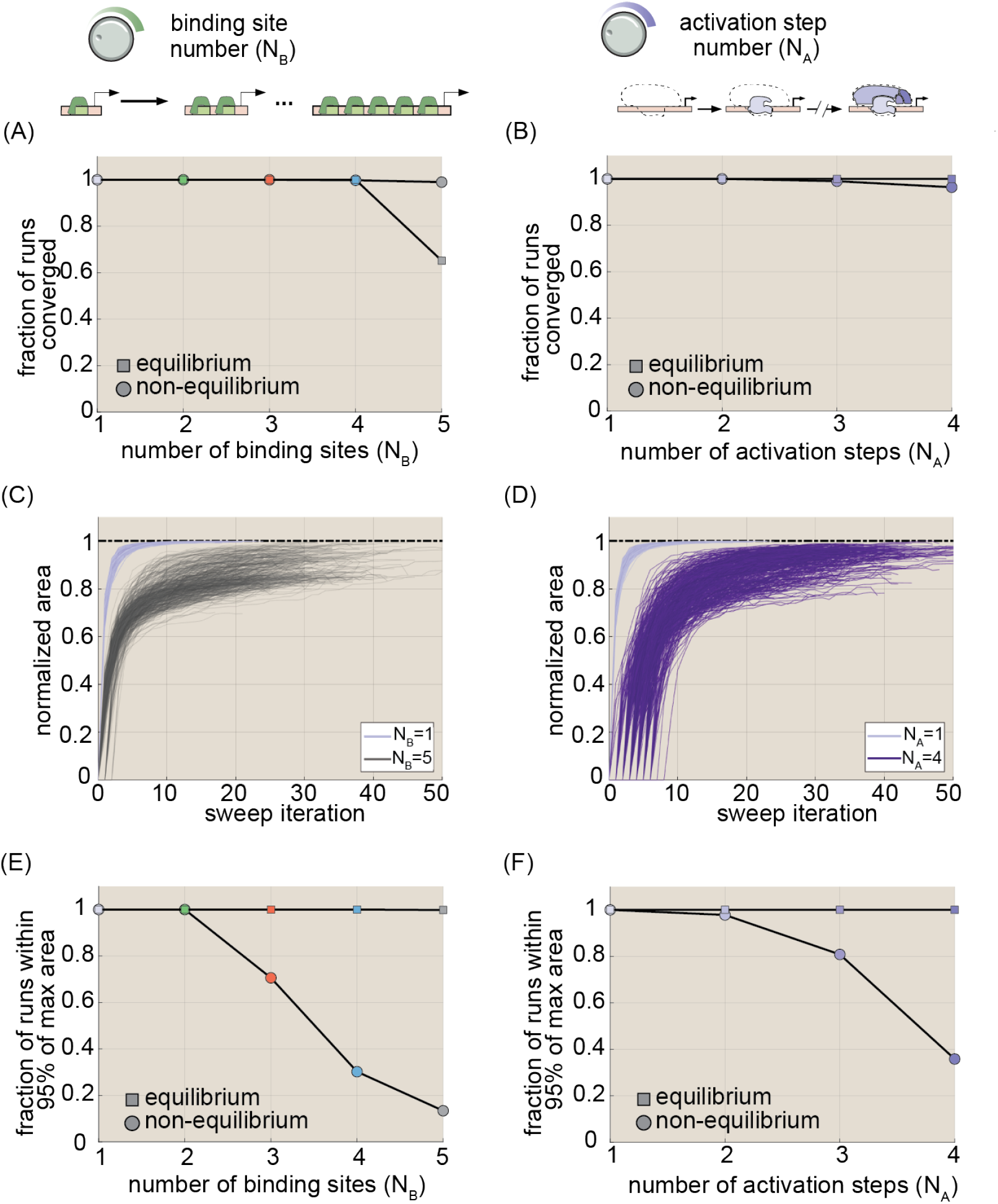
Convergence results for S vs. P parameter sweeps. **(A-B)** Plots showing fraction of parameter sweeps that met convergence criteria for multi-binding site and multi-activation step models, respectively. Squares indicate results for equilibrium models and circles indicate non-equilibrium models. **(C-D)** Plots of area vs. sweep step for different model architectures. Note that the area corresponding to the first (iteration=1) is not recorded by the algorithm, and so has been estimated in each case via linear interpolation. Staggered starts apparent for N_B_ = 5 and N_A_ = 5 models indicate cases where model initialization were aborted one or more times due to an insufficient number of gene circuits meeting quality control criteria. **(E-F)** Fraction of parameter sweeps having a final area within 95% of the global maximum for multi-binding site and multi-step models, respectively. (All results were calculated using 500 independent runs of the sweep algorithm for each model architecture. Transition rate and interaction term magnitudes (*k* and *η*) were constrained such that 10^*−*5^ ≤ *kτ*_*b*_ ≤ 10^5^ and 10^*−*5^ ≤ *η* ≤ 10^5^, where *τ*_*b*_ is the burst cycle time. *η*_*ab*_ and *η*_*ib*_ were further constrained such that *η*_*ab*_ ≥ 1 and *η*_*ib*_ ≤ 1, consistent with our assumption that the transcription factor activates the gene locus.)

As might be expected, the task of exhaustively exploring parameter space becomes more difficult as models become more complex. Note the larger spread in outcomes for the non-equilibrium five binding site (N_B_ = 5) and 4 activation steps (N_A_ = 4) models in Figure A5C and D, respectively. Nonetheless, we find that a significant number of sweeps converge to a consistent maximum area, even for the most complex models considered. Figure A5E and F give the total fraction of sweeps having a final area within 95% of the global maximum as a function of binding site number and activation step number, respectively. First, we see that *100%* of sweeps for equilibrium models uniformly meet this standard for all model architectures considered (squares in Figure A5E and F). Second, our analysis indicates that, even for the extrema (N_B_ = 5 and N_A_ = 5), 13% and 36% of total runs, respectively (67 and 179 sweeps), still achieve final areas comparable to the global maximum, suggesting that the algorithm still does an adequate job of exploring parameter space in these cases.

**Fig. A6.**
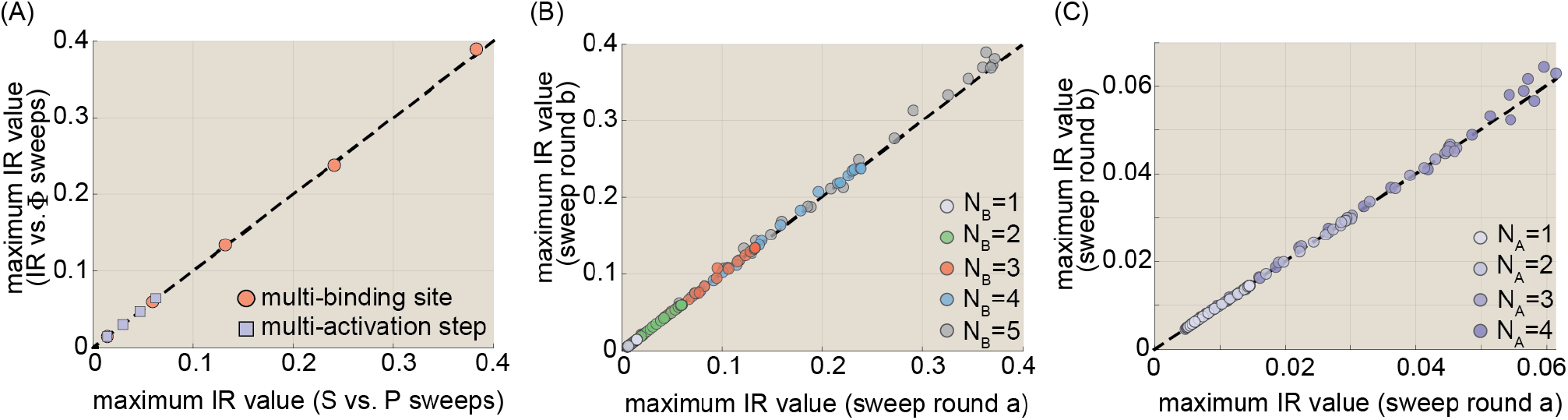
Convergence results for IR vs. Φ parameter sweeps. **(A)** Scatter plot comparing maximum information rate estimated from S vs. P and from IR vs. Φ sweeps. **(B-C)** Scatter plots comparing results for the upper IR bound at different points along the curves shown in Figure 2C and D from two independent rounds of parameter sweeps comprising 200 and 500 separate runs, respectively. Points reflect IR maxima for Φ values ranging from 0.1k_B_T to 5000k_B_T. (Transition rate and interaction term magnitudes (*k* and *η*) were constrained such that 10^*−*5^ ≤ *kτ*_*b*_ ≤ 10^5^ and 10^*−*5^ ≤ *η* ≤ 10^5^, where *τ*_*b*_ is the burst cycle time. *η*_*ab*_ and *η*_*ib*_ were further constrained such that *η*_*ab*_ ≥ 1 and *η*_*ib*_ ≤ 1, consistent with our assumption that the transcription factor activates the gene locus.)

#### G.2. Information vs. energy sweeps

Next, we turn to the one-sided sweeps. First, let’s consider the IR vs. Φ sweep results shown in Figure 2A, C, and D. Because Φ has no natural barrier in parameter space, the convergence metrics considered above do not provide reliable indicators of model convergence. Instead, we make use of the fact that the IR vs. Φ and the S vs. P parameter sweeps should function (either directly or indirectly) to uncover the maximum achievable non-equilibrium information rate for each model architecture. Thus, as a basic test of sweep performance, we checked for the consistency between IR estimates derived from these different sweep modalities. As illustrated in Figure A6A, we find excellent agreement between the maximum IR values derived from the S vs. P (x-axis) and IR vs. Φ (y-axis) parameter sweeps for all model architectures considered. This provides one indication the IR vs. Φ sweeps are fully exploring the relevant parameter space.

As a second check, we compared the IR vs. Φ bounds derived for two separate rounds of parameter sweeps (“round a” and “round b”) comprised of 200 and 500 independent parameter sweeps, respectively. We reasoned that, if our algorithm is accurately recovering the true IR vs. Φ bound for each model architecture, this bound (i) should be replicable across different parameter sweep rounds and (ii) should be insensitive to the precise number of sweep runs per round. For each model architecture, we calculated the maximum IR value returned by sweep rounds a and b for 30 different rates of energy dissipation ranging from 0.1k_B_T (close to equilibrium) to 5000k_B_T (upper limit of x axis in Figure 2D). Figure A6B and C show the results of this exercise for multi-binding site and multi-activation step models, respectively, indicating excellent agreement between different sweep round for all model architectures. This demonstrates that our information vs. energy bounds are highly replicable across different rounds of sweeps. The consistency across round comprised of significantly different numbers of runs provides further evidence that we are conducting a sufficient number of independent sweeps (≥ 200) per run. Taken together, these results and the results from the preceding paragraph provide strong evidence that our algorithm is robustly recovering accurate IR vs. Φ bounds for all models considered.

#### G.3. Information vs. w/c sweeps

Finally, we turn to the parameter sweep results for information (and, correspondingly, decision time) as a function of wrong-to-right activator concentration (*w/c*) shown in Figure 4C-E. We note that the results shown in Figure 5A and C are also derived from these sweeps. Like Φ, *w/c* has no intrinsic boundary in parameter space and, thus, swept area provides a poor indication of convergence. Fortunately, in addition to treating *w/c* as a sweep parameter, we can also conduct 2D parameter sweeps where *w/c* is set at a constant value (e.g., *w/c* = 1000 in Figure S4B). Thus we cross-validate the IR vs. *w/c* bounds returned by the sweeps from Figure 4 by conducting separate sweeps of IR vs. 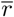 (the mean transcription rate) at different *w/c* values (illustrated in Figure A7A). These sweeps *do* converge, with an average of 80% of runs reaching 93% of the global maximum.

Figure A7B and C show the results of this comparison for three different values of *w/c*: 10, 10^2^, and 10^3^. We focus on the architectures depicted in Figure 4, namely equilibrium systems with 1-5 binding sites (and one activation step) and non-equilibrium systems with 1-4 activation steps (and one binding site). We also test convergence for the non-equilibrium gene circuit with 5 binding sites and 1 activation step shown as a dashed line in Figure 4D. In most cases, we find good agreement between the two methods, suggesting that the IR vs. *w/c* sweeps are generally returning accurate estimates for the IR vs. *w/c* bound. We do note a couple of exceptions, however. First, we see that that IR vs. *w/c* sweeps appear to underestimate the upper IR bound to a significant degree for the non-equilibrium model with 4 activation steps when *w/c* = 10 (circle in upper right-hand corner of Figure A7C). This indicates that the IR vs. *w/c* sweep is performing sub-optimally in this case. However, since this deviation occurs in the extreme low interference regime and our focus in Section E lies on model performance at higher *w/c* levels (*w/c* ≳ 100), where our sweep algorithm performs reliably, it does not impact any conclusions drawn throughout the course of the main text. We note that the IR vs *w/c* sweeps similarly underestimate the IR bound non-equilibrium realizations of the 5 binding site model when *w/c* = 10 (hollow gray circle in upper right-hand corner of Figure A7B). In this case, however, even the IR vs. 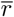 parameter sweeps do not converge reliably, with only 3-4% of sweeps reaching 95% of the global maximum. Thus, we are unable to assess the full extent to which the IR vs *w/c* is sub-optimal in this case. Once again, though, this claims in the main text rely only on the IR bound when *w/c* is large (*w/c* ≳ 10^3^); a regime in which we find that the sweeps perform reliably (hollow gray square in Figure A7B). Thus, we conclude that the IR vs. *w/c* sweeps provide a viable basis for the investigations undertaken in this study.

**Fig. A7.**
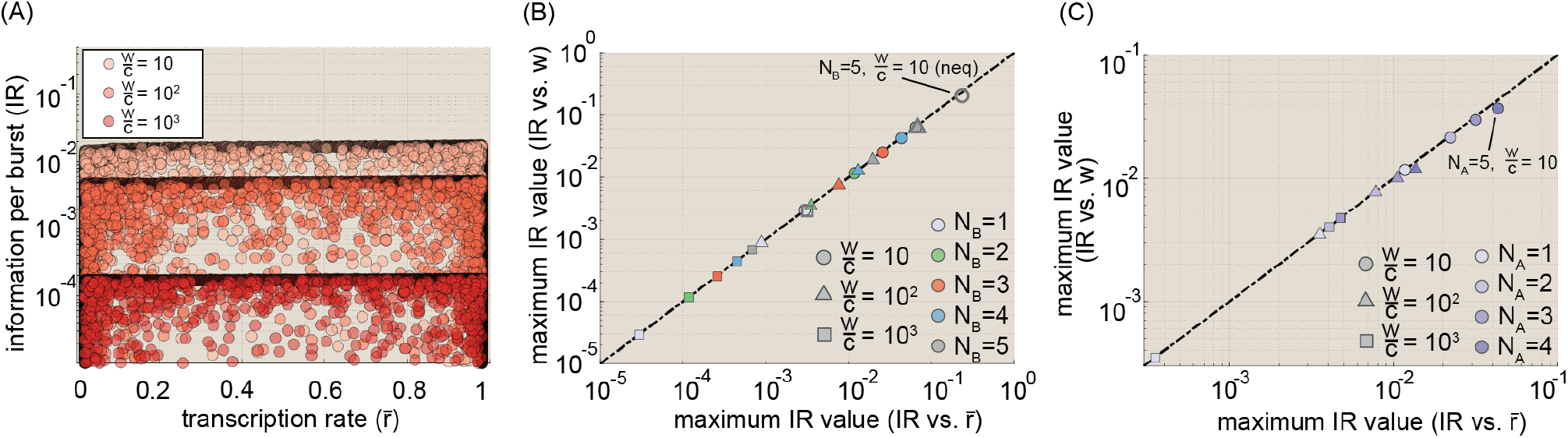
Convergence results for IR vs. *w/c* parameter sweeps. **(A)** Illustrative scatter plot showing IR vs. 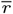 sweep results for the three binding site model at equilibrium for three different values of *w/c*. **(B-C)** Scatter plots comparing parameter sweep results for the upper IR bound at three different *w/c* levels (10, 10^2^, and 10^3^) derived from IR vs. 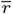 sweeps (x-axis) and IR vs. *w/c* sweeps (y-axis) for equilibrium multi-binding site and non-equilibrium multi-activation step models, respectively. Circles, triangles, and squares indicate *w/c* values of 10, 10^2^, and 10^3^, respectively. Hollow markers in (B) indicate non-equilibrium systems. All other results in (B) are for equilibrium gene circuits (in keeping with Figure 4D). All results in (C) correspond to non-equilibrium gene circuits (in keeping with Figure 4E). (Transition rate and interaction term magnitudes (*k* and *η*) were constrained such that 10^*−*5^ ≤ *kτ*_*b*_ ≤ 10^5^ and 10^*−*5^ ≤ *η* ≤ 10^5^, where *τ*_*b*_ is the burst cycle time. *η*_*ab*_ and *η*_*ib*_ were further constrained such that *η*_*ab*_ ≥ 1 and *η*_*ib*_ ≤ 1, consistent with our assumption that the transcription factor activates the gene locus.)

**Fig. A8.**
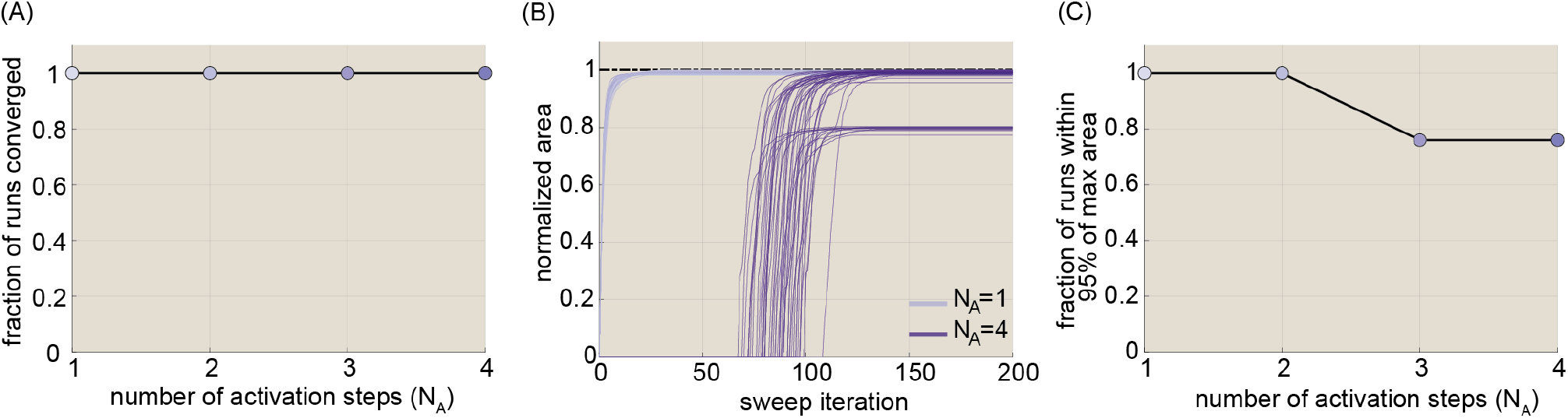
Convergence results for f vs. 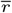 parameter sweeps. **(A)** Plot showing fraction of runs converged as a function of the number of activation steps. All 50 runs converged for each of the four gene circuit models considered. **(B)** Plot showing area spanned in parameter space as a function of iteration number for all 50 runs for the N_A_ = 1 and N_A_ = 4 models. The delayed rise for N_A_ = 4 models reflects the fact that repeated initializations were required to find a sufficient number of gene circuit realizations that adhered magnitude and quality control constraints. **(C)** Fraction of total runs for each model time that reached a final area greater than or equal to the 95% of the global maximum across all runs. (*w/c* was set to 10^3^ for all runs. Transition rate and interaction term magnitudes (*k* and *η*) were constrained such that 10^*−*5^ ≤ *kτ*_*b*_ ≤ 10^5^ and 10^*−*5^ ≤ *η* ≤ 10^5^, where *τ*_*b*_ is the burst cycle time. *η*_*ab*_ and *η*_*ib*_ were further constrained such that *η*_*ab*_ ≥ 1 and *η*_*ib*_ ≤ 1, consistent with our assumption that the transcription factor activates the gene locus.)

#### G.4. Specificity vs. N_A_ results

We claim in the main text that, out of equilibrium, the specificity is bounded by the number of activation steps, such that 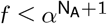. Here *α* is the affinity factor (set to 100) that reflects intrinsic differences in the binding kinetics between cognate and non-cognate factors 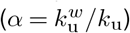. Figure S4B shows parameter sweep results in support of this claim. These results are derived from 2D *f* vs. 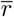 sweeps. Figure A8 shows convergence statistics for these runs for non-equilibrium systems with 1 to 4 activation steps and 1 binding site. We find that all 50 sweep runs met their convergence criteria for each run (Figure A8A) and, further, that no fewer than 76% of runs converged to a 2D area that was within 95% of the global maximum. This indicates that these parameter sweep results converge reliably to consistent overall values for specificity and a function of transcription rate and, thus, that they provide a sound basis for assessing the maximum achievable non-equilibrium specificity as a function of N_A_.

### H. Estimating decision time ranges for different biological systems

#### Caenorhabditis elegans decision time estimation

A recent study by Lee and colleagues (Lee et al., 2019) used live imaging to examine Notch-dependent burst dynamics in the *sygl-1* gene in the germ line of young adult nematodes. Their results indicate that the gene exhibits burst cycle times ranging from 60.5 minutes up to 105.3 minutes (see Figure 2 E and F in (Lee et al., 2019)). Meanwhile, a review article indicated potential values for the cell cycle time for adult germ-line cells in C. *elegans* as ranging from 16 to 24 hours (Hubbard, 2007). A separate study examining nonsense-mediated mRNA decay in C. *elegans* reported a half life of approximately 6 hours for the *rpl-7A* gene (Figure 4k in (Son et al., 2017)). If we take the cell cycle time as the upper time limit for cellular decision-making, this leads to an estimate of 1440*/*60.5 = 23.8 burst cycles.

#### Mus musculus decision time estimation

Burst cycle time estimates were taken from Table A.1 in Appendix A of (Lammers et al., 2020), which indicates times ranging from 30 minutes to a “few hours”. mRNA half life estimates were taken from Table 1 of (Pérez-Ortín et al., 2013), which indicates a range of 30 minutes to 30 hours for mouse cells. To estimate the effective decision time corresponding to an mRNA half-life of 30 hours (1,800 minutes), we recognize that, once mRNA levels have reached a steady state, they will reflect (in effect) a weighted average of the preceding transcriptional activity, where weights moving backward in time contribute

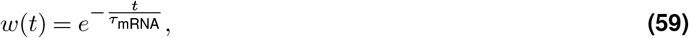

where *t* indicates temporal distance from the present and *τ*_mRNA_ is the exponential time constant, given by 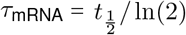. Integrating Equation 59, we find that *τ*_mRNA_ time steps are effectively present in steady-state mRNA levels. Taking 30 minutes as the lower bound for bursting timescales, this yields an upper bound of (1800*/* log 2)*/*30 = 86.6 burst cycles. We note that this estimate is not materially different from the 60 cycle estimate that would be obtained by simply dividing 1,800 by 30.

#### Drosophila melanogaster decision time estimation

We take the duration of nuclear cycle 14, which follows the thirteenth (and final) round of synchronous cellular divisions in early *Drosophila melanogaster* development, as the relevant timescale for cellular decisions in early fruit fly development. Studies have found that the duration of this developmental period varies along the embryo, with a minimum duration of 65 minutes (Foe and Alberts, 1983). To estimate bursting timescales, we use burst inference results from our previous work (Lammers et al., 2020), which indicate a burst cycle time of approximately 2 minutes for the *even-skipped* gene. Thus, we arrive at an upper limit of 65*/*2 = 32.5 cycles.

### I. Higher-order molecular models

Here we provide an overview of key modeling assumptions underlying our approach to modeling gene circuits with multiple activator binding sites or multiple activation steps.

#### I.1. Gene circuits with multiple activator binding sites

A key feature of eukaryotic enhancers is the presence of multiple distinct binding sites for regulatory factors (Vincent et al., 2016; Erokhin et al., 2015). To better understand the impact of variable numbers of binding sites on information transmission, this work examines gene circuit models with between 1 and 5 activator binding sites. In so doing, we maintain the same basic MWC architecture outlined in the context of the simple 4 state model with one activator binding site shown in Figure 1B. No number of bound activators is alone sufficient for mRNA production, but each contributes an extra factor of *η*_ab_ and *η*_ib_ to impact locus activation dynamics. In all cases, we assume a single molecular activation step for multi-binding site models (N_A_ = 1). Finally, we also allow for cooperative interactions between bound activator molecules, which are captured by the interaction term *η*_ub_.

Figure A9A illustrates what this looks like for a model gene circuit with two activator binding sites. The model has eight total states, with four inactive states (top) and four active states (bottom). There are several features to note. First, the transitions between states 2 and 6, which feature two bound activator molecules, are weighted by squaredinteraction terms, 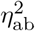 and 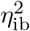, to reflect the regulatory influence of two activators on locus activation dynamics. More generally, if *n* activators are bound, these weights are raised to the nth power (i.e. 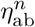 and 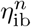). Second, note that unbinding reactions leading out of states 2 and 6 are multiplied by the additional *η*_ub_ mentioned above. This reflects interactions between bound molecules. For simplicity, we assume that *η*_ub_ is the same for both cognate and non-cognate activator species, as well as for interactions between cognate and non-cognate activators. In general, unbinding reactions out of states with *n* activators bound will be weighted by 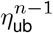 to reflect interactions from the remaining bound factors.

Lastly, a key simplifying assumption that we make in this work is that each activator binding site is identical with respect to its regulatory influence on the gene locus. As a result, it does not matter *which* binding sites are bound, only *how many* are bound. In the context of Figure A9A, this means that states 1 and 5 are functionally identical to states 3 and 7, respectively. Thus, these states can be combined into single coarse-grained states, which leads to an effective model with 6 states, rather than 8. This ability to coarse grain is invaluable for more complex architectures, since it means that the total number of unique molecular states scales as N_A_N_B_, rather than 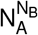.

#### I.2. Gene circuits with multiple molecular activation steps

Setting N_A_ *>* 1 is intended to reflect the reality that multiple distinct molecular reactions—e.g., mediator engagement, PIC assembly, nucleosome displacement, etc.—are necessary preconditions for achieving productive transcription. In the main text we investigate the performance of gene circuits whose transcriptional activity is dictated by 1-4 molecular components, each of which can be either engaged (compatible with transcription) or disengaged (incompatible with transcription). In their simplest interpretation, “engaged” and “disengaged” states might correspond to the presence or absence of some critical component of the transcriptional machinery at the gene locus; however, we remain intentionally non-committal about their physical interpretation, since these generic states are meant to capture a broad swath of potential molecular reactions. For instance, in the case of a nucleosome, the “engaged” state would correspond to the *absence* of the nucleosome (Mirny, 2010). The terms could also capture conformational shifts in key macromolecules such as mediator (Nogales et al., 2017), or in the topology of the gene locus itself.

We assume that each component is required for transcription, such that, in a model with *n* molecular components only molecular states with all *n* components engaged are transcriptionally active, and N_A_ = *n* activation steps are required to achieve locus activation. Furthermore, while in reality each molecular component is likely characterized by heterogeneous dynamics (see, e.g., (Lammers et al., 2020)) we again make the simplifying assumption that each molecular step is identical. As a result, it does not matter which molecular components are engaged, only how many. Figure A9B shows how this logic plays out for the case where N_A_ = 2. As with the N_B_ = 2 case, the model gene circuit has 8 states; however, in this case, only two states (5 and 6)—the ones in which both components are engaged— are transcriptionally active. Note that the binding and unbinding reactions connecting these states are weighted by factors of 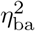 and 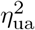, respectively, to reflect the influence of each molecular factor. In general, if *n* components are engaged, these factors are raised to the nth power. In addition, we allow for cooperative interactions between molecular components (curved arrow in states 1,2 and 4-7), captured by the *η*_aa_ and *η*_ia_ terms in Figure A9B. In general these terms are raised to the power of *n* − 1, where *n* is the number of engaged components at the initial molecular states.

#### I.3. Future directions

Throughout this work, we have treated activator binding sites and activation steps as orthogonal axes of gene circuit complexity. In reality, of course, both elements are likely at play in gene regulatory architectures. We choose to investigate the impact of each independently for two chief reasons: first it greatly simplifies exposition and allows us to more easily isolate how each aspect of gene locus architecture interacts with energy dissipation to dictate rates of information transmission. Second, since model complexity scales as N_A_N_B_, we are limited in our ability to accurately explore the performance of models where both N_A_ and N_B_ are large. Improving computational and numerical techniques to permit such explorations represents an interesting future direction. We note also that such models should be tractable without need for additional development if limited to operate at equilibrium.

In addition, we wish to emphasize the potential importance of allowing for heterogeneity, both in the properties of different binding sites along the enhancer and between different molecular components within the activation pathway. This question seems especially interesting in the context of the molecular activation steps. Our simple model with identical steps likely represents the floor of system performance. How much is to be gained when each reaction can adhere to its own kinetics, and exert a distinct kind of regulatory influence over the gene locus?

### J. Deriving normalized sharpness and precision metrics

In Figure 1B, the transcriptional sharpness, *s*, is defined as the first derivative of the transcriptional input-output function multiplied by the activator concentration, *c*^∗^, such that

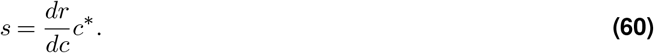

The transcriptional precision, *p*, is defined as the inverse of the intrinsic noise in the transcriptional input-output function:

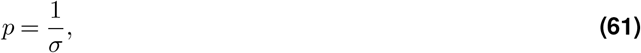

where *σ* is as defined in Equation 11 in the main text. Under these definitions, a key challenge in comparing sharpness and precision levels across different gene circuits is that the upper bounds on both *s* and *p* depend on the fraction of time the system spends in the transcriptionally active state, *π*_a_ (defined in Equation 10). Figure A10A and B illustrate this *π*_a_-dependence for equilibrium and non-equilibrium realizations of the four-state system defined in Figure 1C. As an example: the equilibrium bound on *s* is 0.25 when *π*_a_ = 0.5, but only 0.09 when *π*_a_ = 0.1 (Figure A10A). Since we allow gene circuits to take on different transcription rates (*r* = *π*_a_ *r*_0_) at *C* = *c*^∗^, this *π*_a_-dependence thus confounds our efforts to understand how the molecular architecture of gene circuits—the number of binding sites, number of molecular steps, and presence or absence of energy dissipation—dictates transcriptional performance.

To overcome this issue, we need to normalize *s* and *p* such that they are independent of *π*_a_. Focusing first on sharpness, we were inspired by previous works (Estrada et al., 2016; Grah et al., 2020) to leverage Hill Function as a flexible conceptual tool for extracting generic sharpness measures. The Hill function is defined as:

**Fig. A9.**
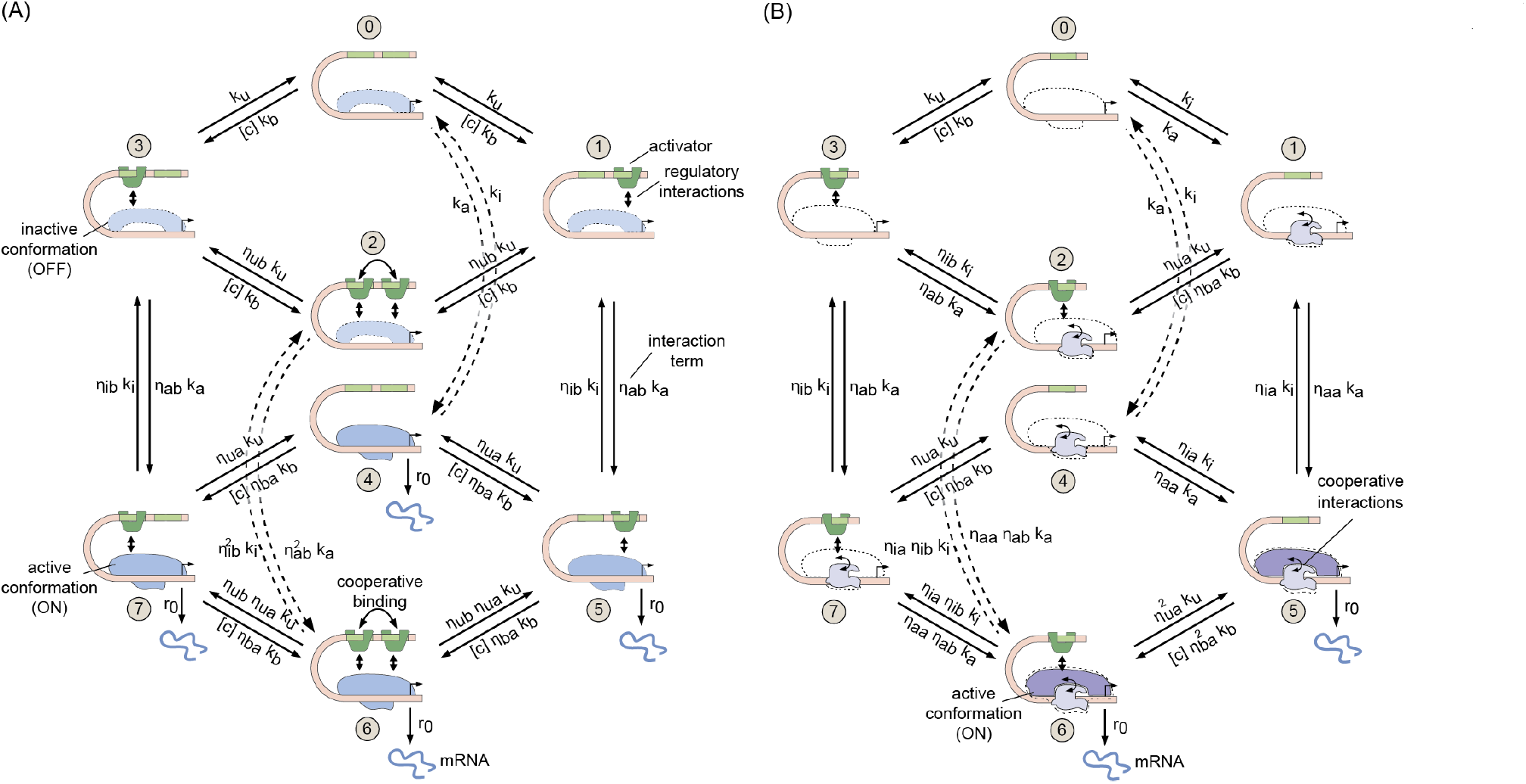
Higher order gene circuit models. **(A)** Cartoon indicating the molecular architecture of a model gene circuit with one activation step and two activator binding sites (N_A_ = 1 and N_B_ = 2). **(B)** Molecular architecture of a model gene circuit with two activation steps and one activator binding site (N_A_ = 1 and N_B_ = 2).

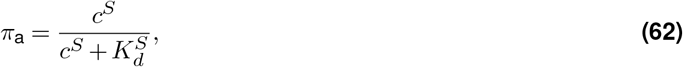

where *c* is the activator concentration, *S* is the Hill coefficient, and *K*_*d*_ is a constant that dictates the location of the function’s half-max point. In general, the input-output functions generated by our model gene circuits will have more complex functional forms, but nonetheless, Equation 62 indicates that we can relate these more complex functions to the Hill function via the shared parameters *π*_a_ and *c*.

The sharpness of the Hill function has the form:

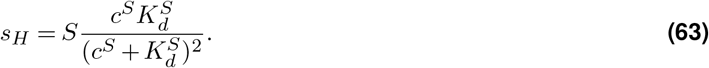

To better relate this to our input-output function, we need to re-express *K*_*d*_ in terms of *C* and *π*_a_. Solving Equation 62 for *K*_*d*_ yields

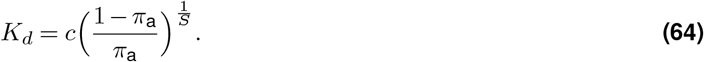

Plugging this in to Equation 63 we obtain, after simplification:

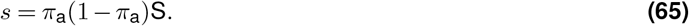

This expression tells us that the sharpness (*s*) of a Hill function with activity level *π*_a_ at *C* = *c*^∗^ is equal to the Hill coefficient, *S*, multiplied by the term *π*_a_(1 − *π*_a_). By rearranging, we can obtain the Hill coefficient as a function of *s* and *π*_a_

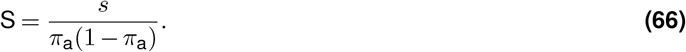

Thus, for a generic gene circuit input-output function with sharpness *s* and expression level *π*_a_ at *C* = *c*^∗^ we can invoke Equation 66 to calculate the Hill coefficient for the equivalently sharp Hill function (Figure A10C). This provides us with a generic measure of transcriptional sharpness that is independent of *π*_a_ and thus can facilitate comparisons across gene circuits that drive differing activity levels at *C* = *c*^∗^ (Figure A10D). We refer to this independent sharpness metric as the “normalized sharpness” in the main text, and denote it with the variable S.

This leads us to the question of transcriptional precision. The two key considerations in defining the normalized precision metric, P, are that (i) we want it to yield a quantity proportional to the information rate when multiplied with S (Equation 66), where

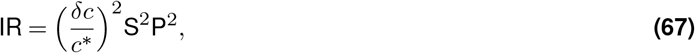

and (ii) we want it to adhere to a single upper bound, regardless of *π*_*a*_. There is only one definition that satisfies the first constraint:

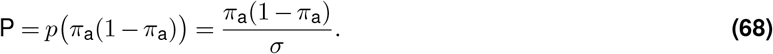

Happily, Equation 68 exhibits consistent upper bounds for all *π*_a_ values, and thus satisfies our second constraint (Figure A10E).

**Fig. A10.**
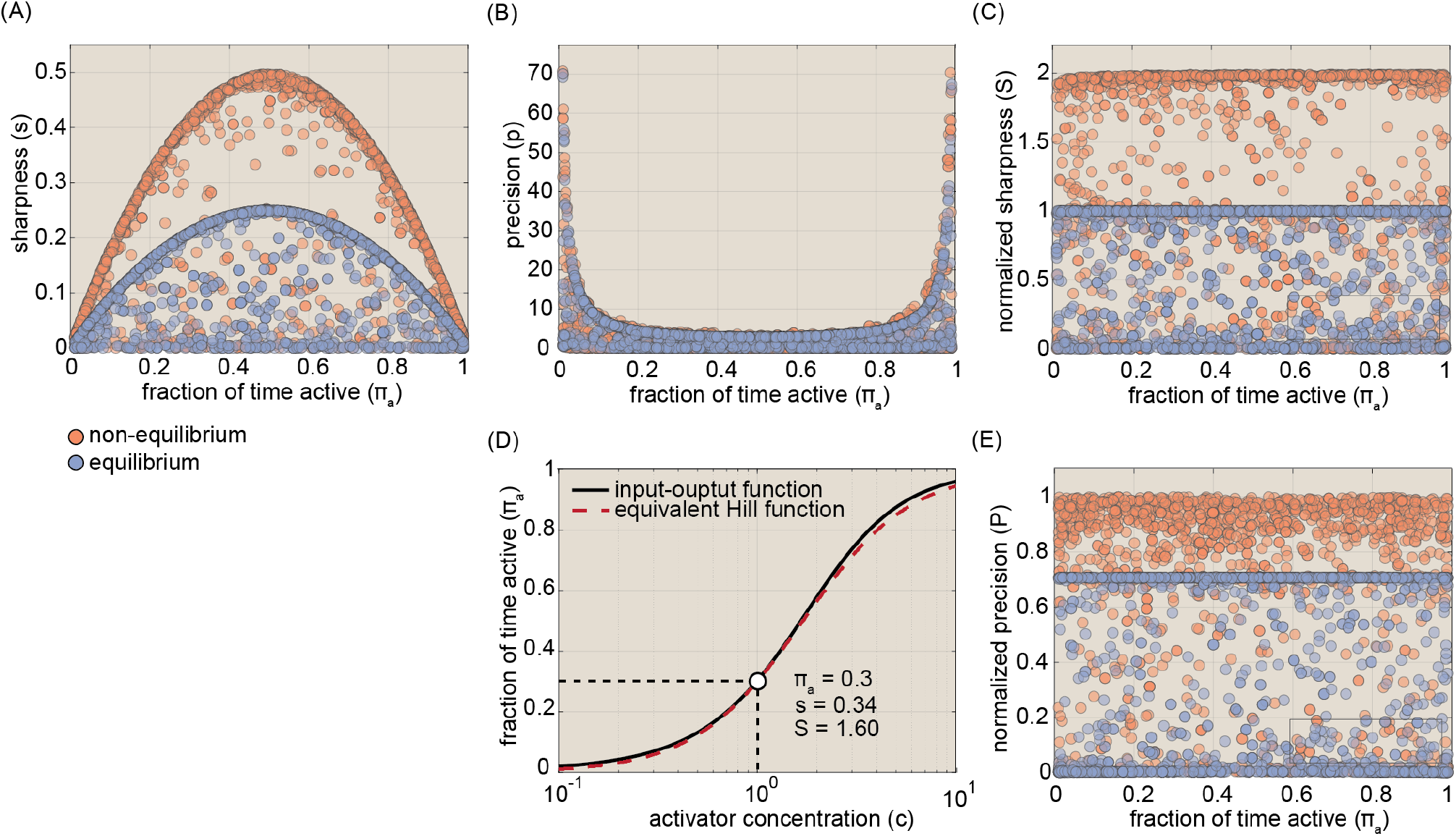
Defining normalized sharpness and precision. **(A)** Plot depicting the upper sharpness limit for equilibrium (blue) and non-equilibrium (red) realizations of the four-state system depicted in Figure 1C. The upper limit depends on the fraction of time spent in the active state, *π*_a_. **(B)** Plot of precision as a function of the transcription rate. Here again, the upper bounds depend on *π*_a_. **(C)** Plot of normalized sharpness as a function of the transcription rate. In the case, the upper limits are invariant. **(D)** Illustration of normalized sharpness concept. For a given input-output curve, we identify normalized sharpness, S, as the Hill coefficient of an equivalently sharp Hill function with the same expression level at *C* = *c*^*∗*^. **(E)** On the other hand, the normalized precision, P, exhibits invariant performance bounds.

### K. Optimal equilibrium four-state gene circuits behave like effective two state systems

In this section, we calculate the normalized sharpness (S) and precision (P) for a simple 2 state gene circuit (Figure A11A) with one ON state and one OFF state and two transition rates, *k*_off_ and *k*_on_. We assume that activator binding dictates fluctuations into and out of the ON state, such that *k*_on_ is proportional to 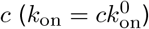. For this simple system, the rate of transcription is given by

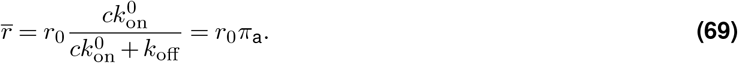

Differentiating this expression with respect to *c* and setting *r*_0_ = 1 (as in main text), we find that

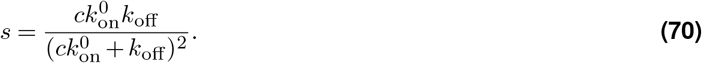

Finally, dividing through by *b* = *π*_a_(1 − _ *π*_a_) yields the normalized sharpness, which is simply given by

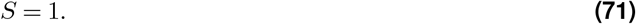

Thus, we see that the two state model is constrained to a normalized sharpness level that represents the upper performance limit for the four-state model operating at equilibrium (blue circles in Figure 3A).

Next, we turn to precision. From Equation 11, we find that

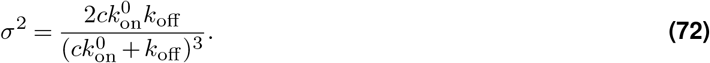

Inverting and multiplying by *b*^2^ gives

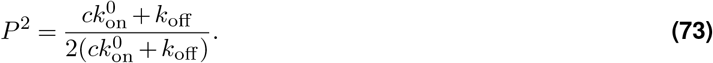

Finally, multiplying through by *τ*_*b*_ (Equation 17) and taking the square root gives

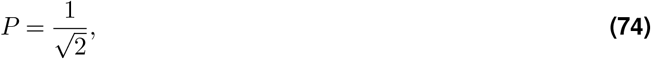

which, again, is equivalent to the upper limit of the four-state gene circuit at equilibrium (Figure 3A).

**Fig. A11.**
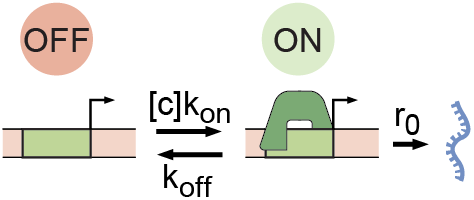
A simple 2 state model of transcription. Cartoon of a simple 2 state gene circuit model in which activator binding and unbinding dictate transitions into and out of a transcriptionally active state.

### L. Sharp and precise non-equilibrium networks exhibit distinct and incompatible microscopic topologies

One simple way to probe the microscopic architectures of different gene circuits is to measure the degree of heterogeneity (or dispersion) in (a) transition rates and (b) state probabilities. We developed entropy-based dispersion metrics ranging from 0 to 1 to quantify how uniform (0) or heterogeneous (1) transition rates and state probabilities were for different realizations of the four-state network shown in Figure 1C. While crude, these measures can provide useful microscopic insights. For instance, in gene circuits with a state probability score of 0 each microscopic state must be equiprobable (*π*_1_ = *π*_2_ = *π*_3_ = *π*_4_ = 1*/*4), while those with a 1 are maximally heterogeneous. In general, maximal heterogeneity corresponds to the case when one and only one state has a nonzero probability; however, since, for simplicity, we have elected here to focus on gene circuits where 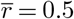, the maximum instead corresponds to a case when two molecular states (one OFF and one ON) have probability *π*_*i*_ = 0.5. Similar considerations hold for the transition rate axis. We conducted parameter sweeps to explore the space of achievable dispersion values for 10,000 non-equilibrium gene circuits (gray circles in Figure A12A).

From Figure A12A, we can see immediately that precise and sharp gene circuits occupy opposite extremes of dispersion space. Specifically, precise systems exhibit highly uniform state probability and transition rate values, while sharp networks are highly heterogeneous, both with respect to the fraction of time spent in each state and the relative magnitudes of their transition rates. These stark differences, as well as the tight clustering of each motif, suggest that sharpness and precision arise from distinct and non-overlapping microscopic topologies.

Detailed examination of maximally precise gene circuits from our parameter sweeps indicated that these systems exhibit highly uniform molecular architectures wherein each microscopic state is equiprobable, all clockwise transition rates are uniform, and all counterclockwise rates are negligible. This results in a “clock-like” system that maximizes the regularity of molecular transitions. Maximally sharp gene circuits, on the other hand, exhibit an all-or-none character, behaving as effective two state systems that spend most of their time either activator-bound and active (0), or unbound and inactive (1), and which have effective ON and OFF rates that are concentration-dependent (see Appendix M for further details).

### M. A hierarchy of microscopic transition rates underpins non-equilibrium sharpness gain

Figure 3A shows that energy dissipation opens up a broad spectrum of S and P values that are not attainable at equilibrium. It is difficult to formulate general statements that apply to all gene circuit models inhabiting these spaces beyond the upper equilibrium limit; however, we can learn much by examining the architecture of gene circuits lying at the outer limits of non-equilibrium performance, since these systems tend to distil the logic underpinning non-equilibrium performance gains into relatively simple regulatory motifs.

Such is the case for the IR-optimized non-equilibrium four-state systems depicted as gray circles in Figure 3A. In Main Text Section D, we found that the driver of this IR is a twofold increase in sharpness relative to the upper equilibrium limit. To realize this twofold sharpness gain, we find that non-equilibrium driving is harnessed to facilitate effective one-way transitions between the active and inactive conformations—specifically, from states 1 to 2 and 3 to 0 in Figure 1C—ensuring that the system will have a strong tendency to complete transcriptional cycles in the clockwise direction (*J >* 0).

In addition to this non-equilibrium driving, sharpness maximization places strict constraints on the relative magnitudes of microscopic transition rates within the network. To understand these constraints, it is instructive to consider a coarse-grained representation of our network with a single ON state (2) and a single OFF state (0). We can obtain expressions for the two effective transition rates in the network by recognizing that they are equal to the inverse of the mean first passage times between states 2 and 0, which we can calculate using Equation 16 from Appendix A.

If we neglect the energetically disfavored transitions from 2 to 1, the effective ON rate (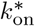 in Figure 3D) takes on a relatively simple form

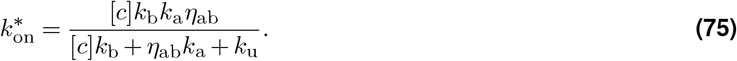

From Equation 75, we see that the effective ON rate becomes proportional to the concentration, *c*, when the factor of [*c*]*k*_b_ becomes negligible in the denominator. The limit where *k*_u_ ≫ [*c*]*k*_b_, *η*_ab_*k*_a_ represents a scenario in which the activator *K*_*d*_ is larger when the network is in the inactive conformation than when in the active conformation such that the activator must bind multiple times (on average) before it succeeds in driving the system into the active conformation. The other limit, when *η*_ab_*k*_a_ ≫ [*c*]*k*_b_, *k*_u_, corresponds to a system where locus activation happens rapidly upon activator binding.

In similar fashion, the effective OFF rate can be expressed as the inverse of the first passage time from 3 to 1

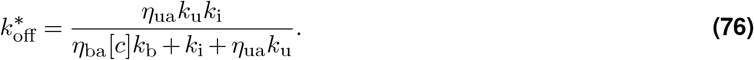

Interestingly, we see that the effective *k*_off_ becomes *inversely* proportional to *c* when activator binding rate exceeds both the unbinding rate and the rate of locus deactivation 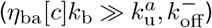. This imbalance causes the system to become kinetically trapped in the active conformation for multiple cycles of activator unbinding and rebinding, with an average duration inversely proportional to *η*_ba_[*c*]*k*_b_.

Thus, when the proper hierarchy of microscopic rates is realized, our four-state network behaves *as though* it were a two state system in which both the on and off rates are concentration dependent, such that

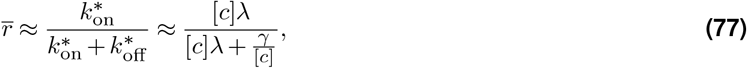

where *λ* and *γ* are coarse-grained transition rates with units of *s*^_1^[*c*]^−1^ and *s*^_1^[*c*], respectively. Repeating the calculations from Appendix K for the above effective two state system will yield an S value of 2 and a P value of 1, in agreement with our numerical results from Figure 3A. We propose that this doubled concentration dependence can be conceptualized as a kind of “on rate-mediated” proofreading. In contrast to classical kinetic proofreading, which works by amplifying intrinsic differences in ligand off rates (Hopfield, 1974; Ninio, 1975), sharp networks amplify the concentration-dependence carried by binding rates, effectively “checking” *C* twice per cycle since both 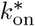 and 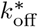 are functions of the activator concentration *c*.

**Fig. A12.**
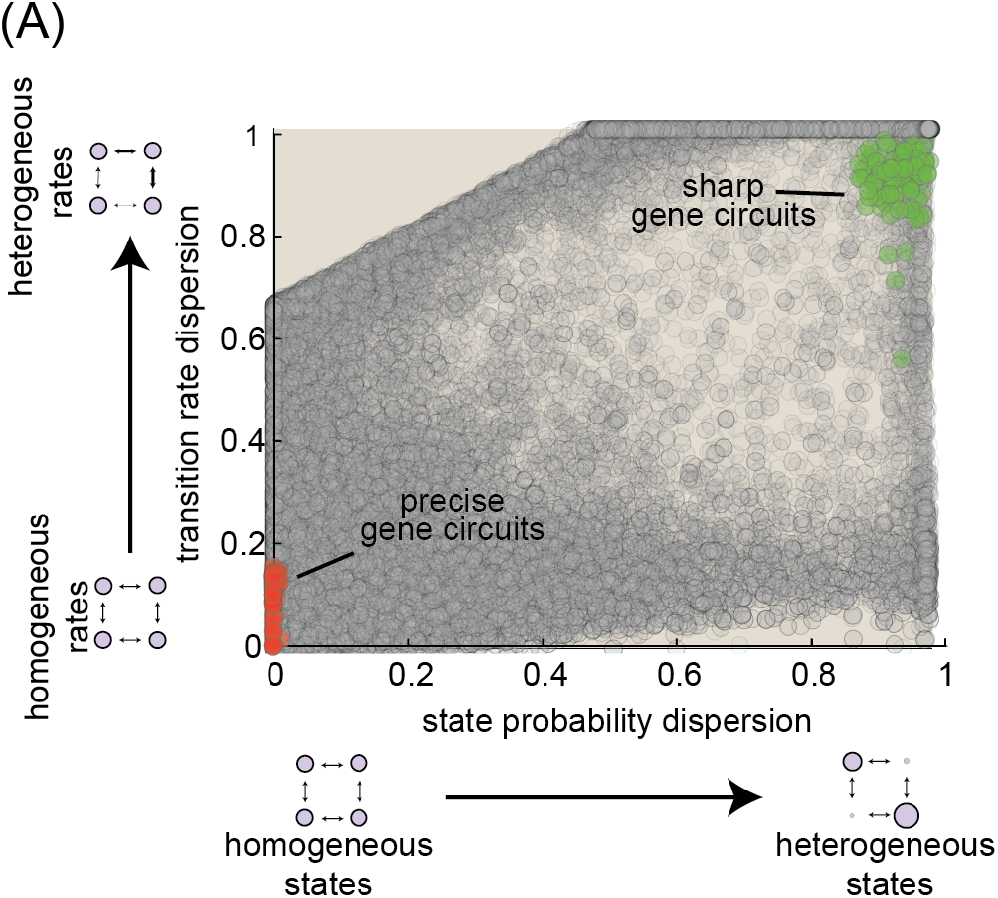
Sharp and precise non-equilibrium networks exhibit distinct and incompatible microscopic topologies. Plot showing dispersion scores for state probabilities and transition rates for 50,000 non-equilibrium networks. Here, a score of 0 indicates maximal uniformity (all rates or probabilities are equal) and a 1 indicates maximal heterogeneity. Green and red circles indicate the scores for the 100 gene circuits within 2% of the maximum achievable non-equilibrium sharpness and precision levels, respectively. (Transition rate and interaction term magnitudes (*k* and *η*) were constrained such that 10^*−*5^ ≤ *kτ*_*b*_ ≤ 10^5^ and 10^*−*5^ ≤ *η* ≤ 10^5^, where *τ*_*b*_ is the burst cycle time. *η*_*ab*_ and *η*_*ib*_ were further constrained such that *η*_*ab*_ ≥ 1 and *η*_*ib*_ ≤ 1, consistent with our assumption that the transcription factor activates the gene locus.)

### N. Non-equilibrium gains in sharpness drive IR increases in more complex regulatory architectures

This appendix section contains additional discussion relating to sharpness-precision tradeoffs for higher-order model architectures with multiple binding sites or multiple activation steps.

#### N.1. Sharpness maximization remains optimal for systems with multiple binding sites

To assess whether sharpness-maximization remains the optimal strategy for more complex architectures featuring multiple activator binding sites, we employed parameters sweeps to examine the space of achievable S and P values for gene circuits with 1-5 activator binding sites (and N_B_ fixed at 1). Figure S2A shows the results of this analysis. For ease of comparison across different models, we plot the relative gains in S and P for each model with respect to their maximum equilibrium values. For instance, the maximum equilibrium S value for the N_B_ = 2 model is 2, so a non-equilibrium gene circuit model with two binding sites that exhibits an S value of 2.5 will be calculated to have a sharpness gain of 2.5*/*2 = 1.25.

Figure S2A reveals that the sharpness-precision tradeoff observed for the one-binding site model persists and, indeed, becomes more severe for systems with additional activator binding sites. We see that the non-equilibrium gain in S is fixed at approximately 2. And while the non-equilibrium gain in P increases from 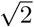 for N_B_ = 1 to approximately 2.25 for N_B_ = 5, these P maxima (peaks in the upper left quadrant of Figure S2A) occur at lower and lower values of S, which renders them more and more disadvantageous from an IR perspective. As a result, when we plot IR-optimal gene circuits for each value of N_B_ (colored circles in Figure S2A), we find that they are invariably located in regions where S*/*S_eq_ ≈ 2 and P*/*P_eq_ ≈ 1. These results demonstrate that spending energy to maximize sharpness remains the key to maximizing transcriptional information transmission, irrespective of the number of activator binding sites.

#### N.2. Multiple activation steps increases upper sharpness bound away from equilibrium

Figure S2B shows the range of achievable non-equilibrium gains in S and P for systems with 1-4 activation steps (and N_B_ = 1). Once again we observe a strong tradeoff between sharpness and precision, which suggests that this incompatibility is a general feature of transcriptional systems. And, once again, we find that IR-maximizing gene circuits (colored circles) lie at or near the right-most edge of achievable parameter space, indicating that dissipating energy to enhance transcriptional sharpness (rather than precision) remains the best strategy for maximizing the IR.

Yet unlike the systems examined in Figure S2A, Figure S2B reveals that the non-equilibrium gain in transcriptional sharpness (S) is not fixed but, rather, increases with the number of molecular steps from a factor of two when N_A_ = 1 to a factor of *five* when N_A_ = 4. This indicates that increasing the number of dissipative molecular steps in the activation pathway raises the upper limit on the sharpness of the transcriptional input-output function, even when the number of binding sites is held constant.

### O. Specificity definitions and details

This Appendix Section uses a simple two state gene circuit model to compare and contrast the specificity definition employed in two recent works (Shelansky and Boeger, 2020; Grah et al., 2020), which compares how a single transcription factor (“TF”) activates at two different gene loci (the “TF-centric” approach)—a target locus with specific binding sites, and a non-cognate locus that lacks binding site—with the definition employed in this work, which focuses on cognate and non-cognate factors competing to activate a single locus (the “gene-centric” approach).

#### O.1. A detailed comparison of specificity definitions for a simple 2-state model of transcription

Figure A13A illustrates the second “Tf-centric” scenario for the case of a simple two state network with a single binding site and no possibility of a conformation change at the locus; however the same idea applies equally well for the 4 state network we considered above, as well as more complicated architectures. Here transcriptional specificity is defined as the ratio of the average steady state transcription rates at on- and off-target gene loci:

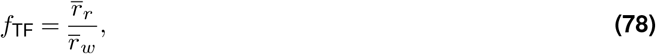

where *f*_TF_ is the specificity under the TF-centric framing of the problem, and *r*_*r*_ and *r*_*w*_ indicate the transcription rates at the cognate (right) and non-cognate (wrong) loci, respectively. In (Shelansky and Boeger, 2020), the authors show that specificity for the two state system shown in Figure A13A is given by:

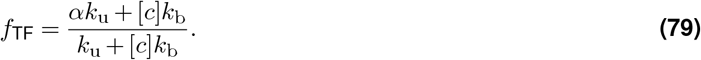

From Equation 79, we see that the activator specificity is bounded from above by *α*. Moreover, this upper performance limit is achieved only in an off rate-dominated regime where *k*_u_ *>>* [*c*]*k*_b_, which the authors in (Shelansky and Boeger, 2020) note leads to a runaway increase in transcriptional noise with increasing specificity under the constraint that the mean transcription rate must remain constant. As a result, the authors conclude that non-equilibrium network architectures are necessary in order to improve specificity and minimize transcriptional noise (Shelansky and Boeger, 2020).

In analogy to the parallel case outlined above, we employ a “gene-centric” definition (Figure A13B), which takes specificity as the ratio of the average number of cognate and non-cognate factors bound while the locus is in a transcriptionally productive state, normalized by concentration:

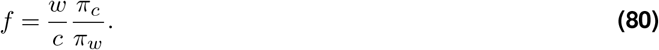

In the case of the two state model shown in Figure A13B, this is simply given by the ratio of fractional occupancies of states 1 and 1^∗^:

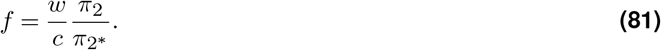

Since in steady state this is necessarily at equilibrium (note the absence of cycles), we can express this ratio as a function of the difference between the energies of cognate and non-cognate factor binding, *ε*_*c*_ and *ε*_*w*_, which leads to

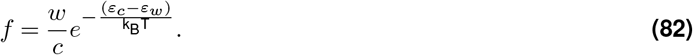

Next, we note that the energies can be expressed as ratios of binding and unbinding rates, such that

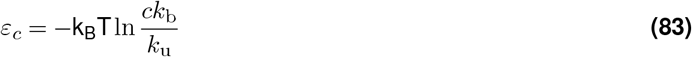

and

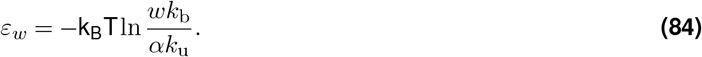

Plugging these two expressions into Equation 82, we have

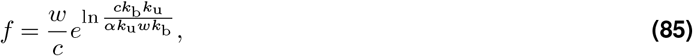

which simplifies to a simple equality

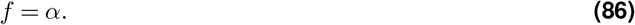

From Equation 86, we see that *f* is simply equal to the binding specificity factor *α* for our three state network, *irrespective of binding kinetics*. Thus, in contrast to (Shelansky and Boeger, 2020), we find that equilibrium gene circuits need not shift towards a noisy, off rate-dominated regime to achieve maximum fidelity; indeed *all* systems necessarily achieve precisely *f* = *α*. Intuitively, this difference stems from the fact that our model captures the effects of kinetic competition between cognate and non-cognate activators: whenever the cognate activator (green square in Figure A13B) is bound, non-cognate factors cannot bind.

A key limitation of this approach is that it neglects the presence of non-specific stretches of regulatory DNA, even at cognate gene enhancers. Thus, to more accurately reflect the specificity challenges faced by real gene loci, a synthesis of the two approaches summarized above will be necessary, which considers competition between cognate and non-cognate factors to bind and activate a gene locus that features both specific binding sites (which favor the cognate activator) and neutral sites (to which all activator species bind non-specifically). One expectation for such a scenario is that the simple equality stated in Equation 86 will no longer hold, and tradeoffs similar to those observed in (Shelansky and Boeger, 2020) will again emerge; although, this time, the severity of these tradeoffs will depend on *w/c*.

#### O.2. Calculating equilibrium specificity for a gene circuit with one binding site and one activation step

Here we extend the arguments from the previous section to show that, at equilibrium, the transcriptional specificity of the six state model gene circuit shown in Figure 4B is fixed at *f*^*eq*^ = *α*, irrespective of molecular details. For this system, the specificity is simply equal to the concentration-normalized ratio of the occupancies of states 2 and 4:

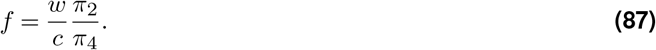

Since we’re assuming equilibrium conditions, we can re-express this as a difference between state energies, such that

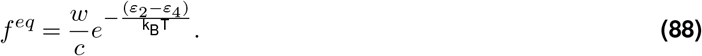

**Fig. A13.**
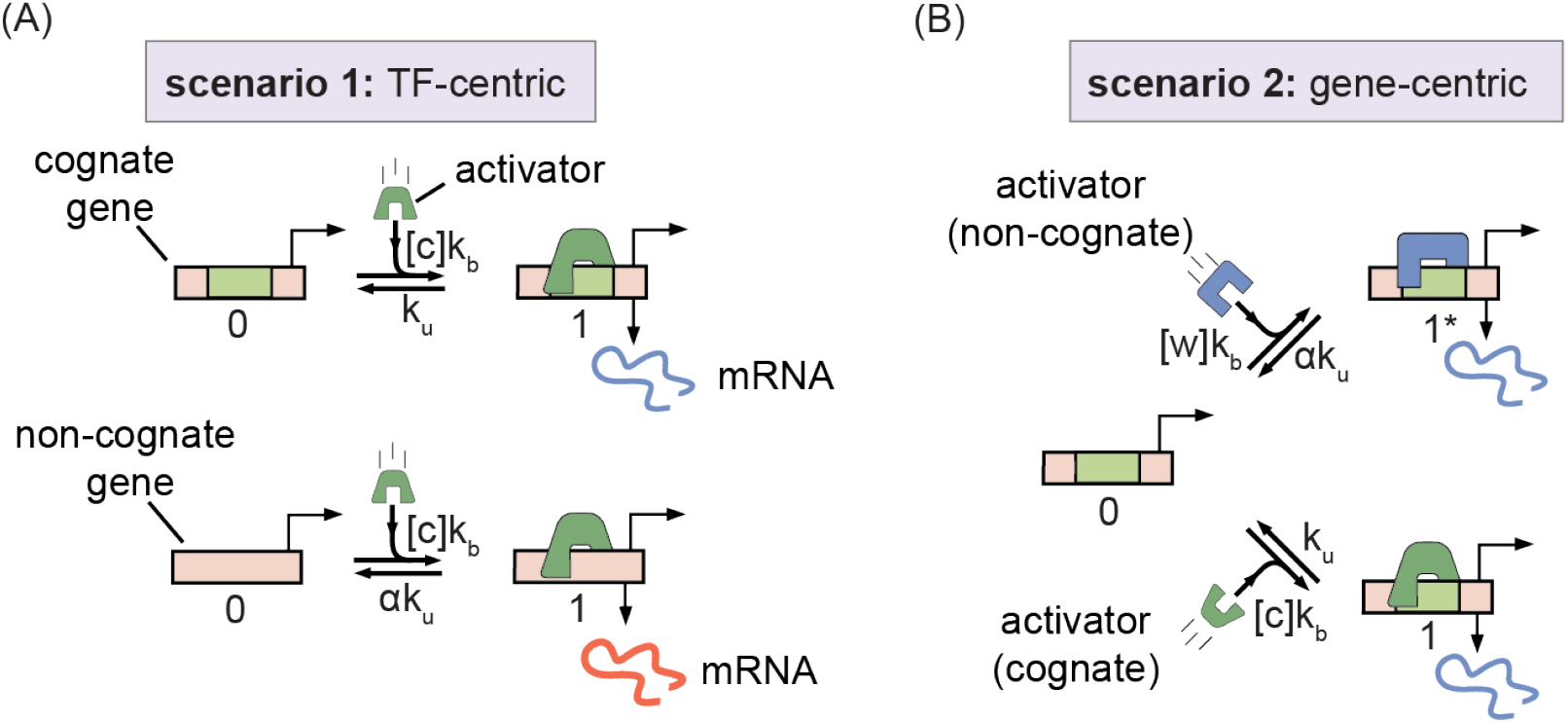
Accounting for the influence of off-target activation. **(A)** An illustration of the parallel definition of activation fidelity. This approach considers the relative amounts of transcription driven by a transcriptional activator at its target locus and at an off target locus. **(B)** Cartoon illustrating “gene-centric” specificity definition, which considers competition between cognate and non-cognate factors to bind and activate a single gene locus.

In each case, we can express the state energies as the sum of the energy due to cognate or non-cognate factor binding with the energetic contributions from being in the active conformation, *ε*_a_, and from interactions between the activator and the locus conformation, *ε*_ab_. This leads to

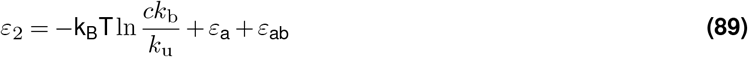

and

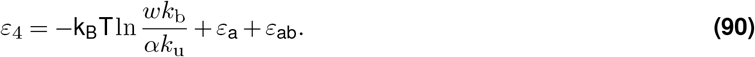

The key is to note the the first terms on the right-hand side of the above expressions are identical to Equations 83 and 84. Since the remaining energy terms are identical, they will cancel out, such that we once again have

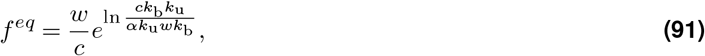

which simplifies to

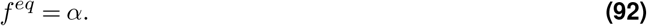

#### O.3. Calculating equilibrium specificity for gene circuits with multiple binding sites

The above arguments can be extended to apply to more complex model architectures with multiple activator binding sites. To do this, we first need to generalize the definition of specificity put forward in the main text (Equation 5) for the case when there is more than one binding site. We define multi-site specificity as the ratio of the number of cognate and non-cognate activators bound to the gene locus (on average) while the gene is in the transcriptionally active (ON) conformation, such that:

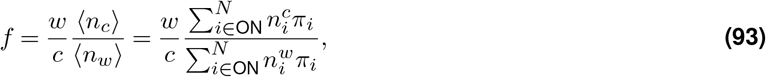

where *i* ∈ *ON* stipulates that state *i* is part of the ON conformation, *π*_*i*_ is the probability of finding the gene locus in state *i*, and where 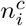 and 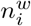 indicate the number of cognate and non-cognate factors bound to the gene locus in state *i*. Note also that we retain the normalizing prefactor of *w/c*.

Now, let’s calculate *f* for an equilibrium gene circuit with two binding sites. Once again, we work with energies since the system is at equilibrium. From Equation 93 we see that only states with at least one cognate or non-cognate factor bound contribute to the numerator and denominator, respectively. As a result, in each case, there are just three distinct molecular states to consider. For the cognate case (numerator), these are 1 cognate bound and 0 non-cognate, 1 cognate and 1 non-cognate, and 2 cognate. The non-cognate case (denominator) follows the same pattern. Drawing from the expression in the previous section, this leads to

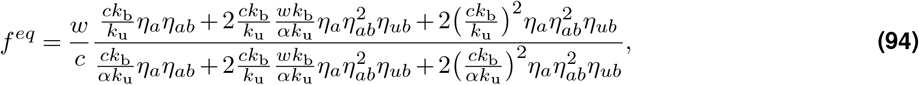

Where *η*_*a*_ is a weight factor corresponding to the active conformation 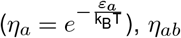 is a weight factor capturing cooperative interactions between the bound activator and the active conformation, and *η*_*ub*_ captures cooperative interactions between bound activator molecules. Note that the three terms in the numerator and denominator of Equation 94 match the ordering of the scenarios given above the equation. Factoring out common multipliers leads to

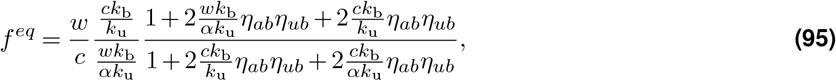

where we now see that the numerator and denominator are identical in the right-most ratio. Thus, we find that

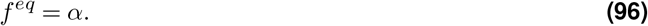

Similar patterns repeat for systems with more binding. See the Mathematica notebook entitled “specificity_ multi_site.nb” in this paper’s git repository (https://github.com/nlammers371/noneq-gene-regulation.git) for a full treatment of the 3 and 5 binding site cases.

### P. Deriving non-equilibrium tradeoff bound between intrinsic sharpness and specificity

In this section, we lay out the key steps in deriving the non-equilibrium tradeoff bound between sharpness and specificity given in Equation 7 in the main text. To do so, we make use of insights gained in Appendix M, where we used first passage times to examine the key microscopic conditions for the twofold gain in sharpness away from equilibrium observed in Figure 3A. Even for the simple six state system illustrated in Figure 4B, our system has eight degrees of freedom when operating away from equilibrium. As such, a key part of our approach will be to first reduce this complexity as much as possible while preserving the salient behaviors, namely the possibility for non-equilibrium gains in sharpness and specificity. After this, we identify a tuning parameter, *β*, that can be used to interpolate between maximally sharp to maximally specific non-equilibrium gene circuit architectures. Since the expressions for non-equilibrium gene circuits are, in general, quite complex, we sketch the key steps here and direct the reader to the Mathematica notebook entitled “sharpness_specificity_bound_derivation.nb” on the project git repository for additional details: https://github.com/nlammers371/noneq-gene-regulation.git. Note that we work in units of *c* throughout, such that *c* = 1.

To begin, we strip unnecessary dimensions from our system. We set *η*_ib_*k*_i_ and *k*_a_ to the same generic rate, *k*_1_. Next, we set *η*_ab_*k*_a_, *η*_ua_*k*_u_, and *k*_b_ to a second rate parameter, *k*_2_. Finally, we set *k*_i_ equal to *βη*_bs_*k*_b_, where *β* is our interpolation parameter. This leaves us with a system with five free parameters, rather than eight.

In Appendix M, we saw that maximally sharp non-equilibrium gene circuits (i) only switch into the active transcriptional conformation when the activator is bound and (ii) only switch *out* of the ON states when the activator is unbound. This amounts to effective one-way transitions from states 1 → 2 (equivalently, 5 → 6) and 3 → 1. We impose this condition by taking the limit where *k*_1_ → 0. Next, we impose the condition uncovered by examination of Equation 75,

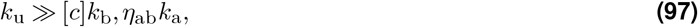

by taking the limit where *k*_u_ approaches infinity.

These limits lead to a further simplified system that can be used to investigate fundamental tradeoffs between intrinsic sharpness and specificity. For this stripped-down system, we find that the expression for specificity, *f*, is quite simple:

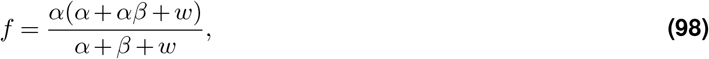

where we see that all dependence on microscopic transition rates has dropped out, with the exception of our interpolation parameter, *β*. Furthermore, tuning *β* causes Equation 98 to shift from equilibrium levels (*f* = *α* when *β* = 0) to the non-equilibrium limits revealed by Figure 5B (*f* = *α*^2^ when *β* ≫ *α, w*).

The normalized sharpness, S, has a slightly more complicated functional form, given by

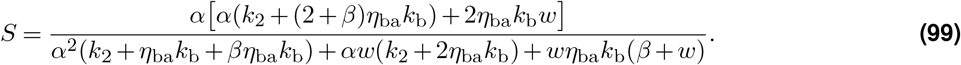

To obtain an expression for the intrinsic sharpness, *S*_0_, we divide through by the specificity prefactor (*p*_*c*_) from Equation 4:

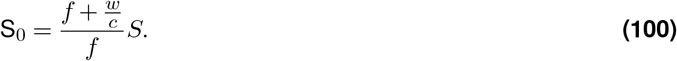

Simplifying and applying the condition that *k*_2_ ≈ 0 leads to

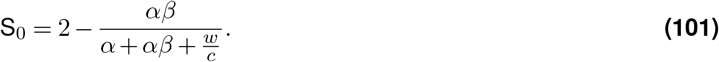

Here again, as with Equation 98, we see that all dependence on the on rate parameters drops away. Further, it is easy to see that this expression goes to 2 when *β* = 0 and 1 when *β* ≫ *α, w*. Thus, when *β* is small, our system exhibits equilibrium levels of specificity and non-equilibrium levels of intrinsic sharpness and, when *β* is large, it exhibits non-equilibrium specificity and equilibrium sharpness levels. Thus, we have succeeded in our initial aim to establish a simplified model that can capture the tradeoffs between sharpness and specificity revealed by our numerical parameter sweeps (Figure 5B).

As a final step, we can solve Equation 98 to obtain an expression for *β* in terms of *f* :

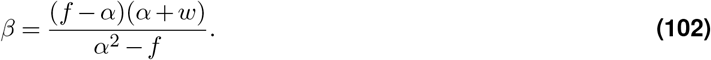

Plugging this expression into Equation 100 and simplifying yields an expression for *S*_0_ as a function of *f* :

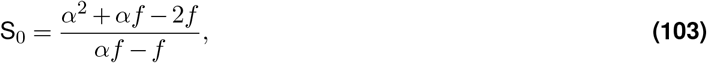

where we assume that *α* ≤ *f* ≤ *α*^2^. Thus, we have obtained the final *S*_0_ expression depicted in Equation 7. Observe that S_0_ ≈ 2 when *f* = *α* and S_0_ ≈ 1 when *f* = *α*^2^. Equation 103 gives the dashed black curve bounding *f* vs. S_0_ sweep results shown in Figure 5B, confirming that it represents the limiting behavior of intrinsic sharpness and specificity for non-equilibrium realizations of the six state model from Figure 4B.

